# Growth substrate limitation enhances anaerobic arsenic methylation by *Paraclostridium bifermentans* strain EML

**DOI:** 10.1101/2023.01.09.523296

**Authors:** Jiangtao Qiao, Hugo Sallet, Karin Lederballe Meibom, Rizlan Bernier-Latmani

## Abstract

Microbial arsenic methylation is established as a detoxification process under aerobic conditions (converting arsenite to monomethylated arsenate) but proposed to be a microbial warfare strategy under anoxic conditions due to the toxicity of its main product monomethylarsonous acid (MMAs(III)). Here, we leveraged a paddy soil-derived anaerobic arsenic methylator, *Paraclostridium bifermentans* strain EML, to gain insights into this process. Strain EML was inoculated into a series of media involving systematic dilutions of Reinforced Clostridial Broth (RCB) with 25 μM arsenite to assess the impact of growth substrate concentration on arsenic methylation. Growth curves evidenced the sensitivity of strain EML to arsenite, and As speciation analysis revealed the production of MMAs(III). Concentrations of MMAs(III) and arsenic methylation gene (*arsM*) transcription were found to be positively correlated with the RCB dilution, suggesting that substrate limitation enhances *arsM* gene expression and associated anaerobic arsenic methylation. We propose that growth substrate competition between microorganisms may also lead to an increase in anaerobic As methylation. This hypothesis was further evaluated in an anaerobic co-couture mode of strain EML with either wild-type *Escherichia coli* K-12 MG1655 (WT) or *E. coli* expressing the MMAs(III)-resistance gene (*arsP*), (ArsP *E. coli*). We found increased MMAs(III) production in the presence of *E. coli* than its absence and growth inhibition of WT *E. coli* to a greater extent than ArsP *E. coli*, presumably due to MMAs(III) produced by strain EML. Taken together, our findings point to an ecological role for anaerobic arsenic methylation, highlighting the role of microbe-microbe competition/interaction in this process.

**IMPORTANCE:** Anaerobic arsenic methylation is enhanced in rice paddy soils under flooding conditions than that under drying conditions, leading to increased methylated arsenic accumulation in rice grains. Unlike the known detoxification role for aerobic arsenic methylation, the ecological role of anaerobic arsenic methylation remains elusive and is proposed to be an antibiotic-producing process involving in microbial warfare. In this study, we interrogated a rice paddy soil-derived anaerobic arsenic-methylating bacterium (*Paraclostridium bifermentans* strain EML) to investigate the effect of growth substrate limitation on arsenic methylation by strain EML in the context of the microbial warfare hypothesis. We provide direct evidence for the role of growth substrate competition in anaerobic arsenic methylation by strain EML. Furthermore, we evidence a feedback loop, by which a bacterium resistant to MMAs(III) enhances its production, presumably through enhanced *arsM* expression resulting from substrate limitation. Our work uncovers complex interactions between an anaerobic arsenic methylator and potential competitors.

## INTRODUCTION

Microbial transformations play an important role in the biogeochemical cycling of arsenic (As) in the environment, and include reduction, oxidation, thiolation, methylation, and demethylation of inorganic and organic As (1–4). These reactions impact the mobility, bioavailability, and toxicity of As compounds (1–3, 5, 6). In recent years, microbial transformations of As in paddy soil have drawn increasing attention because of the potential health risk of dietary exposure of As from rice-containing products (7, 8). For instance, organic As (in particular dimethyl arsenate, DMAs(V)) is commonly detected in rice grains, along with inorganic As (9, 10). As DMAs(V) is much less toxic than arsenite, accumulation of DMAs(V) in rice grains largely reduces its toxicity to humans. However, there is evidence of a correlation between DMAs(V) accumulation in rice grains and rice straight-head disease, a condition that decreases rice crop yields (11, 12). In addition, because analyzing As in rice requries digestion, there is a concern that the highly toxic dimethylated monothioarsenate (DMMTA) may be transformed to and measured as DMAs(V) (13), unknowingly overlooking a potential threat to human health.

Arsenic methylation is a microbially-mediated process involving the transformation of inorganic trivalent As (iAs(III)) into mono-, di-, and trimethylated As compounds and is catalyzed by *S*-adenosyl-methionine methyltransferase (ArsM in prokaryotes) (14–16). Generally, As methylation occurring under oxic conditions is proposed as an iAs(III)-detoxifying process because although more toxic As compounds (monomethylarsonous acid (MMAs(III) and dimethylarsinous acid (DMAs(III)) are produced, they are rapidly oxidized in the presence of O_2_ to their less toxic pentavalent counterparts (monomethylarsonic acid (MMAs(V) and dimethylarsinic acid (DMAs(V)) (16). This paradigm is supported by the fact that the heterologous expression of the *arsM* gene conferred As(III) resistance to an As(III)-sensitive *Escherichia coli* strain under aerobic conditions (16, 17).

In contrast, detoxification is unlikely to be the ecological function of As-methylators inhabiting anoxic environments since organic As products are present in their trivalent forms. Interestingly, the evolutionary history of the *arsM* gene predicts its emergence during the anoxic Archaean era, when MMAs(III) would have been chemically stable (18, 19). This finding suggests that the original function of MMAs(III) could have been to serve as a primitive antibiotic (20, 21). A closer examination of As methylation capability among previously reported and available pure anaerobic strains uncovered either the negligible methylation efficiency or the fortuitous methylation of As upon cell lysis and the release of methyltransferases (22), making it challenging to decipher the physiological function of anaerobic As methylation. At present, confirmed and available anaerobic As-methylating bacteria come from the genus *Paraclostridum* (23, 24). Here, we will study *Paraclostridium bifermentans* strain EML (henceforth strain EML), a fermenter isolated from a paddy soil in Vietnam (24).

In microbial communities, competitive phenotypes (including the production of antibiotics) can arise as a consequence of limited resources (e.g., nutrients and space) (25). Soils represent an ecosystem in which many microorganisms compete for scarce resources, thus competition is widespread (26). We hypothesize that competition for resources may boost As methylation by strain EML, conferring it an advantage over its competitors. In this study, we aim to investigate the effect of growth substrate limitation on As methylation by strain EML in the context of the microbial warfare hypothesis for anaerobic As methylation. Further, we probe direct microbial interaction/inhibition between strain EML and potential competitors (*Escherichia coli* MG1655 either the wild-type strain (WT) or one engineered to express the MMAs(III)-resistance gene (*arsP*)). From an ecological point of view, it is reasonable to expect enhanced As methylation under substrate-limiting conditions as a response to resource competition. According to our hypothesis, strain EML would increase the production of toxic MMAs(III) under growth substrate-limited conditions in order to thwart other microorganisms competing for the same resources. Although DMAs(III) might also function as an antibiotic under anaerobic condition, we focus on MMAs(III) in this study due to the analytical limitations associated with DMAs(III).

## RESULTS

### iAs(III) inhibits strain EML growth

Strain EML was grown under anoxic conditions with various concentrations of RCB (100%, 75%, 50%, or 25% RCB) in the presence and absence of 25 μM iAs(III). The growth curves show that strain EML grew rapidly and reached the mid-exponential phase after about 8 hours in the absence of iAs(III) (Fig. 1). In contrast, the extent of growth of strain EML was lower in the presence of iAs(III) and it reached stationary phase sooner than in the absence of iAs(III), particularly in low growth-substrate media (50% and 25% RCB) (Fig. 1). While strain EML harbors a gene encoding an iAs(III) efflux permease (*acr3*) (Fig. S7), we hypothesize that it may not pump out intracellular iAs(III) sufficiently fast to preclude toxicity.

**FIG 1.**
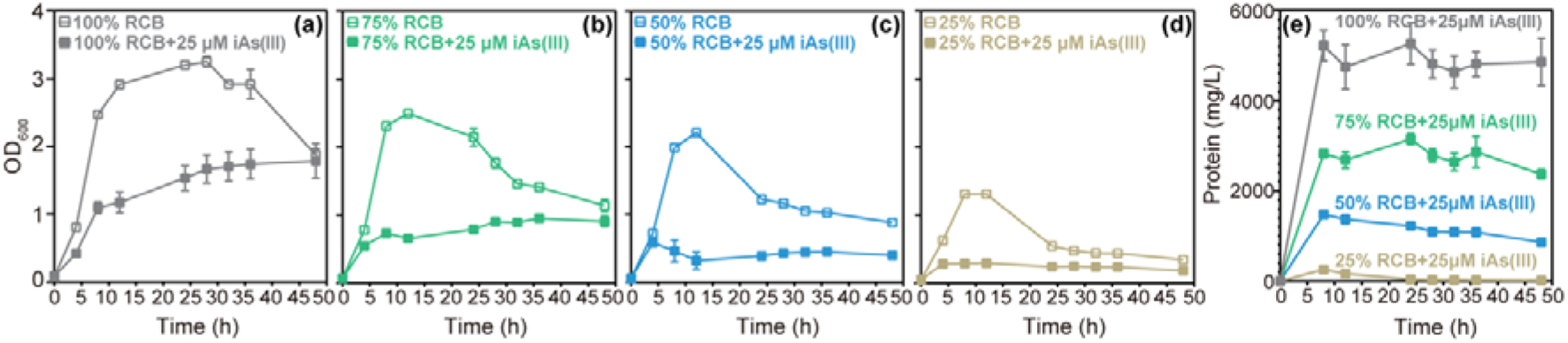
Growth curves (OD_600_ (a-d) and total protein (e)) of *Paraclostridium bifermentans* strain EML in anaerobic dilutions of Reinforced Clostridial Broth (RCB) (100%, 75%, 50%, or 25% RCB) in the presence or absence of 25 μM iAs(III). Data are shown as mean values with error bars. Individual values for each biological triplicate are included in Data Table 1.

### Impact of substrate concentration on MMAs(III) production by strain EML

To profile the dynamics of iAs(III) transformation during anaerobic growth of strain EML, time-dependent changes in As speciation in solution (aqueous) and inside cells (soluble intracellular) were monitored. Analysis of aqueous As species clearly shows that MMAs(III) was gradually produced by strain EML and increased during the exponential growth phase (0-12 hours) and reached a plateau between 24 and 48 hours (post-stationary to death phase) (Fig. S8a-d). Sterile RCB amended with 25 μM iAs(III) exhibited no transformation of iAs(III) (approximately 25 μM iAs(III) was detected at the beginning and end of incubation) (Table S3).

As expected, strain EML exhibited variable growth rates for varying RCB dilutions (Fig. 1), confounding the interpretation of whether growth substrate levels affected the extent of As methylation. Normalization of methylated As (with and without oxidation) to protein concentration (Fig. 2) reveals a trend in normalized MMAs(III) (Fig. 2a) or MMAs(V)/DMAs(V) concentration as a function of RCB dilution (Fig. 2b,c). Indeed, the normalized MMAs(III) concentration decreased in the following order: 25% RCB > 50% RCB > 75% RCB > 100% RCB (Fig. 2). The greatest amount of protein-normalized MMAs(III) was produced by strain EML grown in the highest RCB dilution (25% RCB, 7,766 ± 919 nmol/g protein), which was about 14 (546 ± 39 nmol/g protein), 34 (224 ± 22 nmol/g protein), and 51 (150 ± 21 nmol/g protein) times higher than that in the 50%, 75%, and 100% RCB conditions, respectively (Fig. 2a). Similar patterns were also observed for the oxidized samples, with the highest protein-normalized concentrations of MMAs(V) and DMAs(V) generated in the highest RCB dilution (25% RCB) (Fig. 2b,c). Analysis of soluble intracellular As also clearly shows that strain EML accumulates high amounts of intracellular As(III) and MMAs(V) during anaerobic As methylation, particularly in the highest RCB dilution (25% RCB) (Fig. S9).

**FIG 2.**
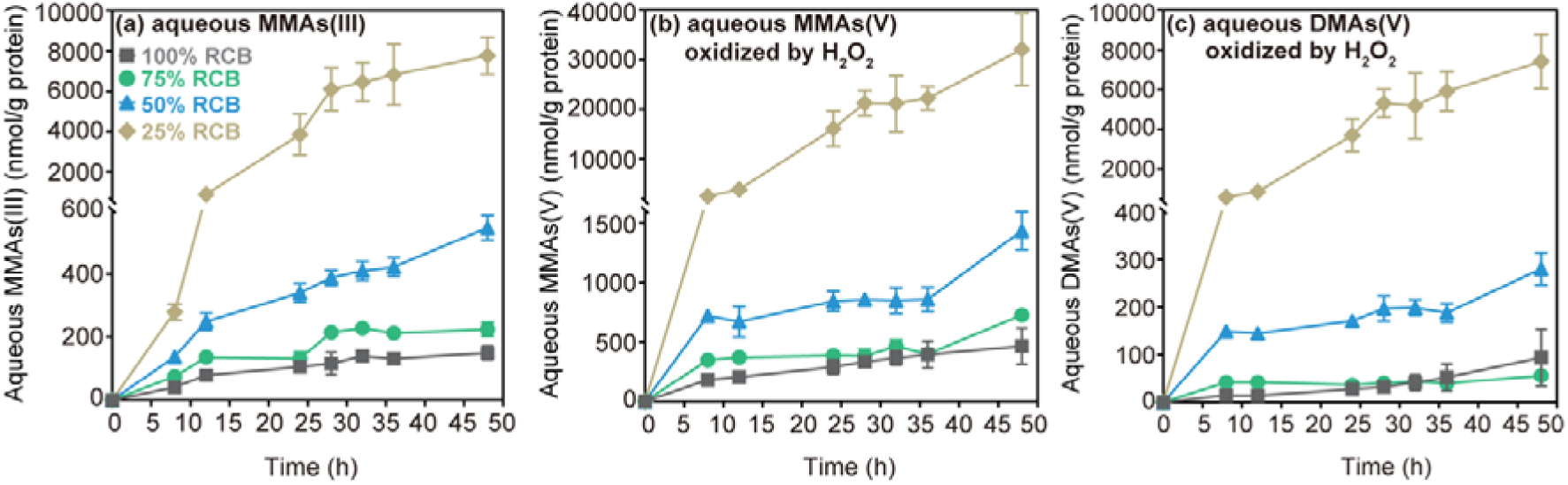
Time-dependent concentrations of protein-normalized aqueous As species in anaerobic RCB dilutions (100%, 75%, 50%, or 25% RCB) inoculated with *Paraclostridium bifermentans* strain EML and 25 μM iAs(III). (a) MMAs(III) (no oxidation), (b) MMAs(V) (post-oxidation) and (c) DMAs(V) (post-oxidation). Individual values for each biological replicate can be found in Data Tables 2 and 19.

### Chemical transformation of MMAs(III) in biological media

While MMAs(III) is clearly produced by strain EML, its chemical stability is often limited, even under anoxic conditions, due to side chemical reactions such as thiolation, resulting in an underestimation of the concentration of MMAs(III) produced. We tested this stability by amending anoxic RCB (100%-25%) or anoxic spent RCB (in which strain EML had grown) with 3 μM MMAs(III) and documented its significant disappearance from solution (after 24 h). Surprisingly, MMAs(III) stability was greatest in 100% RCB and lowest in 25% RCB, suggesting a negative correlation between MMAs(III) stability and medium dilution (Fig. S10 and S11). The limited chemical stability of MMAs(III) in RCB medium (fresh or spent) suggests that MMAs(III) production is being underestimated by our measurements in all conditions but more so in the more dilute RCB medium.

The disappearance of MMAs(III) from solution upon its amendment to RCB medium was puzzling and we hypothesized the formation of methylated-thiolated As species (e.g., monomethyldithioarsenate, MMDTAs(V)), some of which are not identifiable analytically in our system. To probe this possibility, we analyzed As speciation after oxidation of trivalent As species by 10% (v/v) H_2_O_2_ (27–29). Oxidation is expected to transform MMAs(III) into MMAs(V) quantitatively and to oxidize the thiol group in MMDTAs (and other monomethylated-thiolated species) to sulfate, which is released, leaving MMAs(V) as the final product. Indeed, after oxidation, the concentration of MMAs(V) was greater than that of MMAs(III) measured prior to oxidation (Fig. 2 and S8e-h; Text SR4), suggesting the oxidation of monomethylated As compounds other than MMAs(III) to MMAs(V).

Direct evidence of MMAs(III) chemical transformation was provided by the reaction of sulfide with MMAs(III) and subsequent retention of part of the As species by the column (Fig. S12a). Furthermore, following oxidation with H_2_O_2_, the entire As inventory is recovered as MMAs(V) (Fig. S12b). Therefore, we proposed that compounds (such as MMDTAs(V) or others) are formed via the chemical reaction of MMAs(III) with reduced sulfur compounds in the growth medium. These reduced chemical species are likely retained by the HPLC column. If the samples are oxidized prior to measurement, the monomethylated-thiolated species are oxidized to MMAs(V), which is readily eluted. Thus, in oxidized samples, MMAs(V) corresponds to the sum of MMAs(III) and monomethylated-thiolated As species.

### *arsM* gene transcription under variable substrate conditions

In order to investigate whether the transcription of the gene responsible for As(III) methylation (*arsM*) responded to growth substrate concentration, gene expression was quantified using RT-qPCR. We first attempted to use relative expression analysis and evaluated the expression stability of 8 potential reference genes (Table S2) with the qBase plus software. Unfortunately, no optimal number of reference genes could be found due to the relatively high variability among sequential normalization factors (geNorm V > 0.15) and the limited (medium) expression stability was achieved (0.5 < average geNorm M ≤ 1.0) (Fig. S13). We presume that the considerable variation in growth rate due to RCB dilutions (and presence of iAs(III)) markedly impacted gene expression, even for so-called housekeeping genes. Therefore, we turned to absolute quantification. We adjusted the biomass (OD_600_) of strain EML obtained through variable RCB dilutions to approximately the same value prior to RNA extraction to eliminate possible biomass-related biases in RNA extraction and reverse transcription. As expected, the transcripts of strain EML *arsM* gene (adjusted to OD_600_) were significantly higher (*p* < 0.05) in the presence of iAs(III) compared to no iAs(III) controls (Fig. 3a). Among the treatments with iAs(III), we observed that *arsM* gene transcript copy numbers exhibited an opposing trend to substrate content: 25% RCB > 50% RCB > 75% RCB > 100% RCB (Fig. 3a). The highest number of *arsM* transcripts was detected in the most dilute medium (25% RCB + iAs(III), 1.04E+05 ± 2.05E+04 copies/OD_600_), which was approximately 4, 5, and 12 times greater than in the treatments of 50% RCB + iAs(III) (2.63E+04 copies/OD_600_), 75% RCB + iAs(III) (2.11E+04 copies/OD_600_), and 100% RCB + iAs(III) (8.90E+03 copies/OD_600_), respectively (Fig. 3a). This result is deemed robust because, in the absence of iAs(III), the trend follows the opposite direction, i.e., *arsM* expression is highest in the no dilution (100% RCB) condition (Fig. 3a). We attribute the latter trend to imperfect normalization of *arsM* expression (we presume that the expression of *arsM* gene in all dilutions should be the same without iAs(III)) and biases stemming from the differences in expression in cells growing in substrate-replete *vs*. substrate-depleted conditions. However, these biases only strengthen the findings reported in the presence of iAs(III) because they would tend to decrease the expression of *arsM* in the higher dilution conditions, while the finding reports the highest transcript number in those conditions.

**FIG 3.**
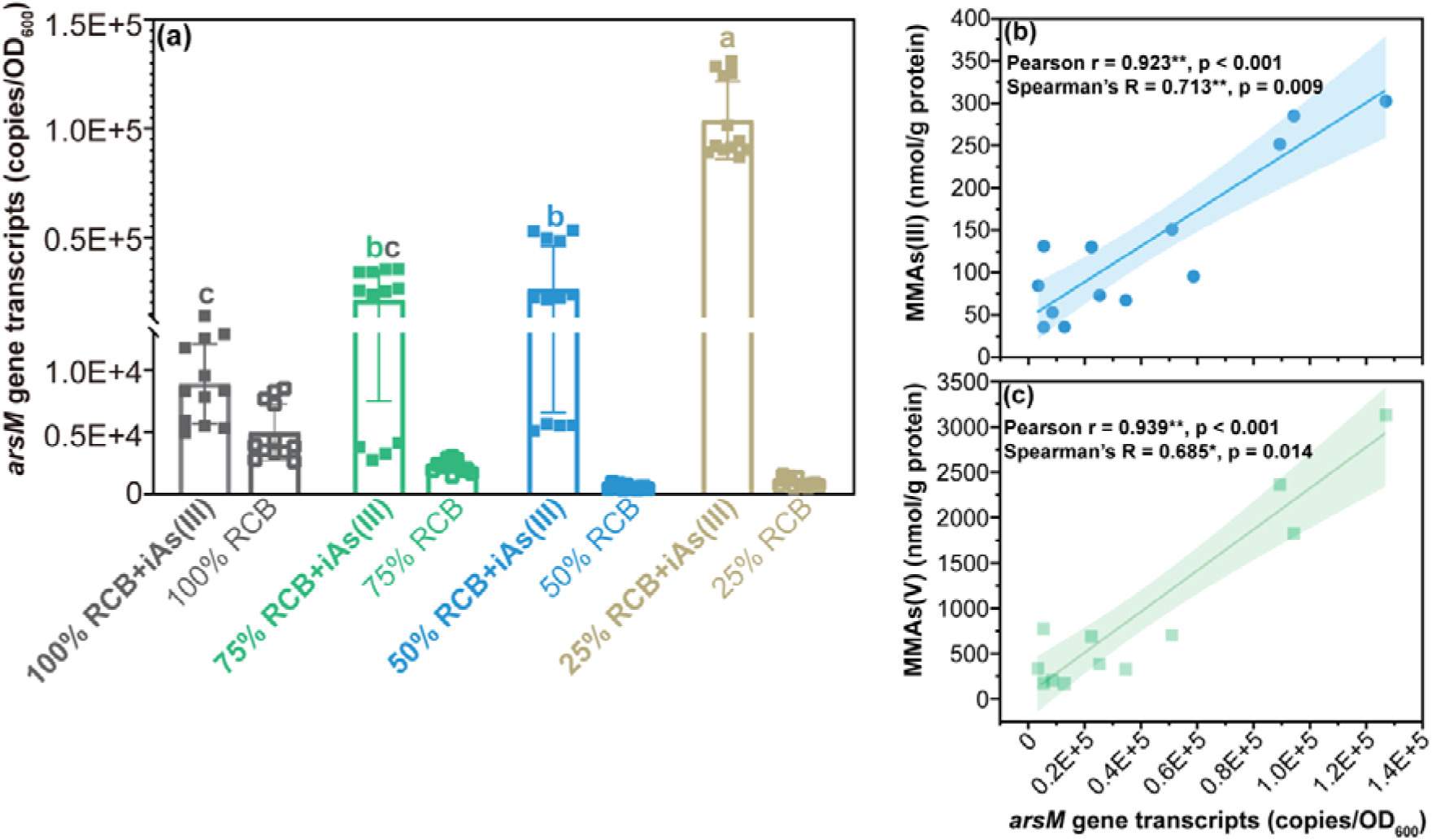
(a) Transcripts of *arsM* gene of *Paraclostridium bifermentans* strain EML in anaerobic RCB dilutions (100%, 75%, 50%, or 25% RCB) in the presence and absence of 25 μM iAs(III) at 8 hours of incubation. (b) and (c) Correlation analysis of *arsM* gene transcripts and concentrations of MMAs(III) (no oxidation), and MMAs(V) (post-oxidation) at 8 hours of incubation. Different letters showed significant difference between RCB dilutions at *p* < 0.05. Individual values for each biological replicate are shown in Data Tables 3 and 20.

Furthermore, correlation analysis suggested a significant (*p* < 0.05) positive correlation between *arsM* gene transcripts and the concentrations of aqueous MMAs(III) (no-oxidation) (Fig. 3b) and aqueous MMAs(V) (post-oxidation) (Fig. 3c). Similarly, the correlation between *arsM* gene transcript number and MMAs(III) content was also supported when using the second experimental method for absolute transcript quantification, which relies on using the same amount of total RNA for reverse transcripts regardless of biomass amount used for the extraction (Fig. S14).

### Impact of strain EML on *E. coli* growth rate

Next, we sought to probe the direct impact of As methylation by strain EML on other microorganisms. Confirmation of As(III)-resistance in *E. coli*, MMAs(III)-sensitivity in WT *E. coli*, and MMAs(III)-resistance in *E. coli* expressing *arsP* (hereafter, ArsP *E. coli*) is provided in Fig. S1-S3 and Texts SR1-SR2, and optimization of the co-culture ratio for incubations in Fig. S4-S6 and Text SR3. WT or ArsP *E. coli* in anaerobic co-culture with strain EML exhibited similar patterns of growth, with rapid growth within 10 hours of incubation, followed by a decline in copy number (Fig. 4a). The growth of ArsP *E. coli* was significantly higher (*p* < 0.01) than that of WT *E. coli* (Fig. 4a), while the growth of strain EML did not significantly differ between the two co-culture treatments (Fig. 4b). Thus, the growth difference between WT *E. coli* and ArsP *E. coli* in anaerobic co-culture with strain EML cannot be explained by growth-rate differences in strain EML during the co-culture period. We propose that the growth rate difference is attributable to the production of toxic MMAs(III) that inhibits the growth of WT *E. coli*, but negligibly affects that of ArsP *E. coli* (Fig. S1 and S2). In addition, the growth of strain EML alone was greater than that of strain EML in co-culture with either *E. coli* strain (Fig. 4b), and it could also be reasonably explained by the competition for growth substrate in co-culture.

**FIG 4.**
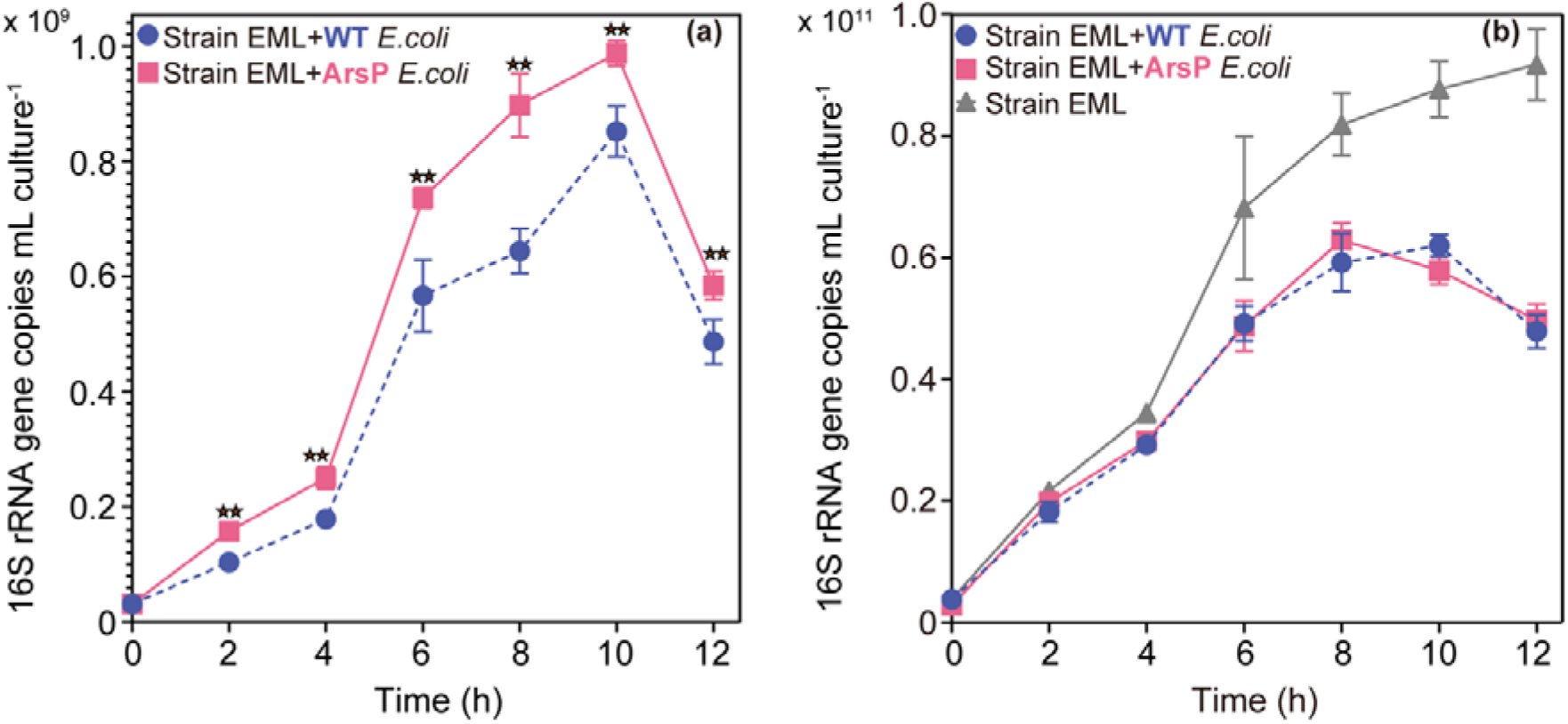
(a) Growth curves (16S rRNA gene copy number) of *Escherichia coli* K-12 wild-type strain MG1655 (WT *E. coli*) and engineered WT *E. coli* harboring a MMAs(III)-resistance gene (*arsP*) (ArsP *E. coli*) in anaerobic co-culture with *Paraclostridium bifermentans* strain EML in anoxic RCB with 25 μM iAs(III). (b) Growth curves (16S rRNA gene copy number) of *Paraclostridium bifermentans* strain EML in anerobic co-culture systems as described above. Two-star symbols represent statistical significance at *p* < 0.01. Individual values for each biological replicate are shown in Data Table 4.

### MMAs(III) production and *arsM* gene transcription during co-culture

From an ecological perspective, we would expect higher MMAs(III) production by strain EML when co-culturing with ArsP *E. coli* than WT *E. coli*, in order to thwart competition for nutrients with the faster growing strain. Indeed, MMAs(III) was found to be the dominant methylated As species and it increased gradually along with the decrease of iAs(III) during the co-culture period (Fig. S15). After normalization to biomass (16S rRNA gene copy number of strain EML) (Fig. 5a), we observed that MMAs(III) concentrations increased in the following order: strain EML < strain EML + WT *E. coli* < strain EML + ArsP *E. coli* (Fig. 5a).

**FIG 5.**
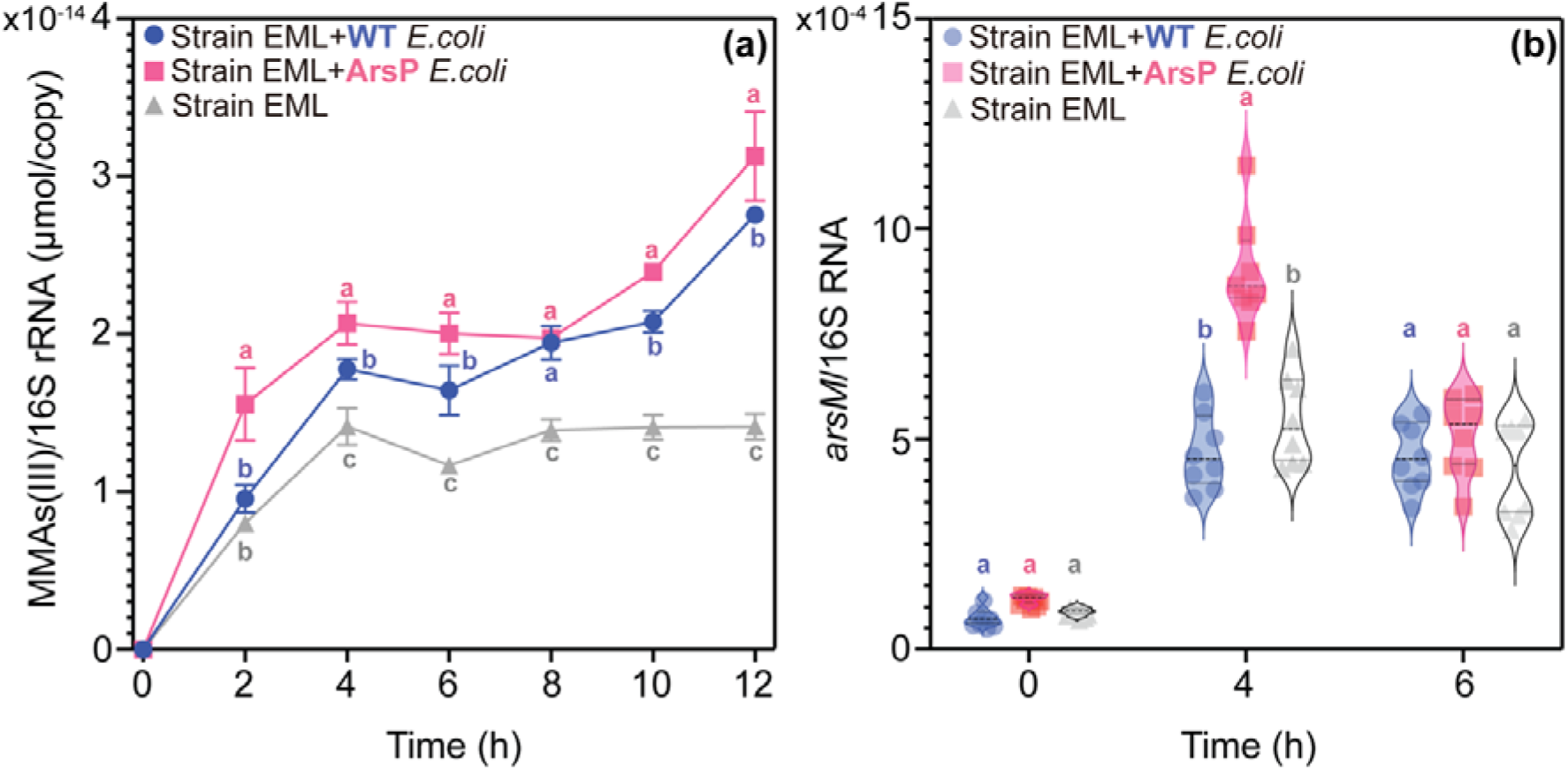
(a) Time-dependent concentration of aqueous MMAs(III) (normalized to 16S rRNA gene copies) in anaerobic co-culture *Paraclostridium bifermentans* strain EML with either WT *E. coli* or ArsP *E. coli* in anoxic RCB with 25 μM iAs(III). (b) Transcripts of *arsM* gene of strain EML (normalized to 16S rRNA gene copies) in anaerobic co-culture systems as described above at 0, 4, and 6 hours of incubation. Different letters indicate significant difference at *p* < 0.05. Individual values for each biological replicate are shown in Data Tables 5 and 21.

RT-qPCR was further used to investigate how substrate competition would impact the transcription of the *arsM* gene in anaerobic co-culture systems. The abundance of transcribed *arsM* gene in strain EML was measured during the exponential growth phase (4 h and 6 h) in anoxic co-culture systems (Fig. 5b). It is clear that the transcription of *arsM* gene in strain EML (normalized to its 16S rRNA gene copy number) was significantly (*p* < 0.05) higher in co-culture with ArsP *E. coli* than WT *E. coli* at 4 h, at mid-exponential phase (Fig. 5b). This difference in expression is consistent with more MMAs(III) produced in the strain EML and ArsP *E. coli* co-culture than that in the strain EML and WT *E. coli* system (Fig. 5a). Additionally, strain EML alone also produces less MMAs(III) than eithe*r E. coli* co-culture system but its *arsM* expression does not significantly differ from that of the co-culture including WT *E. coli* (Fig. 5b).

## DISCUSSION

In this study, we provide evidence that trivalent monomethylated As, MMAs(III) is produced as the dominant methylated As species by the recently isolated anaerobic As-methylating bacterium, *Paraclostridium bifermentans* strain EML, during growth in the presence of iAs(III) (Fig. 2). The concentration of MMAs(III) increases during the exponential growth phase and remains stable in the stationary phase (Fig. 1 and 2). This strongly suggests that anaerobic As methylation resulted from the activity of a functional ArsM from strain EML, rather than by the fortuitous methylation of As owing to the release of methyltransferases upon cell lysis as evidenced for methanogens in a previous study (22). An *arsM*-containing *ars* operon (*arsM*-*acr3*-*MPPE*-*arsR*1) was identified in strain EML (Fig. S7), supporting this interpretation.

A major question remains: what is the ecological function of generating a product (MMAs(III)) that is more toxic than the substrate (iAs(III))? At first glance, this would appear to be deleterious to the microorganism. However, this process could result in a beneficial outcome if two conditions are fulfilled. The first condition is that trivalent methylated As compounds serve as antibiotics to inhibit other potentially competing microorganisms (21). Because MMAs(III) and DMAs(III) are thermodynamically stable under anoxic conditions, they persist sufficiently long to be effective as antibiotics for anaerobes. The second condition is that MMAs(III), which is produced intracellularly, is exported to the extracellular space, precluding self-toxicity. If the rate of efflux of MMAs(III) is greater than that of iAs(III), As methylation would represent a net detoxification process. The conditions propitious for anaerobic As methylation remain elusive. A comparative study of As methylation across aerobic and anaerobic microorganisms revealed that, despite encoding a functional ArsM, anaerobes did not necessarily methylate iAs(III) (22).

This observation was partially attributed to the efficient efflux of iAs(III) in anaerobes (but not in aerobes), preventing sufficient accumulation of iAs(III) intracellularly for methylation to occur. Knocking out the *acr3* gene, encoding the iAs(III)-specific efflux pump in the anaerobe *Clostridium pasteurianum*, resulted in an obvious increase in intracellular iAs(III) but, overall, the As methylation efficiency improved little (22). A recent paper showed an increase in As methylation (under oxic conditions) upon knocking out the iAs(III) transporter gene *arsB* in cells of *E. coli* expressing *arsM* (30). Nonetheless, while iAs(III) influx is an important control on As methylation, we surmised that, in addition to intracellular iAs(III), other factors may control anaerobic As methylation. If the microbial warfare hypothesis for anaerobic As methylation holds, anaerobic As methylation might be triggered by an environmental signal suggesting obstacles to optimal growth (e.g., limited resources) or by specific metabolites produced by other microbial community members signaling their presence as potential competitors (22, 24).

Here, the first aim was to investigate the impact of substrate competition on anaerobic As methylation by generating growth substrate limitations. This point was probed by growing strain EML in dilutions of RCB medium and measuring the extent of As methylation. We found opposing trends between growth substrate content and concentration of aqueous MMAs(III) (Fig. 2) or *arsM* gene transcript numbers in the presence of iAs(III) (Fig. 3). Strain EML grown at lower growth substrate conditions produced higher (protein-normalized) concentrations of MMAs(III) (Fig. 2) and exhibited higher (protein-normalized) expression of the *arsM* gene (Fig. 3), suggesting that the cells responded to growth substrate limitation by increasing As methylation. Taken together, these results demonstrate an important role for growth substrate availability in regulating anaerobic As methylation by strain EML. The potential responses by strain EML to substrate availability at the cellular level are summarized in Fig. 6a: once strain EML senses signals from substrate depletion or other potential competitors, as a response, it activates the As-methylating system, particularly by up-regulating the expression of *arsM* gene to produce enough MMAs(III) and cope with use of growth substrates by competitors. Meanwhile, the expression of the MMAs(III) efflux permease gene *arsP* could also be up-regulated, leading to efficient MMAs(III) extrusion to the environment to avoid self-toxicity. In addition, we observed the accumulation of intracellular As(III) preferentially in the low-growth substrate conditions, confirming the close relationship between intracellular iAs(III) and high As methylation potential, as previously evidenced (22). One possible explanation for the highly accumulated intracellular As(III) could be attributable to the inhibition and repression of the iAs(III) efflux permease encoded by the gene *acr3* in strain EML under substrate-limited condition, i.e., less efficient iAs(III) efflux system leads to higher intracellular iAs(III) buildup in cytoplasm, facilitating further anaerobic As methylation. Another possible explanation is that the uptake of iAs(III) via the aquaporin channels into the cell could be stimulated under substrate-limited conditions (low dissolved organic matter) compared to that under substrate-sufficient conditions (high dissolved organic matter) (31).

**FIG 6.**
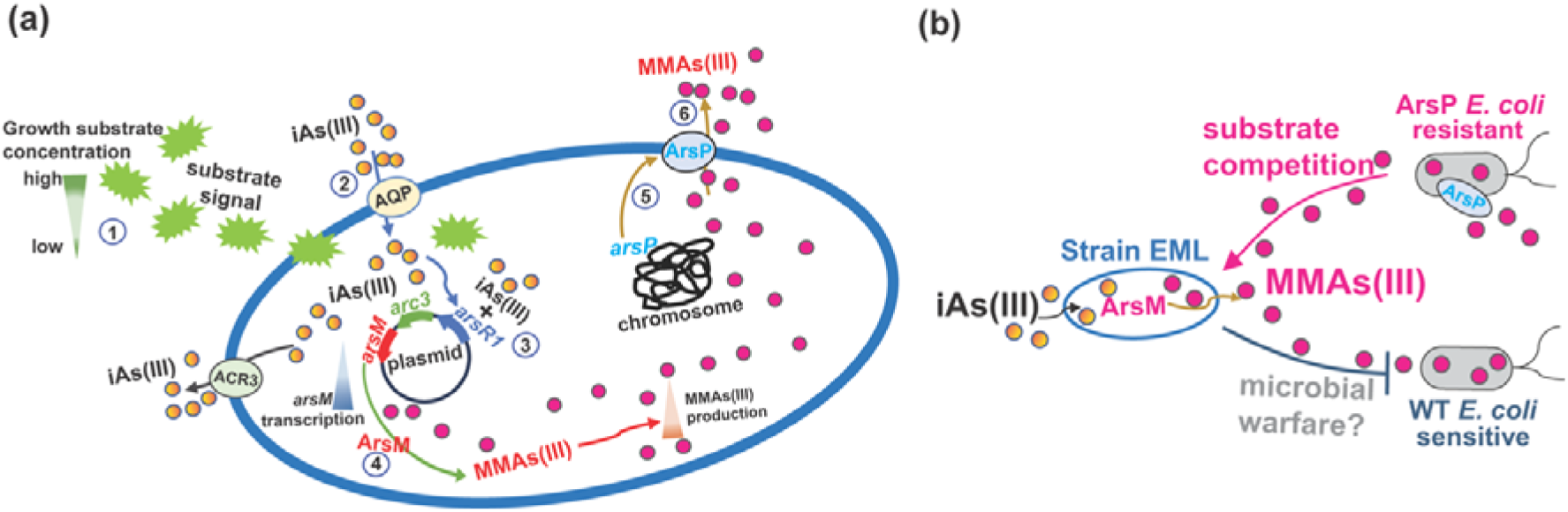
(a) Schematic of *Paraclostridium bifermentans* strain EML under anoxic growth substrate-limiting condition. Strain EML increases MMAs(III) production. Numbers denote steps in the response by strain EML: l7 Receiving signal of substrate depletion, l7 Entry of iAs(III) into cell, l7 Repressor ArsR1 binds to iAs(III) allowing transcription of *arsR*1, *acr*3, and *arsM* genes, l7 ArsM methylates iAs(III) to MMAs(III), l7 Transcription of *arsP* is activated (mechanism unknown) and ArsP produced, and l7 MMAs(III) is effluxed via ArsP. (b) Schematic of the interaction *P. bifermentans* strain EML with competing *E. coli*. Microbial inhibition and competition between MMAs(III)-producing strain EML and MMAs(III)-resistant and -sensitive *E. coli* strains under anoxic conditions.

Next, direct evidence of microbial inhibition by microbial MMAs(III) production was sought by the anaerobic co-culture of strain EML with either MMAs(III)-sensitive WT *E. coli* or MMAs(III)-resistant *E. coli* (i.e., ArsP *E. coli*) in RCB medium (Fig. 4 and 5). We found a two-way interaction between strain EML and *E. coli* (as outlined in Fig. 6b): on the one hand, MMAs(III) produced by strain EML inhibits the growth of WT *E. coli* to a greater extent than that of ArsP *E.coli* (Fig. 4 and 6b), supporting the microbial warfare hypothesis. On the other hand, we interpret less growth by WT *E. coli* to result in less substrate depletion as compared to ArsP *E. coli* and, in turn, lower expression of *arsM* in strain EML (Fig. 5 and 6b). However, it is also conceivable that other factors (e.g., signaling) cause lower *arsM* expression. Therefore, it is reasonable to conclude that MMAs(III) production inhibits microorganisms unprepared to detoxify it and that strain EML responds to substrate limitation by increasing *arsM* expression and, thus MMAs(III) production. While more MMAs(III) is produced by strain EML co-cultured with ArsP *E. coli* than WT *E. coli*, the growth of strain EML is comparable in the presence of either *E. coli* strain (Fig. 4b). A proposed explanation for this observation is that, while MMAs(III) production is a response to substrate limitation, it does not necessarily thwart the competitor (i.e., the prey) sufficiently to impact resource use, and thus growth. Indeed, the impact of resource competition is evident from comparing the growth of strain EML with or without *E. coli* (Fig. 4b).

Efflux of MMAs(III) is required for effective delivery of this antibiotic to other microorganisms and, to avoid self-toxicity (as stated in condition 2 above). Specific/nonspecific MMAs(III) efflux permease genes *arsP* (32) and *arsK* (33) are known and the co-evolution of *arsM* and *arsP* was previously evidenced, suggesting a strategy of MMAs(III) efflux by MMAs(III)-producing and MMAs(III)-resistant microorganisms (18–20). The identification of two chromosomally encoded *arsP* genes in strain EML indicates that it may be capable of effluxing MMAs(III) to the extracellular space (Fig. S16) and variable *arsP* expression across conditions (not measured) may account for the similarity in the extent of *arsM* expression by strain EML growth with WT *E. coli* and strain EML alone.

In this study, we provide direct evidence for the role of growth substrate competition in anaerobic As methylation by strain EML. Furthermore, we evidence a feedback loop, by which a bacterium resistant to MMAs(III) enhances its production, presumably through enhanced *arsM* expression as a result of substrate limitation. Therefore, the work uncovers complex interactions between an anaerobic As methylator and potential competitors. Nevertheless, in addition to substrate limitation, nevertheless, other as-yet-identified signals/factors in the rice-soil system could also contribute to the high anaerobic As methylation activity in flooded paddy fields. Further work is clearly needed, first to uncover the mechanism of regulation of *arsM* gene expression by the growth substrate concentration and second to elucidate other factors that may control anaerobic As methylation. Substantial understanding of the controls on anaerobic As methylation is required for the development of strategies to limit As methylation in rice paddy soils.

## METERIALS AND METHODS

### Growth Experiment

The anaerobic As-methylating bacterium, *Paraclostridium bifermentans* strain EML (henceforth strain EML) was previously isolated from an anaerobic paddy soil enrichment (24, 34). To investigate how growth substrate availability affects As methylation activity, dilutions (v/v) of Reinforced Clostridial Broth (RCB) (Oxoid Ltd) medium in Milli-Q water (100% RCB, 75% RCB, 50% RCB, and 25% RCB) were prepared in 120 mL serum bottles containing 50 mL medium (Supporting information Table S1). The medium was brought to a boil for 5 min to remove O_2_, then cooled down under a 100% N_2_ gas flow to room temperature and dispensed into individual culture serum bottle under the same N_2_ atmosphere. The bottles were then sealed with sterile rubber stoppers and crimped with aluminum caps and the headspace was flushed with 100% N_2_ to ensure anaerobic conditions before autoclaving at 121°C for 15 mins.

A pre-culture of strain EML was grown in RCB anaerobically to mid-exponential growth phase. Strain EML (∼0.5 mL) was inoculated into each RCB dilution containing 25 μM iAs(III) as sodium arsenite (or into the equivalent no-iAs(III) control) in triplicate using sterile N_2_-flushed syringes and needles. The inoculum represents approximately 1% of the total volume (v/v). All the bottles were incubated at 30°C in the dark without shaking. A total of eight experimental conditions were selected (Table S1). At selected time points and for each condition, triplicate bottles were sampled for growth, which was quantified using both optical density at 600 nm (OD_600_) and total protein content estimated using a BCA protein assay kit (Thermo Scientific, MA, USA). For quantification of the expression of the *arsM* gene, triplicate cultures were sampled for RNA extraction at 8 and 24 h. To test the stability of MMAs(III) in RCB medium, additional abiotic control experiments (including fresh/spent RCB medium supplemented with MMAs(III), and chemical reaction between MMAs(III) and sulfide) were performed in duplicate (see Supporting Information Text Methods SM1).

### Arsenic Speciation

At each time point and for each condition, aqueous and intracellular As speciation and total As were measured. Samples for aqueous As species and total As were obtained from 1 mL of culture collected with sterile, N_2_-flushed syringes and needles, filtered through 0.22 μm cellulosic membrane filters, and stored in 1 mL 1% HNO_3_ (≥ 69 %, Honeywell Fluka). Additionally, to analyze As species and total As post-sample oxidation, another 1 mL of culture was obtained as described above and oxidized by adding 10% (v/v) hydrogen peroxide (w/v) (H_2_O_2_, 30%, Reactolab SA) and was stored overnight in a 1% HNO_3_ solution. For soluble intracellular As species, 1 mL of culture was collected, the cells were pelleted at 8,000 *g* for 5 min, and stored at -20°C until use. To release soluble intracellular As, the cell pellets were lysed in a lysis buffer (0.1% Triton X-100, 0.1% SDS, 10 mM EDTA, and 1 mM Tris-HCl) at 95°C for 15 min by vortexing every 3 min (Viacava et al., 2020). The lysed cell suspension was subsequently centrifuged at 8,000 *g* for 5 min, and the pellet was resuspended in 200 μL 1 x PBS buffer and used for protein determination as described above. The supernatant was filtered through 0.22 μm filters and reserved for As speciation and total As analysis.

Both aqueous and soluble intracellular As speciation were determined by high performance liquid chromatography and inductively coupled plasma mass spectrometry (HPLC-ICP-MS) on an Agilent 8900 ICP-QQQ instrument. A previously described anion exchange protocol using the step-gradient elution mode with an As Spec anion exchange fast column (50 mm x 4.0 mm, PrinCen, Guangzhou, China) was used.^21^ Six As standards were available: MMAs(III) as methyldiiodoarsine (Santa Cruz Biotechnology Inc.), TMAs(V)O as trimethyl arsine oxide (Argus Chemicals Srl., Italy), DMAs(V) as sodium dimethylarsinate (ABCR, Germany), MMAs(V) as monomethylarsonic acid (Chemservice, PA, USA), As(III) as sodium arsenite (NaAsO_2_) (Sigma-Aldrich, MO, USA),and As(V) as sodium arsenate dibasic heptahydrate (Na_2_HAsO_4_·7H_2_O) (Sigma-Aldrich, MO, USA). In addition, monomethylmonothioarsonic acid (MMMTAs(V)) was synthesized as previously described (Text SM1) (35). Total aqueous and soluble intracellular As concentrations were measured using the same ICP-MS instrument in stand-alone mode (24).

### RNA extraction and RT-qPCR

After sampling, each culture was amended with RNAprotect Bacteria Reagent (Qiagen, Hilden, Germany) following the manufacturer’s recommendations to stabilize RNA and prevent its degradation. The RNeasy Mini Kit (Qiagen) was used following the manufacturer’s instructions with an initial sample preparation protocol from the Qiagen RNAprotect bacteria reagent handbook. Protocol 5 (enzymatic lysis, proteinase K digestion, and mechanical disruption of bacteria) was employed for cell lysis prior to RNA purification. Genomic DNA digestion was completed during RNA purification using the RNase-Free DNase set (Qiagen). Reverse transcription was performed with QuantiTect Rev. Transcription Kit (Qiagen). Detailed information about designing a specific *arsM* gene primer set, optimizing the PCR amplification condition, and constructing an *arsM* plasmid to use for standard curve are described in Text SM2. The RT-qPCR was carried out in a Mic PCR system (Bio Molecular Systems, Mic) using SYBR Green Master Mix. The reactions (10 μL total volume) contained 5 μL of 2 × SensiFAST™ SYBR No-ROX Kit (Bioline, London, UK), 0.2 μM of each *arsM* gene primer, 2.5 μL of cDNA, and 1% (v/v) bovine serum albumin (BSA) (Sigma). A 10-fold dilution series containing 10^7^-10^1^ copies of strain EML *arsM* plasmid DNA was used to generate a standard curve. All samples were run in quadruplicates. A NRT (no-reverse transcriptase control) and a NTC (no template control) were both included as negative controls.

### Quantification of *arsM* gene transcripts

To get a comprehensive understanding of *arsM* gene expression under variable substrate conditions, both relative and absolute quantification methods were attempted. For relative quantification, the specific primer sets for the housekeeping genes considered and the corresponding amplification conditions are shown in Table S2. However, these genes are not used in this study because they do not meet the minimum requirement for a stable reference gene (see results in Section *arsM* gene transcription). For absolute quantification, we were concerned that the mRNA yield would be variable across conditions due both to biological reasons (e.g., rate and extent of growth) and biases introduced by RNA extraction. To obtain robust results, we compared two methods of absolute quantification. The first method entailed adjusting biomass for each culture prior to RNA extraction to ensure that RNA was extracted from the same amount of biomass (OD_600_) regardless of conditions. The same volume of total RNA was used for reverse transcription, and the expression data (*arsM* copy numbers) were presented relative to OD_600_ (biomass). The second method consisted of quantifying extracted total RNA and using the same amount of RNA from all conditions in reverse transcription. The expression data were then normalized to the corresponding protein concentration.

### Anaerobic co-culture system

To provide direct evidence of the effects of substrate competition among microbes and associated microbe-microbe interactions on anaerobic As methylation, anaerobic co-culture systems (predator-prey systems) were established. The “predator” was strain EML (24). One “prey” was wild type *Escherichia coli* K-12 strain MG1655 (WT), which is sensitive to MMAs(III) (Fig. S1 and S2) and resistant to As(III) (36) (Fig. S3 and Text SM3). The other was the same strain in which the *arsP* gene was integrated into the chromosome by the mini-Tn7-based gene integration method (37) and controlled by the promoter from the *fnr*S gene (Texts SM4-SM6), a highly conserved, anaerobically induced small RNA (38). The *arsP* gene encodes ArsP, a MMAs(III) efflux permease that extrudes trivalent organoarsenicals from cells (32). The expression of *arsP* in *E. coli* confers MMAs(III) resistance under anoxic conditions. The confirmation of As(III)-resistance in WT *E. coli* and *E. coli* expressing *arsP* (hereafter, ArsP *E. coli*), MMAs(III)-sensitivity in WT *E. coli*, and MMAs(III)-resistance in ArsP *E. coli* are provided in Fig. S1-S3 and Supporting Information Text Results SR1-SR2.

Co-culture treatments consisting of (i) strain EML + WT *E. coli* + 25 μM iAs(III), (ii) strain EML + ArsP *E. coli* + 25 μM iAs(III), or (iii) the control with only strain EML + 25 μM iAs(III) were conducted in triplicate as described above in anaerobic serum bottles containing 50 mL 100% RCB medium. Given the difference in growth rate between strain EML and *E. coli*, variable inoculation ratios between the co-culture members were tested and the optimal ratio found to be 10% EML (v/v), that is, the cell pellet from a 5 mL exponential phase culture of strain EML in RCB and 50 μL exponential phase culture of *E. coli* (WT or ArsP *E. coli*) in 50 mL of RCB (Fig. S4-S6; Texts SM7 and SR3). The optimal co-culture cell ratio is the ratio which is required for enough strain EML cells present in the co-culture system to produce at least 1 μM MMAs(III), otherwise *E. coli* would dominate the co-culture system owing to the distinct growth rates between strain EML and *E. coli*. During the anaerobic co-culture period, aqueous As speciation, the growth rate, and *arsM* gene transcripts were measured by HPLC-ICP-MS, qPCR (by quantification of the 16S rRNA gene copy numbers of strain EML, and the two *E. coli* strains), and RT-qPCR, respectively.

## Supporting information

supporting methods SM1-SM7; supporting results SR1-SR4; supplemental figures S1-S18

supporting methods SM1-SM7; supporting results SR1-SR4; supplemental figures S1-S18; supplemental tables S1-S3.

Raw dataset for all figures

## ACKNOWLEDGMENTS

The authors would like to acknowledge Nicolas Jacquemin for identification and visualization of arsenic-methylating related genes in strain EML, Colin Volet for characterization of the methylated-thiolated arsenic standards and the EPFL Central Environmental Laboratory. This work was supported by the Swiss National Science Foundation (SNSF) NCCR Microbiomes (565196) and the National Science Foundation of China (42277119).

## Supporting Information

Additional supporting information can be found online in the Supporting Information section at the end of this article.

## REFERENCES

1. Páez-Espino D, Tamames J, de Lorenzo V, Cánovas, D. (2009) Microbial responses to environmental arsenic. Biometals, 22, 117−130.

2. Stolz JF, Basu P, Santini JM, Oremland RS. (2006) Arsenic and selenium in microbial metabolism. Annual Review of Microbiology, 60, 107−130.

3. Tamaki S, Frankenberger WT. (1992) Environmental Biochemistry of Arsenic. *In* Reviews of Environmental Contamination and Toxicology: Continuation of Residue Reviews; Ware, G. W., Eds.; Springer New York: New York, pp 79−110.

4. Zhu YG, Xue XM, Kappler A, Rosen BP, Meharg AA. (2017) Linking genes to microbial biogeochemical cycling: lessons from arsenic. Environmental Science & Technology, 51, 7326–7339.

5. Slyemi D, Bonnefoy V. (2012) How prokaryotes deal with arsenic. Environmental Microbiology Reports, 4, 571−586.

6. Zhu YG, Yoshinaga M, Zhao FJ, Rosen BP. (2014) Earth abides arsenic biotransformations. Annual Review of Earth and Planetary Sciences, 42, 443−467.

7. Davis MA, Signes-Pastor AJ, Argos M, Slaughter F, Pendergrast C. Punshon T, Gossai A, Ahsan H, Karagas MR. (2017) Assessment of human dietary exposure to arsenic through rice. Science of The Total Environment, 586, 1237−1244.

8. Williams P, Price A, Raab A, Hossain S, Feldmann J, Meharg AA. (2005) Variation in arsenic speciation and concentration in paddy rice related to dietary exposure. Environmental Science & Technology, 39, 5531−5540.

9. Meharg AA, Williams PN, Adomako E, Lawgali YY, Deacon C, Villada A, Cambell RC, Sun GX, Zhu YG, Feldmann J. (2009) Geographical variation in total and inorganic arsenic content of polished (white) rice. Environmental Science & Technology, 43, 1612−1617.

10. Zavala YJ, Gerads R, Gürleyük H, Duxbury JM. (2008) Arsenic in rice: II. Arsenic speciation in USA grain and implications for human health. Environmental Science & Technology, 42, 3861–3866.

11. Chhabra R, Goyal P, Singh T, Vij L. (2021) Understanding straighthead: a complex physiological disorder of rice (Oryza sativa L.). Acta Physiologiae Plantarum, 43, 1−14.

12. Tang Z, Wang Y, Gao A, Ji Y, Yang B, Wang P, Tang Z, Zhao FJ. (2020) Dimethylarsinic acid is the causal agent inducing rice straighthead disease. Journal of Experimental Botany, 71, 5631–5644.

13. Dai J, Chen C, Gao AX, Tang Z, Kopittke PM, Zhao FJ. (2021) Dynamics of dimethylated monothioarsenate (DMMTA) in paddy soils and its accumulation in rice grains. Environmental Science & Technology, 55, 8665−8674.

14. Challenger F. (1951) Biological methylation. Advances in Enzymology and Related Areas of Molecular Biology, 12, 429−491.

15. Marapakala K, Packianathan C, Ajees AA, Dheeman DS, Sankaran B, Kandavelu P, Rosen BP. (2015) A disulfide-bond cascade mechanism for arsenic(III) *S*-adenosylmethionine methyltransferase. Acta Crystallographica Section D, 71, 505−515.

16. Qin J, Rosen BP, Zhang Y, Wang G, Franke S, Rensing C. (2006) Arsenic detoxification and evolution of trimethylarsine gas by a microbial arsenite *S*-adenosylmethionine methyltransferase. Proceedings of the National Academy of Sciences of the United States of America, 103, 2075−2080.

17. Qin J, Lehr CR, Yuan C, Le XC, McDermott TR, Rosen BP. (2009) Biotransformation of arsenic by a Yellowstone thermoacidophilic eukaryotic alga. Proceedings of the National Academy of Sciences of the United States of America, 106, 5213−5217.

18. Chen SC, Sun GX, Rosen BP, Zhang SY, Deng Y, Zhu BK, Rensing C, Zhu YG. (2017) Recurrent horizontal transfer of arsenite methyltransferase genes facilitated adaptation of life to arsenic. Scientific Reports, 7, 1−11.

19. Chen SC, Sun GX, Yan Y, Konstantinidis KT, Zhang SY, Deng Y, Li XM, Cui HL, Musat F, Popp D, Rosen BP, Zhu YG. (2020a) The Great Oxidation Event expanded the genetic repertoire of arsenic metabolism and cycling. Proceedings of the National Academy of Sciences of the United States of America, 117, 10414−10421.

20. Chen J, Yoshinaga M, Rosen BP. (2019) The antibiotic action of methylarsenite is an emergent property of microbial communities. Molecular Microbiology, 111, 487−494.

21. Chen J, Rosen BP. (2020b) The arsenic methylation cycle: how microbial communities adapted methylarsenicals for use as weapons in the continuing war for dominance. Frontiers in Environmental Science, 8, 43.

22. Viacava K, Meibom KL, Ortega D, Dyer S, Gelb A, Falquet L, Minton NP, Mestrot A, Bernier-Latmani R. (2020) Variability in arsenic methylation efficiency across aerobic and anaerobic microorganisms. Environmental Science & Technology, 54, 14343−14351.

23. Jiang Z, Shen X, Shi B, Cui MJ, Wang Y H, Li P. (2022) Arsenic mobilization and transformation by ammonium-generating bacteria isolated from high arsenic groundwater in Hetao Plain, China. International Journal of Environmental Research and Public Health, 19, 9606.

24. Viacava K, Qiao JT, Janowczyk A, Poudel S, Jacquemin N, Meibom KL, Shrestha HK, Reid MC, Hettich RL, Bernier-Latmani R. (2022) Meta-omics-aided isolation of an elusive anaerobic arsenic-methylating soil bacterium. The ISME Journal, 16, 1740–1749.

25. Ghoul M, Mitri S. (2016) The ecology and evolution of microbial competition. Trends in Microbiology, 24, 833–845.

26. Mallon CA, Poly F, Le Roux X, Marring I, van Elsas JD, Salles JF. (2015) Resource pulses can alleviate the biodiversity–invasion relationship in soil microbial communities. Ecology, 96, 915–926.

27. Currier JM, Saunders RJ, Ding L, Bodnar W, Cable P, Matoušek T, Creed JT, Stýblo M. (2013) Comparative oxidation state specific analysis of arsenic species by high-performance liquid chromatography-inductively coupled plasma-mass spectrometry and hydride generation-cryotrapping-atomic absorption spectrometry. Journal of Analytical Atomic Spectrometry, 28, 843−852.

28. Scheer J, Findenig S, Goessler W, Francesconi KA, Howard B, Umans JG, Pollak J, Tellez-Plaza M, Silbergeld EK, Guallar E. (2012) Arsenic species and selected metals in human urine: validation of HPLC/ICPMS and ICPMS procedures for a long-term population-based epidemiological study. Analytical Methods, 4, 406−413.

29. Yathavakilla SKV, Fricke M, Creed PA, Heitkemper DT, Shockey NV, Schwegel C, Caruso JA, Creed JT. (2008) Arsenic speciation and identification of monomethylarsonous acid and monomethylthioarsonic acid in a complex matrix. Analytical Chemistry, 80, 775−782.

30. Chen J, Rosen BP. (2023) Arsenite methyltransferase diversity and optimization of methylation efficiency. Environmental Science & Technology, 57, 9754−9761.

31. Yoon H, Stenzler B, Abu-Ali L, Asta MP, Poulain AJ, Reid MC. (2023) Effect of iron and dissolved organic matter on bioavailable arsenite under anaerobic conditions. ACS EST Water, 3, 3676–3686.

32. Chen J, Madegowda M, Bhattacharjee H, Rosen BP. (2015) ArsP: a methylarsenite efflux permease. Molecular Microbiology, 98, 625−635.

33. Shi K, Li C, Rensing C, Dai X, Fan X, Wang G. Efflux transporter ArsK is responsible for bacterial resistance to arsenite, antimonite, trivalent roxarsone, and methylarsenite. Applied and Environmental Microbiology, 84, e01842–18.

34. Reid MC, Maillard J, Bagnoud A, Falquet L, Le Vo P, Bernier-Latmani R. (2017) Arsenic methylation dynamics in a rice paddy soil anaerobic enrichment culture. Environmental Science & Technology, 51, 10546−10554.

35. Kerl CF, Schindele RA, Brulggenwirth L, Colina Blanco AE, Rafferty C, Clemens S, Planer-Friedrich B. (2019) Methylated thioarsenates and monothioarsenate differ in uptake, transformation, and contribution to total arsenic translocation in rice plants. Environmental Science & Technology, 53, 5787–5796.

36. Carlin A, Shi W, Dey S, Rosen BP. (1995) The *ars* operon of Escherichia coli confers arsenical and antimonial resistance. Journal of Bacteriology, 177, 981–986.

37. Choi KH, Schweizer HP. (2006) mini-Tn7 insertion in bacteria with single attTn7 sites: example *Pseudomonas aeruginosa*. Nature Protocols, 1, 153–161

38. Roca I, Ballana E, Panosa A, Torrents E, Gibert I. (2008) Fumarate and nitrate reduction (FNR) dependent activation of the *Escherichia coli* anaerobic ribonucleotide reductase nrdDG promoter. International Microbiology, 11, 49–56.

