## supporting methods SM1-SM7; supporting results SR1-SR4; supplemental figures S1-S18 for "Growth substrate limitation enhances anaerobic arsenic methylation by *Paraclostridium bifermentans* strain EML"

**Anaerobic arsenic methylation as a microbial warfare strategy**

Main content: 11 text sections (supporting methods SM1-SM7; supporting results SR1-SR4), 20 figures (S1-S20), 24 tables (excel files), and references (33 pages).

**TEXT**

**Supporting Methods:**

**Text SM1: Chemical stability of MMAs(III) (abiotic controls).** All abiotic controls were conducted at 30°C in the dark without shaking. Three types of controls were prepared: (i) For fresh-RCB abiotic controls, systematic dilutions of RCB (100%, 75%, 50%, or 25% RCB) were prepared in 120 mL serum bottles containing 50 mL medium following the anaerobic procedures described in the Materials and Methods section. MMAs(III) (3 μM) was added to each medium in duplicate using sterile, N_2_-flushed syringes and needles. (ii) For spent-RCB abiotic controls, a pre-culture of strain EML was grown in RCB anaerobically to mid-exponential growth phase, then the spent medium was prepared by filtering out the cells with a sterile Nalgene Rapid-Flow 0.1 μm PES membrane filter (Thermo Fisher) twice in an anoxic gas chamber (Coy Laboratory Products). Filtered spent medium (25 mL) was injected into the pre-autoclaved triplicate serum bottles with a 100% N_2_ atmosphere, followed by the addition of 3 μM MMAs(III) to each medium replicate using sterile, N_2_-flushed syringes and needles. (iii) To assess the chemical reaction between MMAs(III) and sulfide, anaerobic sterile MQ water was prepared in duplicate serum bottles, amended with 3 μM MMAs(III) and then 6 μM sodium sulfide using sterile, N_2_-flushed syringes and needles.

For all cases, samples to analyze aqueous As speciation were obtained from 1 mL of solution collected with sterile, N_2_-flushed syringes and needles, filtered through 0.22 μm filters, and stored at -80°C after directly flash-freezing in liquid nitrogen. To probe whether methylated and thiolated As species were formed, As species post-sample oxidation in the MMAs(III)-sulfide system was oxidized by adding 10% (v/v) H_2_O_2_ and was kept overnight in a 1% HNO_3_ solution. As speciation was determined by HPLC-ICP-MS on the Agilent 8900 ICP-QQQ instrument (Agilent Technologies) using the previously described anion exchange protocol.^1^ Three As standards were prepared, including MMAs(III) and MMAs(V) (both commercially available), and MMMTAs(V) (was synthesized). The standard solution of monomethyl monothioarsonic acid (MMMTAs(V)) was prepared by reaction of MMAs(V) (CH_3_AsNa_2_O_3_) with Na_2_S according to a method previously described.^2^ The structure of synthesized MMMTA was determined by Waters Acquity-I-UPLC Class system (Waters Corporation, Milford, MA, USA) coupled with a Waters Vion IMS-QTof Mass Spectrometer equipped with LockSpray (Leucine-enkephalin (200 pg/µL). The purity was determined by using HPLC-ICP-MS and ICP−MS. The synthesized MMMTA contained 93.1% MMMTA, 3.4% MMA, 1.5% MMDTA, 1.9% arsenate, and 0.1% DMDTA.

**Text SM2: *arsM* PCR and plasmid standard.** The specific qPCR primers for strain EML *arsM* gene were designed by Benchling online research platform (https://benchling.com/faq). The primers were designed to be 18 to 23 bp in length and to generate amplicons in the size range of 80 to 200 bp. The primers for *arsM* gene from strain EML are: arsM-F: 5’-AAAAGCAGTCAAACCCGGCTCC-3’ and arsM-R: 5’-ACTTAGGCTTAGGGTGTGGAAACC-3’. To calculate qPCR efficiency, we established the standard curves with 10-fold dilutions of strain EML genomic DNA to ensure the primer set had an amplification efficiency within 90-110%.

The *arsM* gene PCR product of strain EML was amplified by PCR using the arsM-F and arsM-R primer set. The reaction mixture (50 μL) contained 25 μL 2 x GoTaq Green Master Mix (Promega), 0.2 μM each primer, 50 ng strain EML genomic DNA template, and 1% (v/v) bovine serum albumin (BSA) (Sigma). PCR amplification was performed in a Biometra Trio Thermocycler (Biometra, Göttingen, Germany) according to the following program: an initial denaturation at 95°C for 5 min, followed by 40 cycles of 5 s at 95°C, 20 s at 60°C and 20 s at 72°C; a final extension at 72°C for 5 min. The PCR product was purified using Wizard SV Gel and PCR Clean-Up System (Promega) and cloned into a pGEM-T Easy vector by the TA-Cloning method using white-blue screening to select positive clones (Promega). The cloned plasmid was purified using Wizard® Plus SV Minipreps DNA Purification System (Promega). The DNA insert was verified by PCR amplification using the vector-specific M13F and M13R primers and was further sequenced on an Applied Biosystems 310 genetic analyzer (Applied Biosystems, ABI).

**Text SM3: Validation of iAs(III)-resistance in *E. coli*.** To investigate As(III) resistance in *E. coli* strain MG1655 and rule out the possible confounding effects on growth caused by the presence of As(III) in co-culture systems, anaerobic microcosms were prepared by inoculating 120 mL serum bottles with 50 mL anaerobic 100% RCB medium. Pre-cultures of WT *E. coli* and ArsP-expressing *E. coli* (aka ArsP *E. coli*) were grown in RCB anaerobically to mid-exponential growth phase. Both WT and ArsP *E. coli* (~0.5 mL) were inoculated separately into each RCB medium supplied with 10, 15, and 25 μM As(III) in triplicate using sterile N_2_-flushed syringes and needles. At selected time points and for each condition, triplicate bottles were sampled and growth was measured as optical density at 600 nm (OD_600_) in a spectrophotometer. In addition, aqueous As species was also quantified at the beginning and the end of the incubation by HPLC-ICP-MS as described in the main text Materials and Methods section (Arsenic Speciation).

**Text SM4: Construction of recombinant MMAs(III)-resistant *E. coli*.** pTNS2 (#64968), pFLP3 (#64946), and pUC18R6KT-mini-Tn7T-Km (#64969) plasmids^3^ were purchased from Addgene. The ArsP protein from *Campylobacter jejuni*^4^ was reverse translated (using *E. coli* codon usage) and a gene block (gBlock) fragment (1084 bp) was synthesized by IDT (Integrated DNA Technologies). It contains the fnrS promoter (fnrSp) located upstream from the codon-optimized *arsP* gene (including 50 nt upstream the gene), which was flanked by two restriction enzyme recognition sites (SacI and EcoRI). To clone *fnrS*p-*arsP* into the delivery vector pUC18R6KT-mini-Tn7T (containing kanamycin (Kan) and ampicillin (Amp) resistance selectable markers), the vector was digested with EcoRI and SacI (New England BioLabs) and the linearized vector was purified by agarose gel electrophoresis using the Wizard SV Gel and PCR Clean-Up System (Promega). The *fnrS*p-*arsP* gBlock was digested with the same restriction enzymes and purified on column with the Wizard SV Gel and PCR Clean-Up System (Promega). Digested pUC18R6KT-mini-Tn7T vector and *fnrS*p-*arsP* gBlock were ligated overnight at 16°C with T4 DNA ligase (New England BioLabs). The ligation mixture was then transformed into *Escherichia coli DH5α λpir* by heat shock followed by selection on LB agar with 100 μg/mL Amp. A positive clone was selected and plasmid DNA extracted using the Wizard® Plus SV Minipreps DNA Purification Systems (Promega).

The integration of the *arsP* gene into the chromosome of strain MG1655 (in *att*Tn7, downstream of *glmS*) was achieved as described.^3^ Briefly, the pUC18R6KT-mini-Tn7T-arsP and the helper plasmid pTNS2 were electroporated into *E. coli* K12 strain MG1655. The transformants with insertions were selected on LB-Kan agar plates and positive single colonies were transferred on both LB-Kan and LB-Amp agar (confirm loss of the plasmid). Single Kan-resistant and Amp-sensitive clones were picked and the insertion of *arsP* gene into the genome of strain MG1655 was confirmed by colony PCR with glmS-down (genome-specific; 5'-GCACATCATCGAGATGCC) and neomycin resistance gene (*neo*) (5'-CGTTGGCTACCCGTGATATT) primers. To rule out any potential interference of the selection marker, neo gene, in the newly constructed *arsP*-harboring *E. coli* (aka ArsP *E. coli*) relative to WT *E. coli*, the gene was removed from the genome of ArsP *E. coli* by Flp-mediated marker excision. Briefly, electrocompetent ArsP *E. coli* cells were transformed with pFLP3, plated on LB-Amp agar and positive single colonies patched onto both LB-Amp and LB-Kan agar plates. The Kan-sensitive and Amp-resistant single colonies were selected and further streaked onto LB+5% sucrose plates (to induce pFLP3 loss) and incubated at 37°C until sucrose-resistant colonies appeared. Then the sucrose-resistant colonies were patched on LB-Amp, LB-Kan, and LB+5% sucrose agar plates separately and only sucrose-resistant colonies were selected and re-streaked several times on LB agar until a single colony was finally picked. Insertion of the *arsP* gene into the chromosome of strain MG1655 was confirmed by PCR with genome-specific primers glmS_down and pstS_R (5'- GGAAGAACCGATACCCTG), followed by sequencing of the PCR product.

**Text SM5: Validation of** **MMAs(III)-resistance in ArsP *E. coli*.** To confirm the MMAs(III)-resistance in the constructed ArsP-expressing *E. coli* (hereafter ArsP *E. coli*) and test the toxicity of MMAs(III) to WT *E. coli*, anaerobic microcosms were established in 120 mL serum bottles containing 50 mL anaerobic 1/4 tryptic soy broth (TSB) medium. We have tested the stability of MMAs(III) in various media, including 100%, 75%, 50%, and 25% RCB, 100%, 75%, 50%, and 25% TSB, LB, the spent RCB (in which strain EML had grown), and 2 x YTG (yeast extract-tryptone-glucose) medium, and found that MMAs(III) is most stable in 25% TSB medium than the rest of tested media (data not shown). Thus, 1/4 TSB medium is selected for MMAs(III) toxicity test. Thus, pre-cultures of WT *E. coli* and ArsP *E. coli* were grown in 1/4 TSB anaerobically to mid-exponential growth phase. Approximately 0.5 mL of WT *E. coli* and ArsP *E. coli* were inoculated separately into 50 mL 1/4 TSB supplied with 3 μM MMAs(III) using sterile N_2_-flushed syringes and needles. A total of four treatments were conducted, including (i) WT *E. coli* + 3 μM MMAs(III), (ii) ArsP *E. coli* + 3 μM MMAs(III), (iii) WT *E. coli*, and (iv) ArsP *E. coli*, which were carried out at 30°C in dark without shaking. At selected time points and for each condition, triplicate bottles were sampled and growth was measured as optical density at 600 nm (OD_600_) in a spectrophotometer. In addition, aqueous As species was also quantified by HPLC-ICP-MS to monitor the transformation of MMAs(III).

**Text SM6: The toxicity of 1 μM MMAs(III) to WT *E. coli*.** According to anaerobic As methylation by strain EML, it could produce approximately 0.8 μM MMAs(III) during the exponential growth phase in 100% RCB supplied with 25 μM As(III) (Figure S9). Thus, the toxicity of 1 μM of MMAs(III) to WT *E. coli* was also tested to mimic *in vivo* MMAs(III) concentration as described above. Two treatments, (i) WT *E. coli* + 1 μM MMAs(III), (ii) ArsP *E. coli* + 1 μM MMAs(III) were conducted at 30°C in dark without shaking. At selected time points and for each condition, triplicate bottles were sampled and growth was measured were determined by measuring the optical density at 600 nm (OD_600_).

**Text SM7: Optimization of the inoculation ratio of co-culture strains.** Given the distinct growth rates of strain EML and *E. coli*, we needed to ensure that strain EML was sufficiently abundant in the co-culture system to produce at least 1 μM MMAs(III), otherwise *E. coli* would dominate the co-culture system. To do so we first inoculated 0.5 mL of a pre-culture of strain EML at exponential growth phase into 50 mL 100% RCB supplied with 25 μM As(III), allowing it to grow for 6 h to reach its mid-exponential growth phase. Then either 0.3 mL or 0.03 mL pre-culture of WT *E. coli* or ArsP *E. coli* (exponential growth phase) was added back into the system. During the co-culture incubation, 2 mL of culture from duplicate bottles were collected using sterile, N_2_-flushed syringes and needles at 0, 4, 8, and 12 hours. Cells were pelleted by centrifugation at 8000 g for 10 min and DNA was extracted by the DNeasy PowerSoil Pro Kit (Qiagen). The growth rates of WT *E. coli* and ArsP *E. coli*, and strain EML were determined using qPCR by quantifying the 16S rRNA gene copy number of *E. coli*. The specific primer sets targeting the 16S rRNA genes from *E. coli* and strain EML were designed by Benchling online research platform as described above. The qPCR primer set for 16S rRNA gene from *E. coli* is: E-16S-F: CTAGGCGACGATCCCTAGCTGG and E-16S-R: GCCTTCTTCATACACGCGGCAT. The primer targeting 16S rRNA gene from strain EML is: EML-16S-F: ACTCTTGCGAGCGTACTTCCCA and EML-16S-R: GGCGGCTCTCTGGACTGTAACT. The qPCR was carried out in a Mic PCR system (Bio Molecular Systems, Mic) using SYBR Green Master Mix. The reactions (10 μL total volume), contained 5 μL of 2 × SensiFAST™ SYBR No-ROX Kit (Bioline, London, UK), 0.2 μM of each *E. coli 16S rRNA* gene primer, 2.5 μL of DNA, and 1% (v/v) bovine serum albumin (BSA) (Sigma). The thermal cycling conditions were: 95°C 5 min, followed by 50 cycles of 95°C 5 s, 60°C 20 s and 72°C 20 s. A 10-fold dilution series containing 10^7^-10^1^ copies of *E. coli* plasmid DNA was used to generate a standard curve. The *E. coli* and strain EML 16S rRNA gene plasmid standards were constructed by the TA-Cloning method as described above.

In addition to first growing strain EML to its exponential phase, then adding *E. coli* into the co-culture system. We also estimated various inoculum ratios of *E. coli* strains and strain EML, i.e., 1:2, 1:4, 1:8, 1:16, and 1:32 (v/v) in duplicate in 50 mL anoxic 100% RCB supplied with 25 μM As(III). The inoculum size of *E. coli* is fixed (0.05 mL of pre-culture of *E. coli* at exponential growth phase). At selected points and for each condition, bacterial cultures were collected and subject to DNA extraction, and further qPCR was used to determine the growth rates of WT or ArsP *E. coli*, and strain EML as described above.

Rather than directly mixing strain EML with *E. coli*, we further tried to pellet various concentrations of strain EML pre-cultures, then mixed with *E. coli*, and added into co-culture system in 50 mL anoxic 100% RCB supplied with 25 μM As(III). The inoculum size of *E. coli* was set at 0.05 mL and various strain EML cell concentrations were prepared, i.e., 10%, 20%, 30%, and 40%, which are equivalent to the cell pellets of 5, 10, 15, and 20 mL of strain EML pre-culture, respectively. At 0, 4, 8, and 12 h, the co-culture was sampled and DNA was extracted for qPCR quantification of the growth rates of WT *E. coli* and ArsP *E. coli*, and strain EML during anaerobic co-culture incubation.

**Supporting Results:**

**Text SR1: As(III)-resistance in *E. coli*.** Anaerobic growth of WT *E. coli* and ArsP *E. coli* was conducted in 100% RCB supplied with 10, 15, and 25 μM As(III). As illustrated in Figure S3 and Table S15, both *E. coli* strains exhibited similar growth patterns under different As(III) concentrations, i.e., grew rapidly during 20 hours of incubation, then reached a plateau after 24 hours. This result demonstrates that the presence of As(III) did not affect the growth of WT *E. coli* and ArsP *E. coli* in RCB.

**Text SR2: MMAs(III)-resistance in ArsP-expressing *E. coli*.** Anaerobic growth of WT *E. coli* and ArsP *E. coli* was carried out in 1/4 TSB supplied with 3 μM MMAs(III). The growth curves evidenced distinct growth dynamics of WT and ArsP *E. coli* in the presence of MMAs(III) (Figure S1 and Table S16). The growth of WT *E. coli* was completely inhibited by MMAs(III) while that of ArsP *E. coli* was only slightly affected compared with the no-MMAs(III) controls (Figure S1a). As speciation analysis show that MMAs(III) remained quite stable in the medium with WT *E. coli* (Figure S1b), while when incubating with ArsP *E. coli*, the concentration of MMAs(III) started to decline after 24 hours of incubation, with approximately 1.8 μM MMAs(III) retained at the end of incubation (48 hours) (Figure S1c). The decrease in MMAs(III) occurred during the stationary growth phase and we presume that most of MMAs(III) may be trapped inside the cells due to the lower MMAs(III) extrusion efficiency in the stationary cells than the exponential ones. Besides MMAs(III), a small proportion of MMAs(V) (< 0.3 μM) was also detected and this could be due to abiotic oxidation in the medium or analysis artifacts resulting from sample preservation and storage. To simulate *in vivo* MMAs(III) production by strain EML, the WT *E. coli* and ArsP *E. coli* were further inoculated in anoxic 1/4 TSB with 1 μM MMAs(III) as described above (Figure S2 and Table S17). Similarly, the presence of 1 μM MMAs(III) inhibited the growth of WT *E. coli* but to lesser extent than 3 μM MMAs(III) (Figures S1 and S2). While the growth of ArsP *E. coli* was barely affected by MMAs(III) relative to that of the no-MMAs(III) controls (Figure S2). Taken together, these data show that the recombinant *arsP* gene in *E. coli* can be functionally expressed under anoxic condition, conferring resistance to MMAs(III). In addition, the presence of 1 MMAs(III) is high enough to inhibit the growth of the MMAs(III)-sensitive strain.

**Text SR3: Optimization of the inoculation ratio of co-culture strains.** Different combinations of strain EML and the *E. coli* strains (either WT or ArsP *E. coli*) were compared in 100% RCB with 25 μM As(III) to optimize the inoculation ratio for the two co-culture members. The first attempt was: growing strain EML to its exponential growth phase, then adding 0.3 mL of WT *E. coli* or ArsP *E. coli* into the co-culture system. The growth curves show that either *E. coli* strain barely grew after 20 hours of incubation (Figure S4a and Table S18a). This could be attributed to either the initial inoculum size of *E. coli*, that might be too large so that it grew too fast to detect the growth dynamics, or the depletion of growth nutrients in the medium. The second attempt was: growing strain EML to its exponential growth phase, then added 0.03 mL of either WT *E. coli* or ArsP *E. coli* into the co-culture system. Similarly, neither *E. coli* strain exhibited obvious growth (Figure S4b and Table S18b) and we assume that nutrients were depleted. The third attempt was: growing strain EML and either *E. coli* strain simultaneously with various inoculation ratios. We found that both *E. coli* strains showed similar growth patterns when co-culturing with strain EML under the ratios of *E. coli* and strain EML = 1:4, 1:8, 1:16, and 1:32 (Figure S5 and Table S19). Compared with the ratios of 1:16 and 1:32, the growth of *E. coli* and strain EML was more obvious at the ratios of 1:2, 1:4, and 1:8 (Figure S5). The fourth attempt was: mixing the various concentrated strain EML cells with both *E. coli* strains (individually) instead of adding strain EML first to the medium, i.e., 10%, 20%, 30%, and 40% (v/v) strain EML + either WT *E. coli* or ArsP *E. coli.* The growth of ArsP *E. coli* was much higher than that of WT *E. coli* while strain EML showed similar growth rates in the 10% and 40% strain EML with *E. coli* treatments (Figure S6 and Table S20). As speciation analysis demonstrated that MMAs(III) was the dominant methylated As species, with 1.47 and 1.57 μM aqueous MMAs(III) detected in 10% EML + either WT *E. coli* or ArsP *E. coli*, respectively (Figure S6b). The concentration of MMAs(III) increased along with the cell density of strain EML. For instance, in the 40% strain EML + WT *E. coli* or ArsP *E. coli*, 3.37 and 3.19 μM aqueous MMAs(III) were detected, respectively (Figure S6b). Therefore, an optimal ratio for the co-culture system was determined, i.e., 10% strain EML cell pellet with 0.05 mL *E. coli* cell culture to investigate the microbial warfare hypothesis for anaerobic As methylation.

**Text SR4: Chemical transformation of MMAs(III) in biological media.** In the oxidized samples, a low concentration of DMAs(V) (Figure S9e-f and Table S5) and an unknown As peak (Figure S19) were also detected. We propose that a thiolated and/or methylated species may be generated and retained in the anion-exchange column. However, upon reaction with H_2_O_2_, it is partially oxidized to pentavalent methylated As species (DMAs(V)) and readily eluted and separated. The species corresponding to the abovementioned unknown peak has not been identified. As mass balance was evaluated by the sum of all As species detected (using HPLC-ICP-MS) divided by the sum of total As measured (using ICP-MS) (Figure S20 and Table S24). The overall mass balance for As in our study is around 74%-118% (no oxidation) and 89%-150% (oxidation), suggesting that chemical oxidation of the sample recovers more of the As chemical species, albeit in altered form.

FIGURES

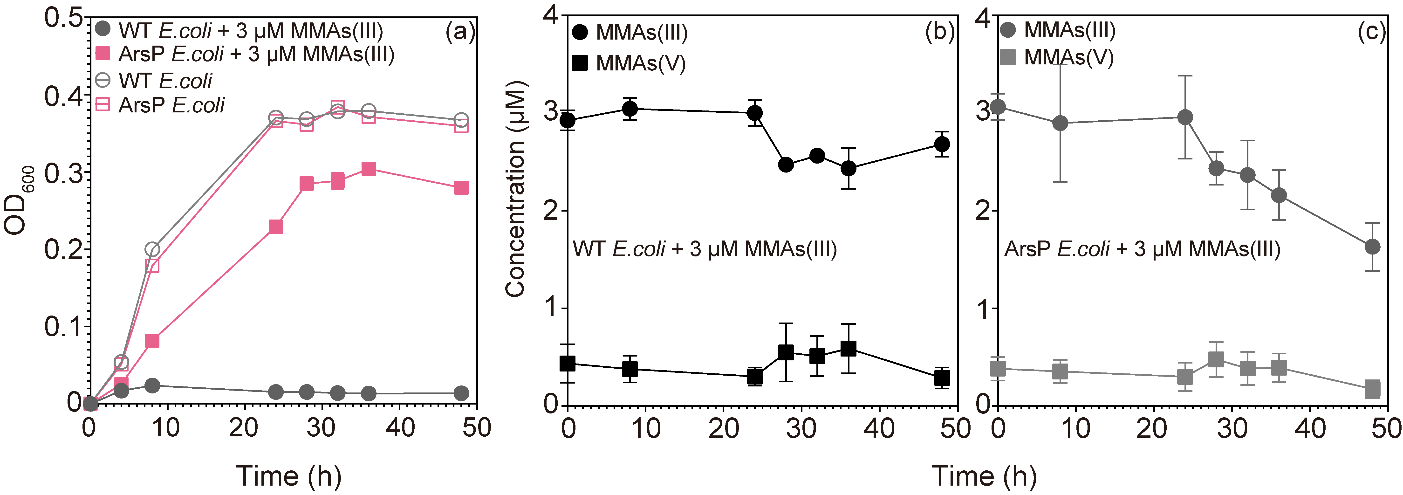

**Figure S1.** (a) Growth curves (OD_600_) of *Escherichia coli* K-12 wild-type strain MG1655 (WT *E. coli*) and engineered WT *E. coli* harboring an expressing MMAs(III)-resistance gene (*arsP*) (ArsP *E. coli*) in anoxic 1/4 tryptic soy broth (TSB) medium in the presence or absence of 3 μM MMAs(III). (b) and (c) Time-dependent stability of aqueous MMAs(III) in 1/4 TSB incubating with WT *E. coli* and ArsP *E. coli*. Data are shown as mean values with error bars. Individual values for each biological triplicate are included in Supporting Information Table S16.

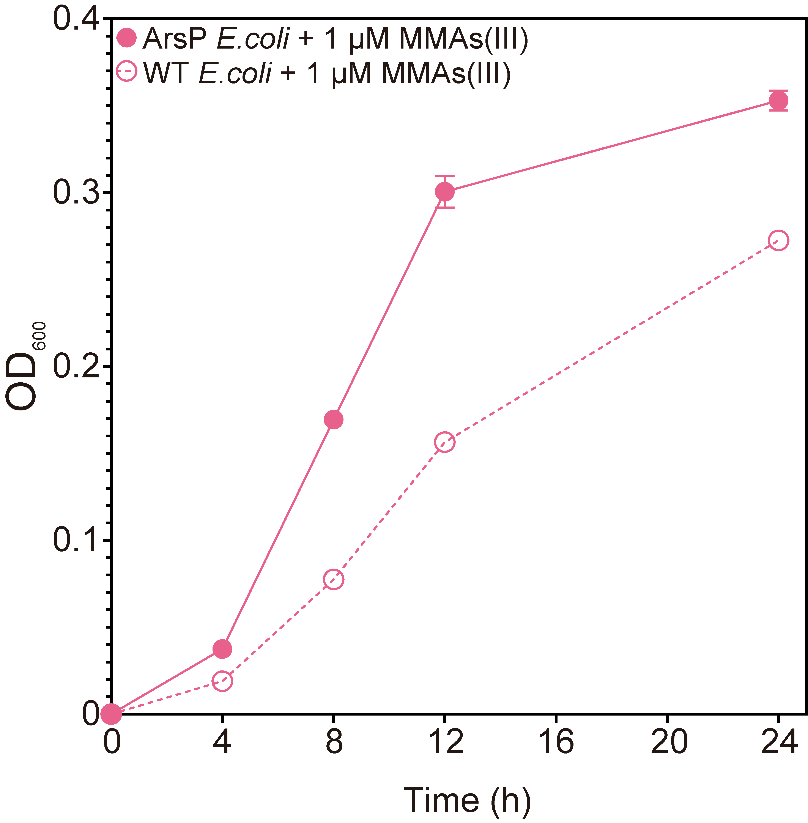

**Figure S2.** Growth curves (OD_600_) of WT *E. coli* and ArsP *E. coli* in anoxic 1/4 TSB in the presence or absence of 1 μM MMAs(III). Data are shown as mean values with error bars. Individual values for each biological triplicate are included in Supporting Information Table S17.

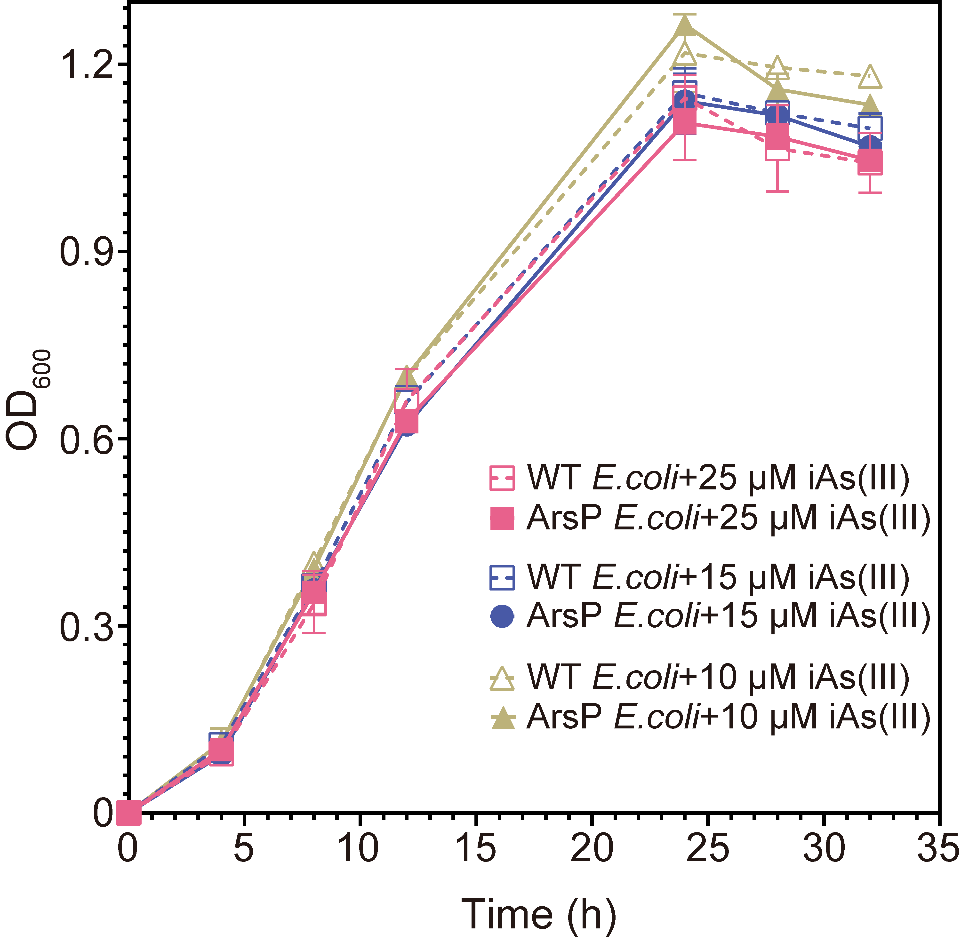

**Figure S3.** Growth curves (OD_600_) of WT *E. coli* and ArsP *E. coli* in anoxic RCB in the presence or absence of 10, 15, or 25 μM iAs(III). Data are shown as mean values with error bars. Individual values for each biological triplicate are included in Supporting Information Table S15.

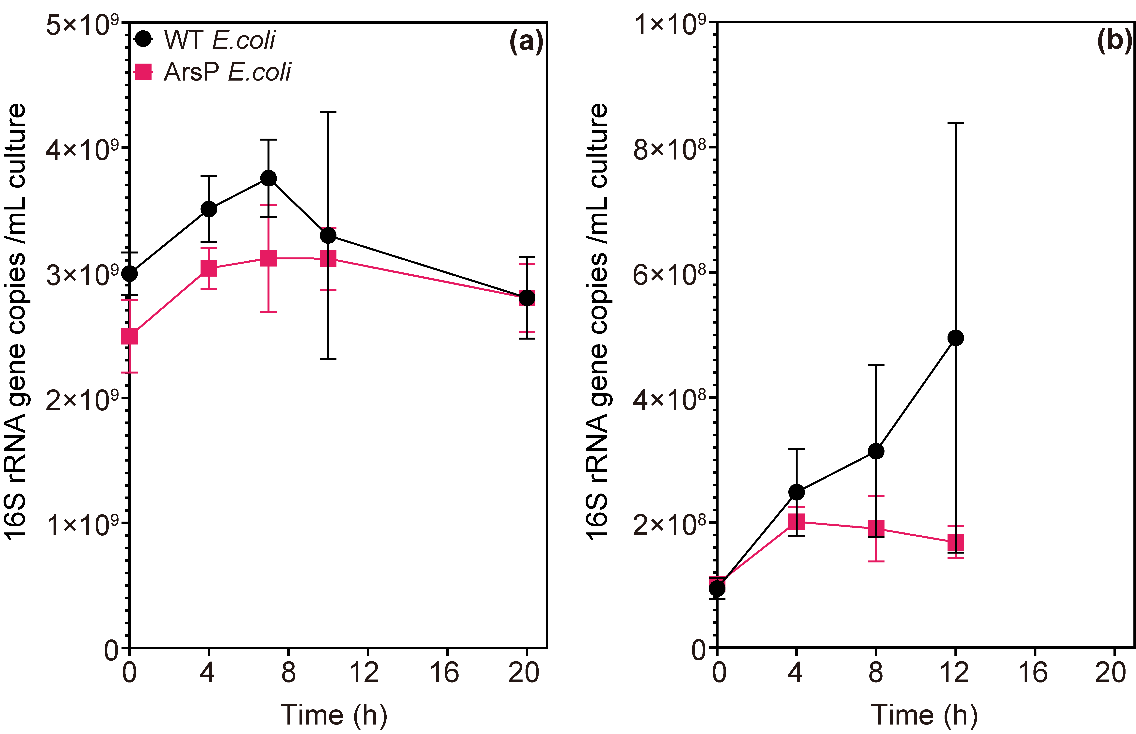

**Figure S4.** Growth curves (16S rRNA gene copy number) of WT *E. coli* and ArsP *E. coli* in anaerobic co-culture with *Paraclostridium bifermentans* strain EML in anoxic RCB supplied with 25 μM iAs(III). The result corresponds to the first and second attempts to optimize inoculation ratio between the co-culture strains in Supplementary Information Text Results SR3. E. coli (either strain) was added only after 6 hours of strain EML growth alone. The two conditions are: (a) 0.5 mL strain EML + 0.3 mL *E. coli*, (b) 0.5 mL strain EML + 0.03 mL *E. coli*. Data are shown as mean values with error bars. Individual values for each biological triplicate are included in Supporting Information Table S18.

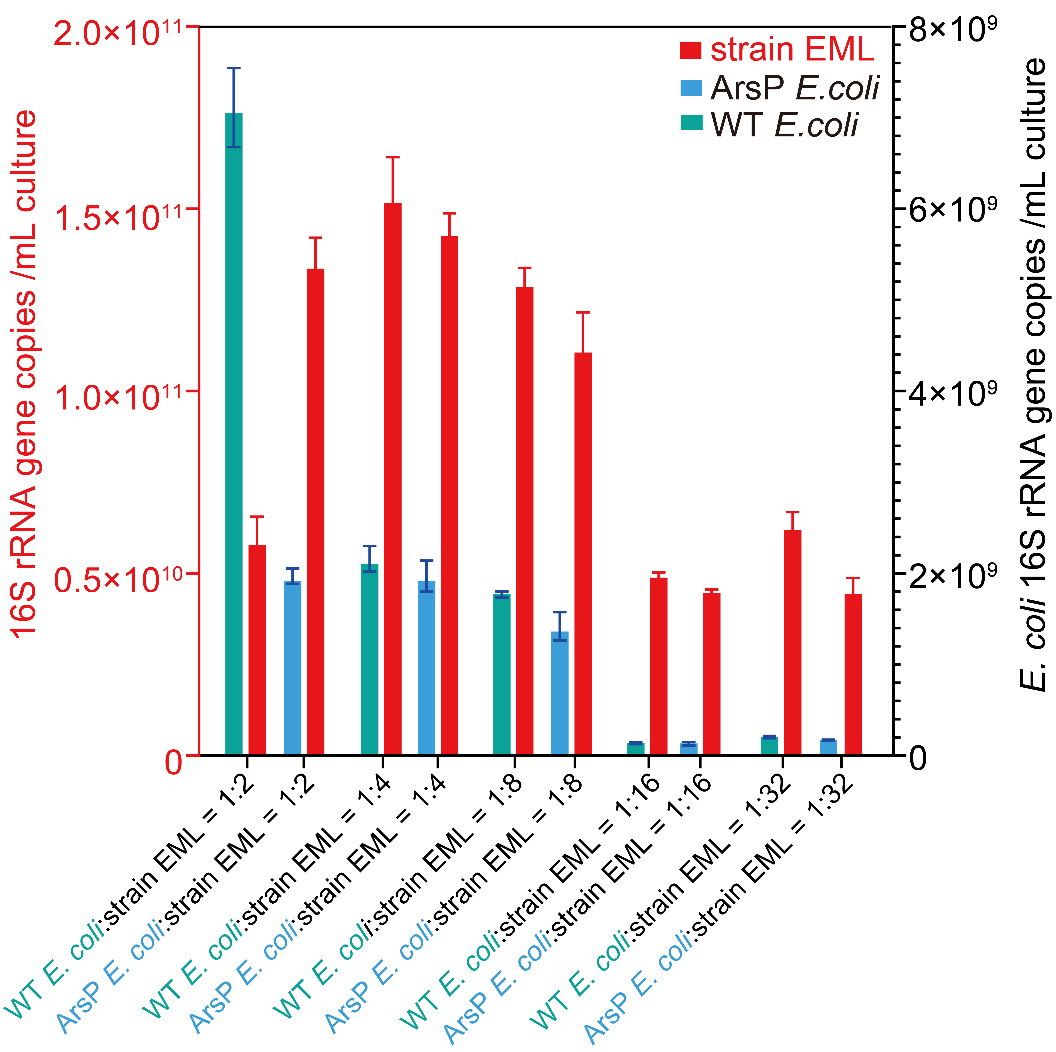

**Figure S5.** Growth (16S rRNA gene copy number) of the co-culture members in anoxic RCB supplied with 25 μM iAs(III) after 12 hours of incubation. The result corresponds to the third attempt to optimize inoculation ratio between the co-culture strains in Supplementary Information Text Results SR3. *Paraclostridium bifermentans* strain EML (red) was co-culture with either WT *E. coli* (green) or ArsP *E. coli* (blue) anaerobically at various inoculation ratios of *E. coli*: strain EML = 1:2, 1:4, 1:8, 1:16, and 1:32 (v/v). Data are shown as mean values with error bars. Individual values for each biological triplicate are included in Supporting Information Table S19.

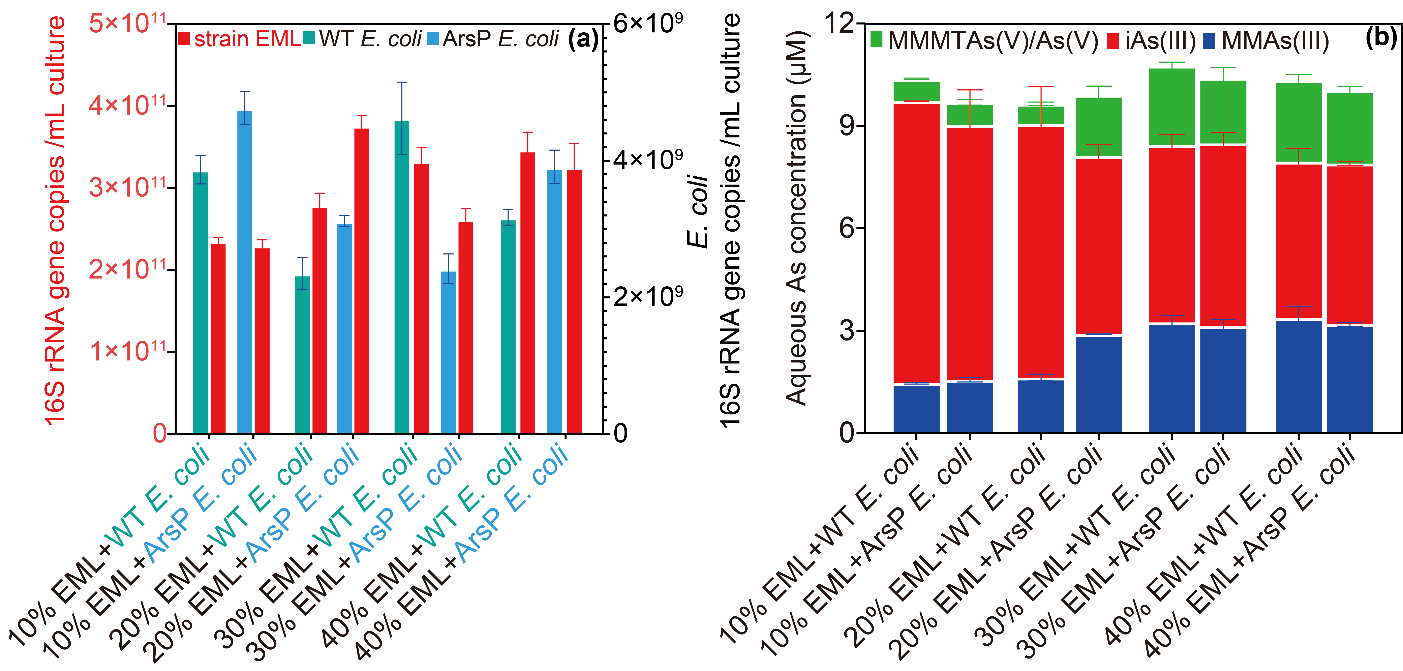

**Figure S6.** (a) Growth (16S rRNA gene copy number) of the co-culture members in anoxic RCB supplied with 25 μM iAs(III) after 12 hours of incubation. The result corresponds to the fourth attempt to optimize inoculation ratio between the co-culture strains in Supplementary Information Text Results SR3. *Paraclostridium bifermentans* strain EML (red) (various volumes of cell culture) was co-cultured with either WT *E. coli* (green) or ArsP *E. coli* (blue) (a 50 μL culture at exponential phase). The amount of strain EML in RCB medium was systematically varied from the cell pellet of a 5 mL culture (10%), 10 mL culture (20%), 15 mL culture (20%), or 20 mL culture (40%). (b) Aqueous As speciation in the above-described co-culture systems. Data are shown as mean values with error bars. Individual values for each biological triplicate are included in Supporting Information Table S20.

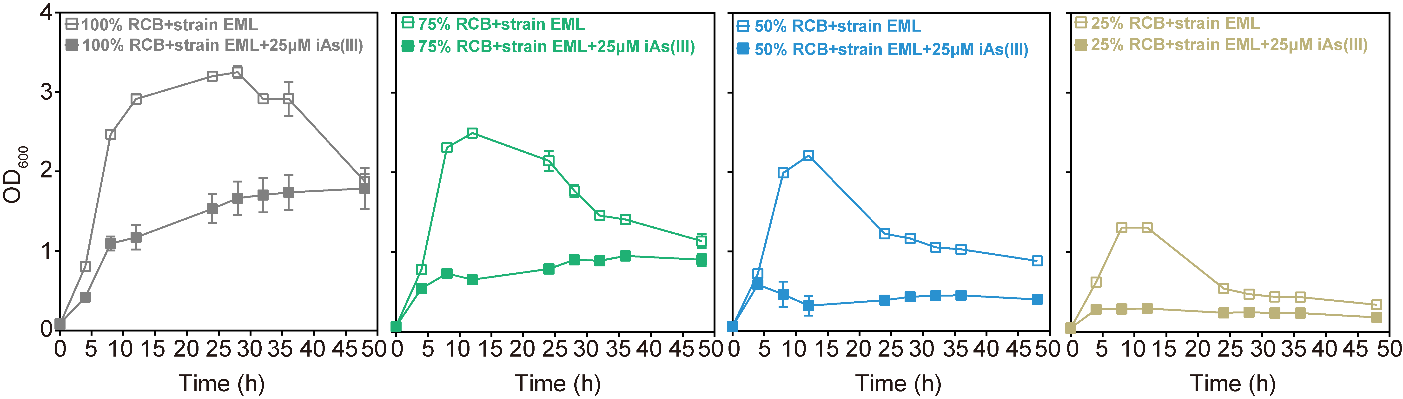

**Figure S7.** Growth curves (OD_600_) of *Paraclostridium bifermentans* strain EML in anaerobic dilutions of Reinforced Clostridial Broth (RCB) (100%, 75%, 50%, or 25% RCB) in the presence or absence of 25 μM iAs(III). Data are shown as mean values with error bars. Individual values for each biological triplicate are included in Supporting Information Table S3.

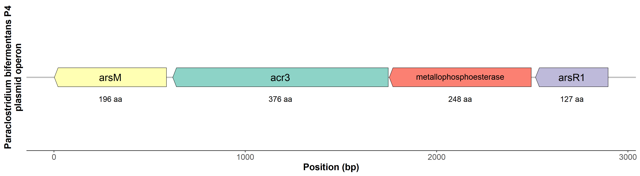

**Figure S8.** Genetic organization of the plasmid-encoded *ars* operon from *Paraclostridium bifermentans* strain EML. Arrows represent open reading frames and orientation of transcription.

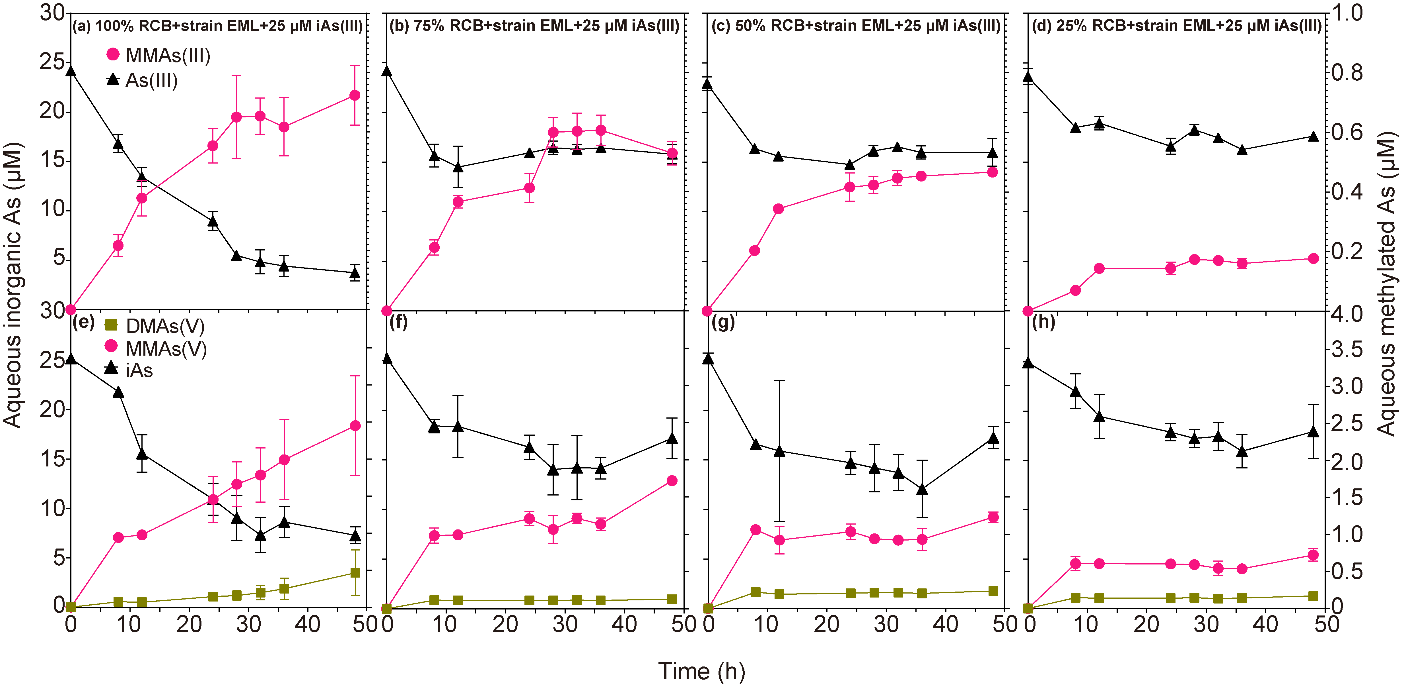

**Figure S9.** Time-dependent concentrations of aqueous As species produced by *Paraclostridium bifermentans* strain EML in anaerobic RCB dilutions (100%, 75%, 50%, or 25% RCB) in the presence of 25 μM iAs(III). (a)-(d), aqueous As species determined by HPLC-ICP-MS. (e)-(f), aqueous As species determined by HPLC-ICP-MS post-oxidation with H_2_O_2_. Individual values for each biological replicate are listed in Supporting Information Table S5.

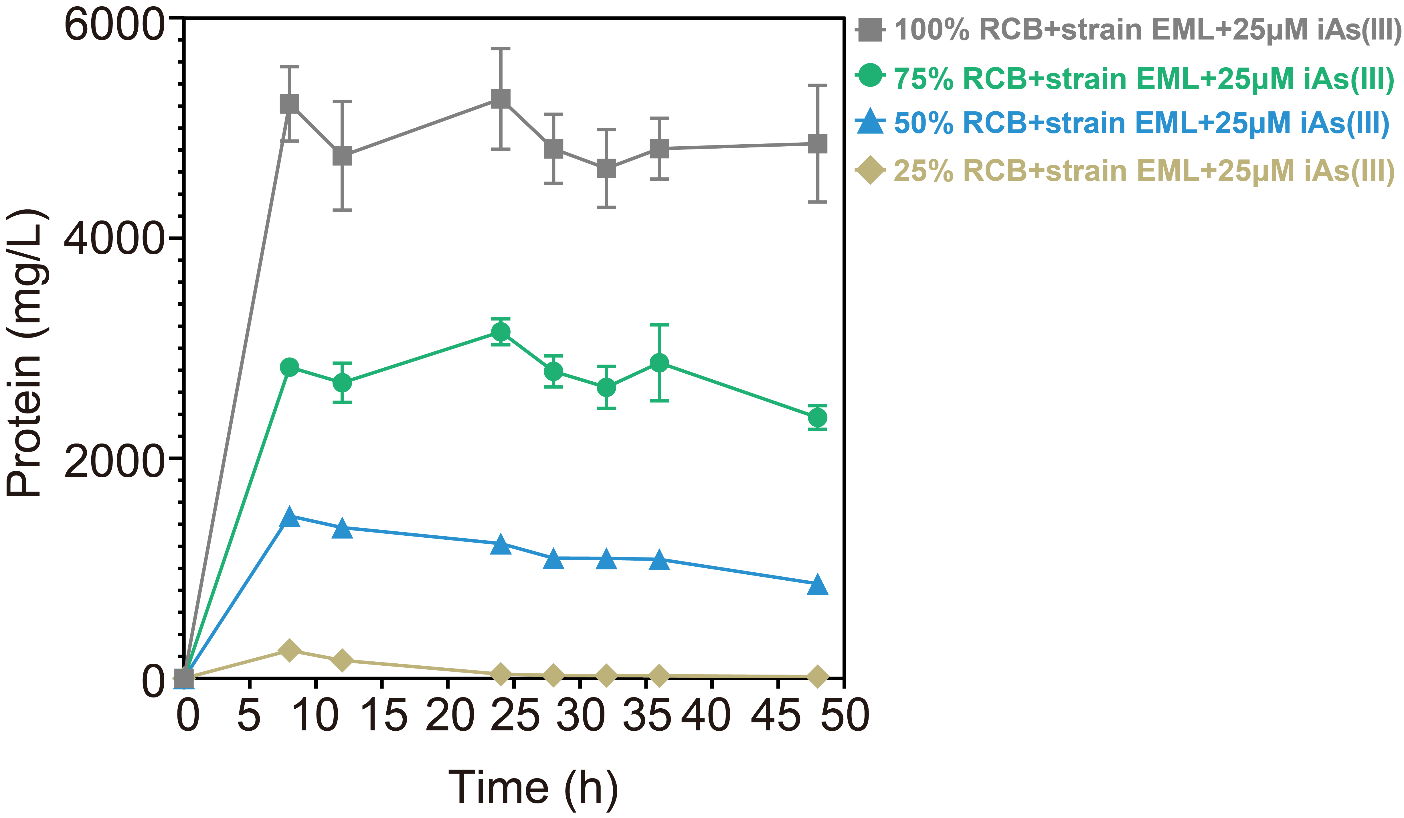

**Figure S10.** Growth curves (total protein) of *Paraclostridium bifermentans* strain EML in anaerobic RCB dilutions (100%, 75%, 50%, or 25% RCB) in the presence of 25 μM iAs(III). Data are shown as mean values of triplicate cultures with error bars. Individual values for each biological replicate are included in Supporting Information Table S4.

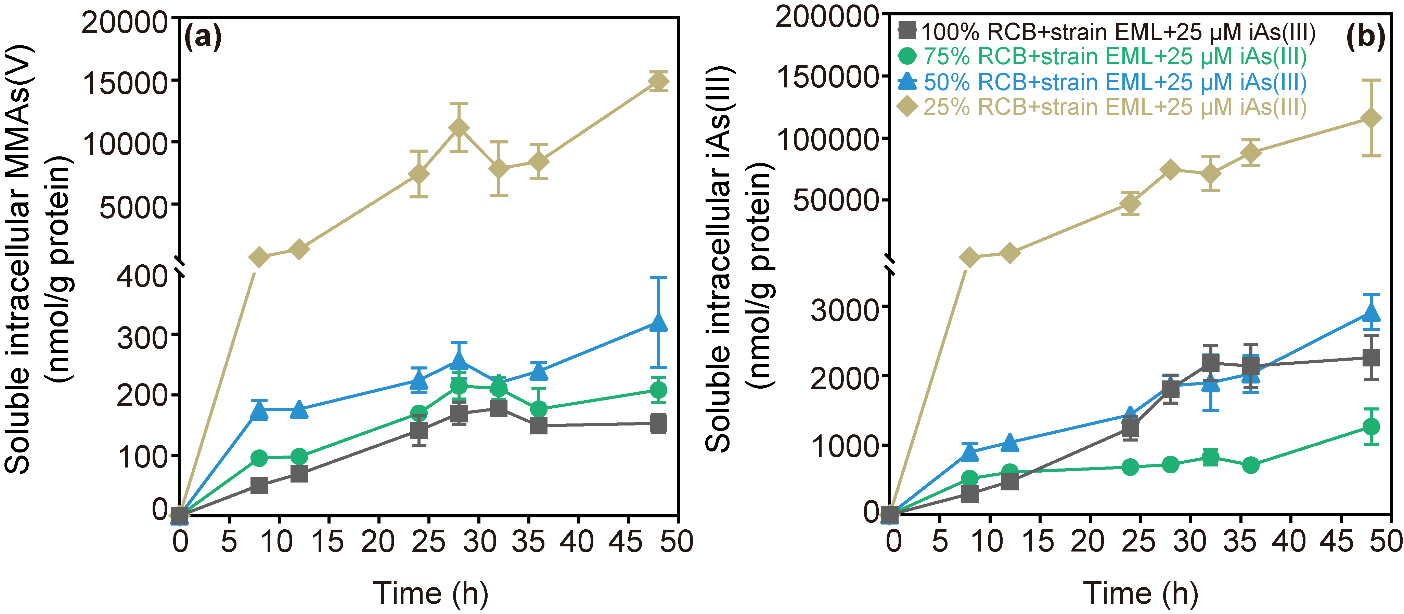

**Figure S11.** Time-dependent concentration of normalized soluble intracellular MMAs(V) (a) and intracellular iAs(III) (b) in anoxic RCB dilutions (100%, 75%, 50%, or 25% RCB) inoculated with *Paraclostridium bifermentans* strain EML and amended with 25 μM iAs(III). Individual values for each biological replicate can be found in Supporting Information Tables S8 and S9.

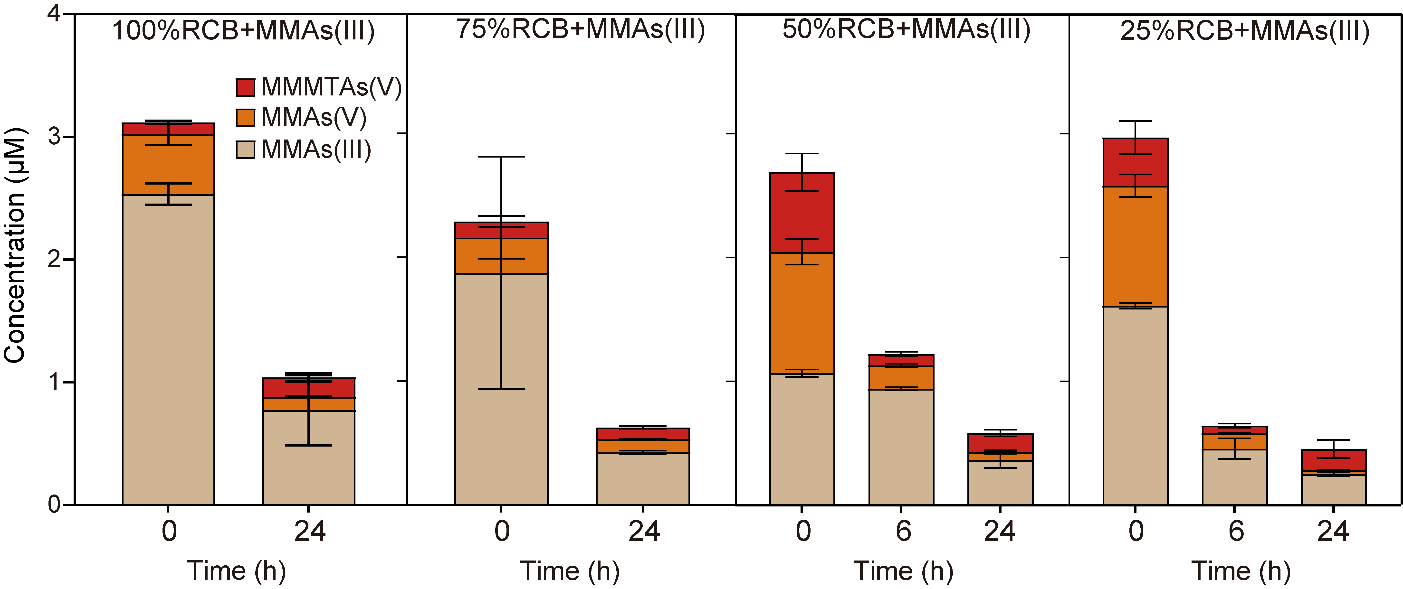

**Figure S12.** Time-dependent stability of aqueous MMAs(III) (3 μM) in anoxic RCB dilutions (100%, 75%, 50%, or 25% RCB). Individual values for each replicate are listed in Supporting Information Table S10.

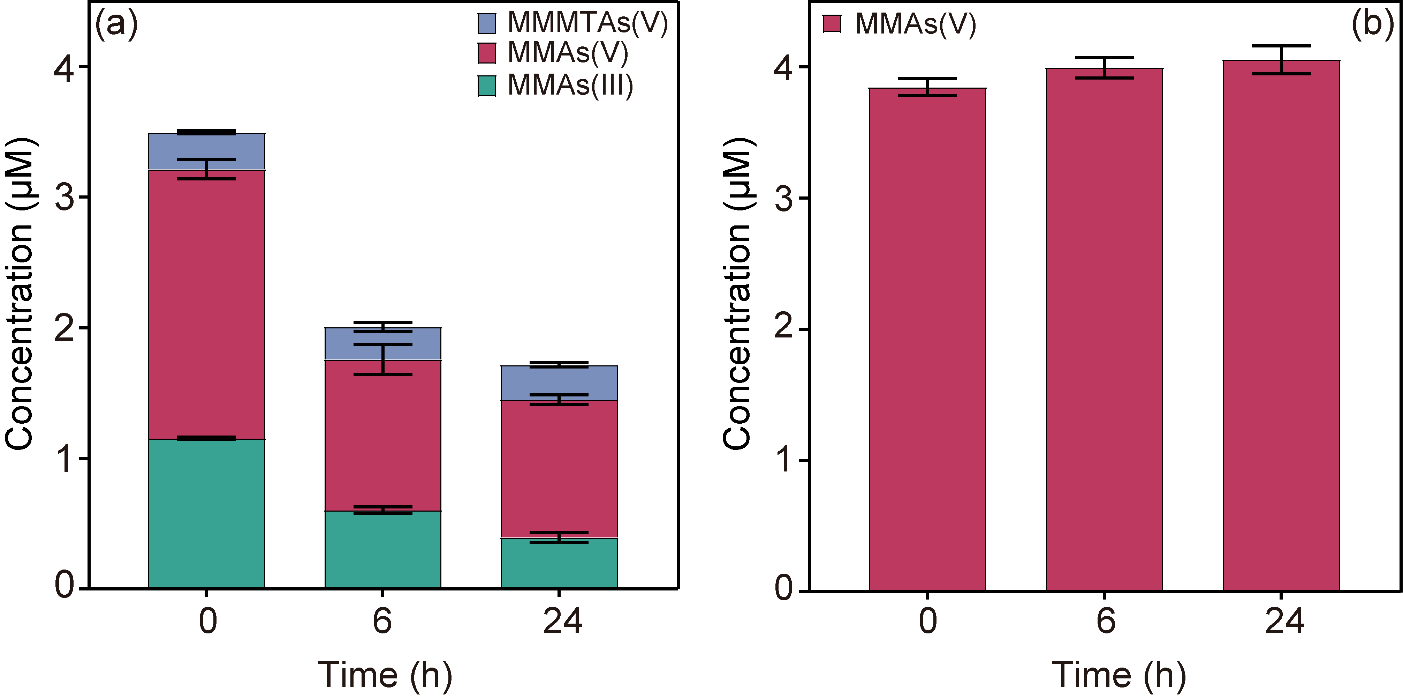

**Figure S13.** Time-dependent stability of aqueous MMAs(III) (3 μM) in anoxic spent RCB (100% RCB). (a) As species without oxidation, and (b) As species post-oxidation. Individual values for each replicate are listed in Supporting Information Table S11.

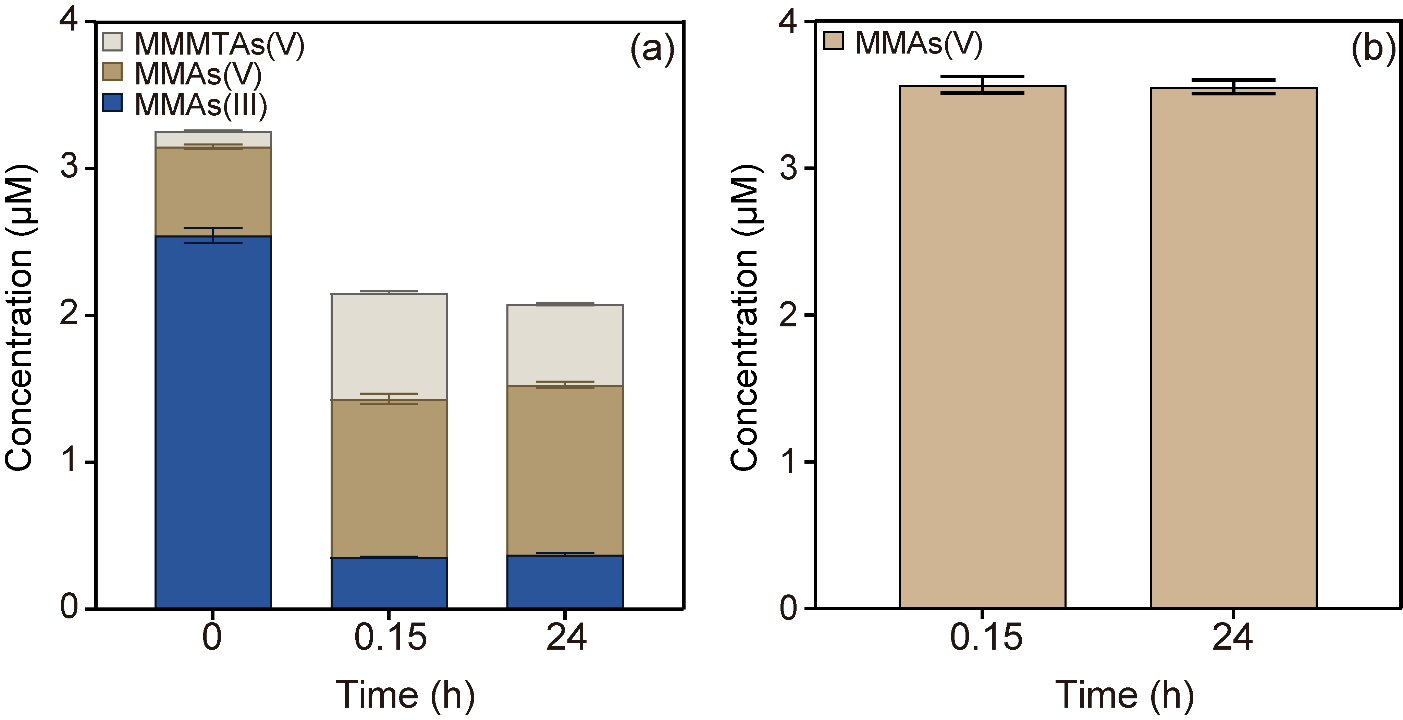

**Figure S14.** Time-dependent chemical reaction between MMAs(III) and sulfide in anoxic water solution. (a) As species without oxidation, and (b) As species with post-oxidation. Individual values for each replicate are listed in Supporting Information Table S12.

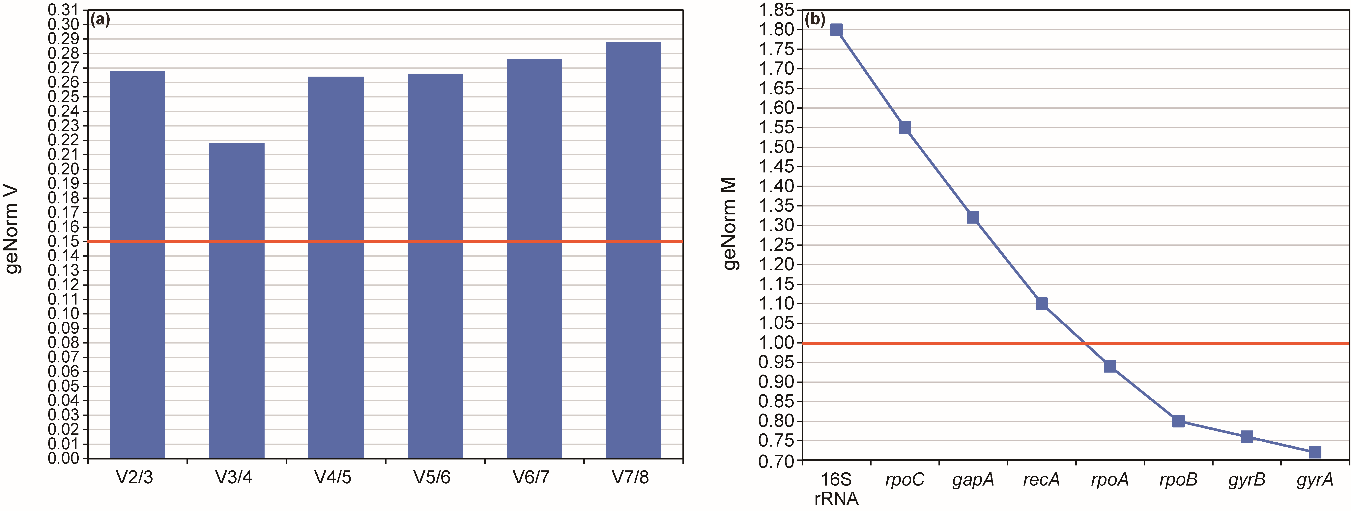

**Figure S15.** (a) The selected minimum number of reference genes based on the average pairwise variation (V) of normalization factors and (b) the calculated expression stability of the candidate reference genes using the geNorm program within qBase plus software. The red line indicates the minimum thresholds (geNorm V (0.15) and geNorm M (1.0)) required for reference genes. None of the genes fulfill this requirement.

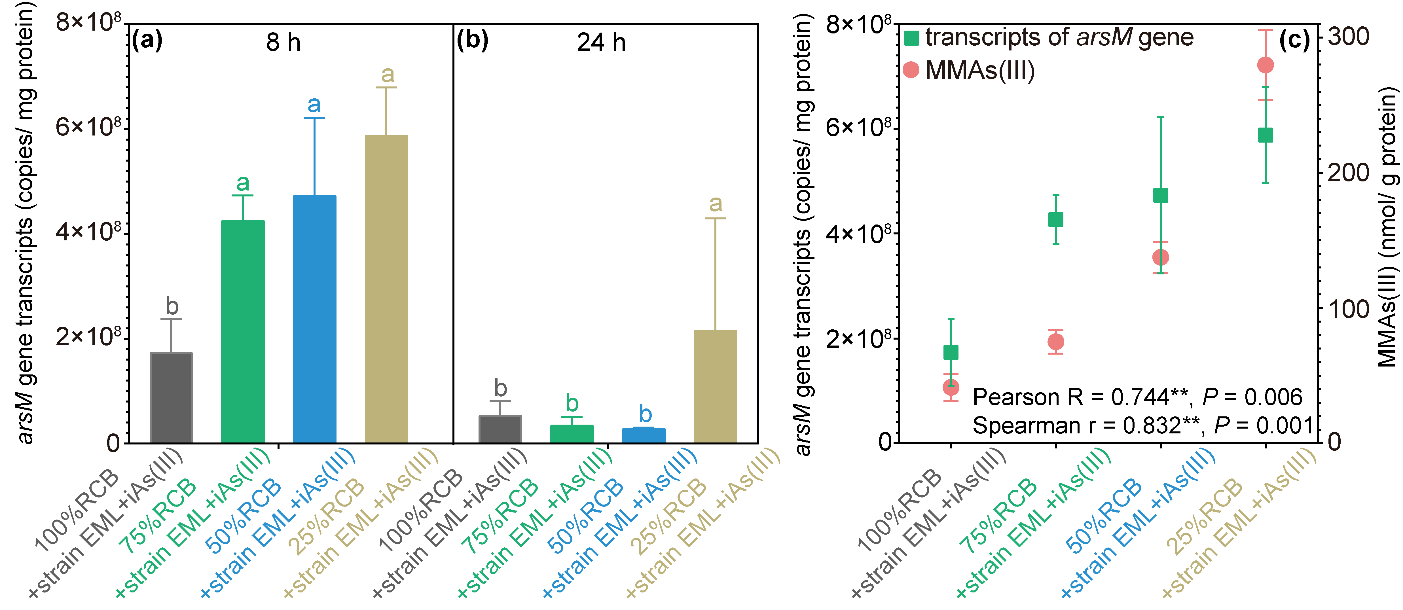

**Figure S16.** Transcript numbers of *arsM* gene of *Paraclostridium bifermentans* strain EML in anoxic dilutions (100%, 75%, 50%, and 25%) of RCB in the presence of 25 μM iAs(III) at 8 and 24 hours of incubation (a). Correlation analysis of *arsM* transcripts and MMAs(III) concentration at 8 hours of incubation (b). Different letters showed significant difference at *P* < 0.05. Individual values for each biological replicate are shown in Supporting Information Table S14. This figure is different from Figure 2 in that the absolute quantification relies on using the same amount total RNA for reverse transcription regardless of biomass amount used for the RNA extraction.

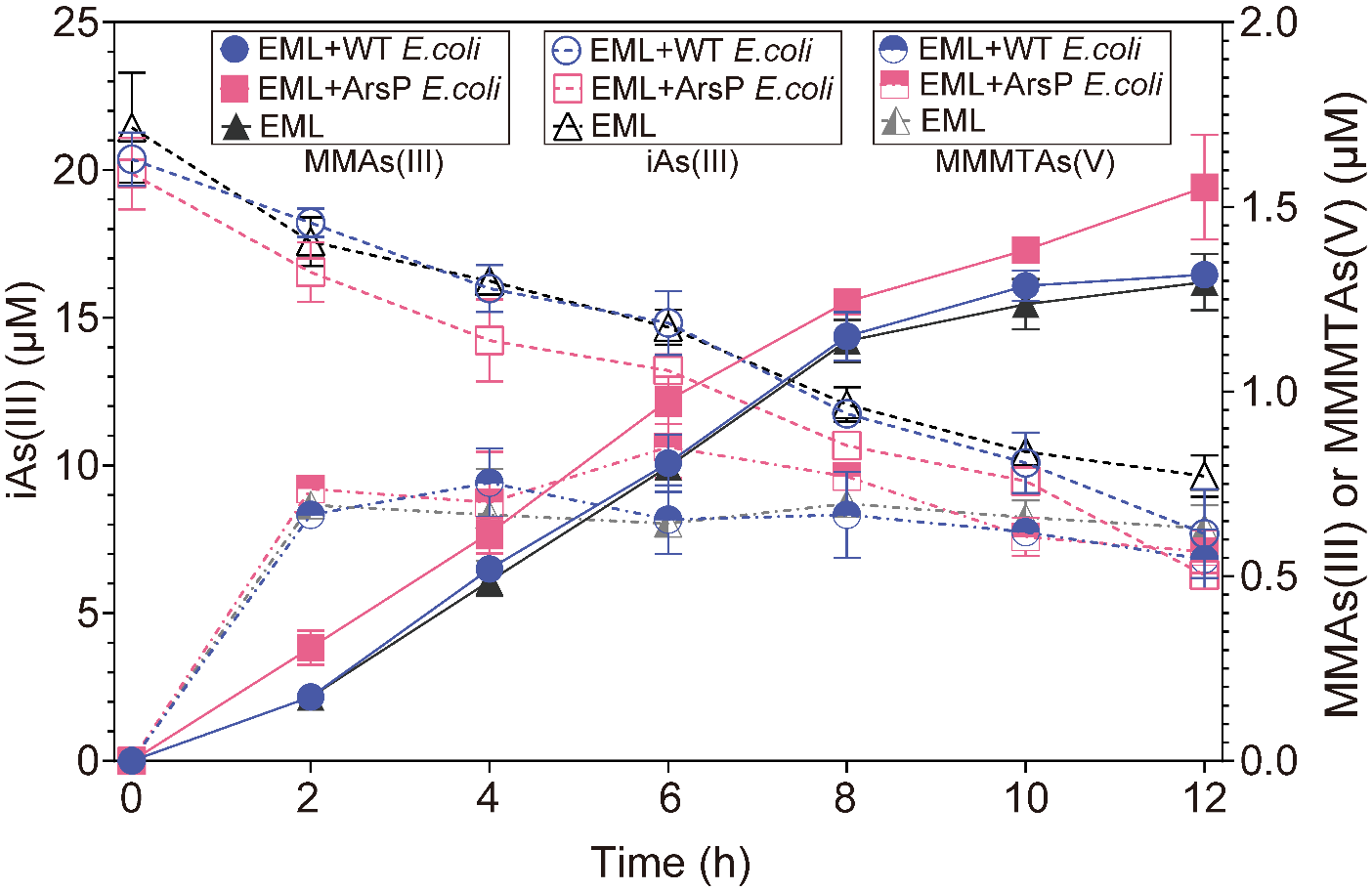

**Figure S17.** Time-dependent concentrations of aqueous As species in anaerobic co-culture *Paraclostridium bifermentans* strain EML with either WT *E. coli* or ArsP *E. coli* in anoxic RCB with 25 μM iAs(III). Individual values for each biological replicate can be found in Supporting Information Table S22.

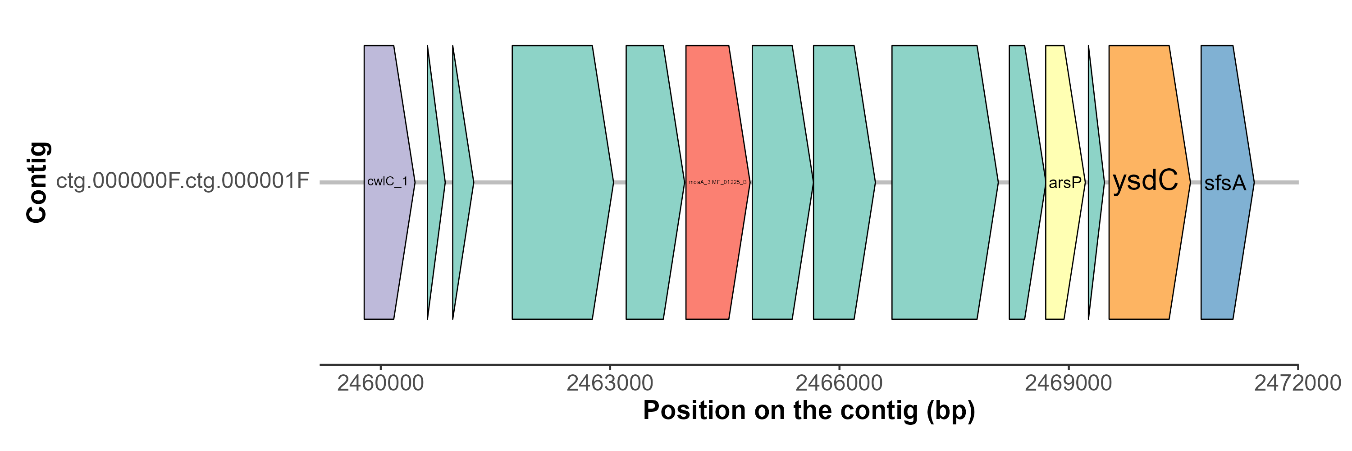

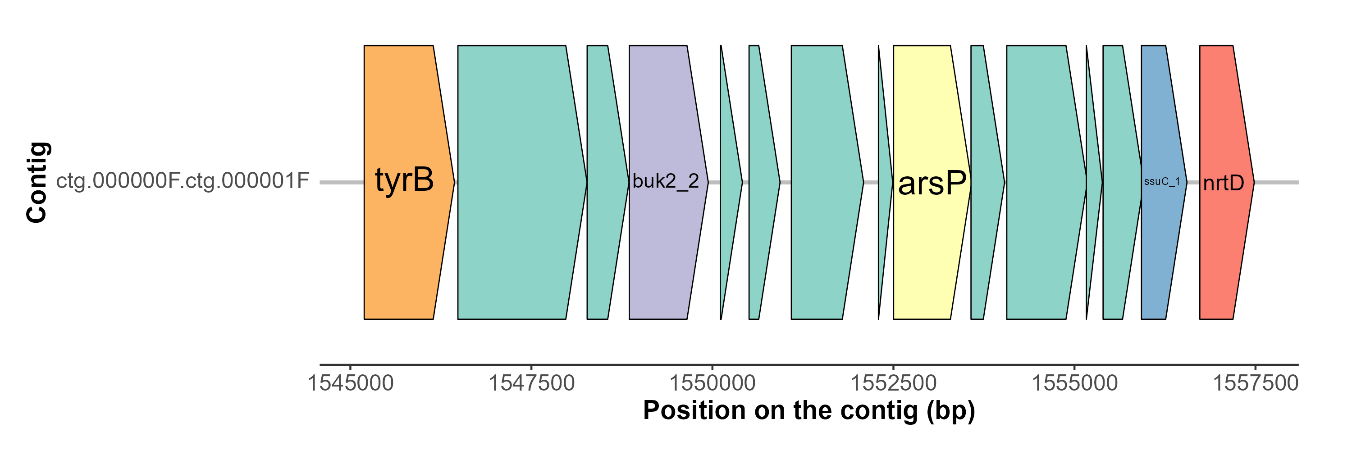

**Figure S18.** The two chromosomally encoded *arsP* genes identified from *Paraclostridium bifermentans* strain EML. Arrow represents open reading frames and orientation of transcription. *ysdC*: Putative aminopeptidase YsdC, *sfsA*: sugar fermentation stimulation protein A, *ssuC*: aliphatic sulfonate ABC transporter membrane subunit, *nrtD*: nitrate/nitrite transport system ATP-binding protein. No other identifiable arsenic-related genes were identified in the vicinity of these two gene.

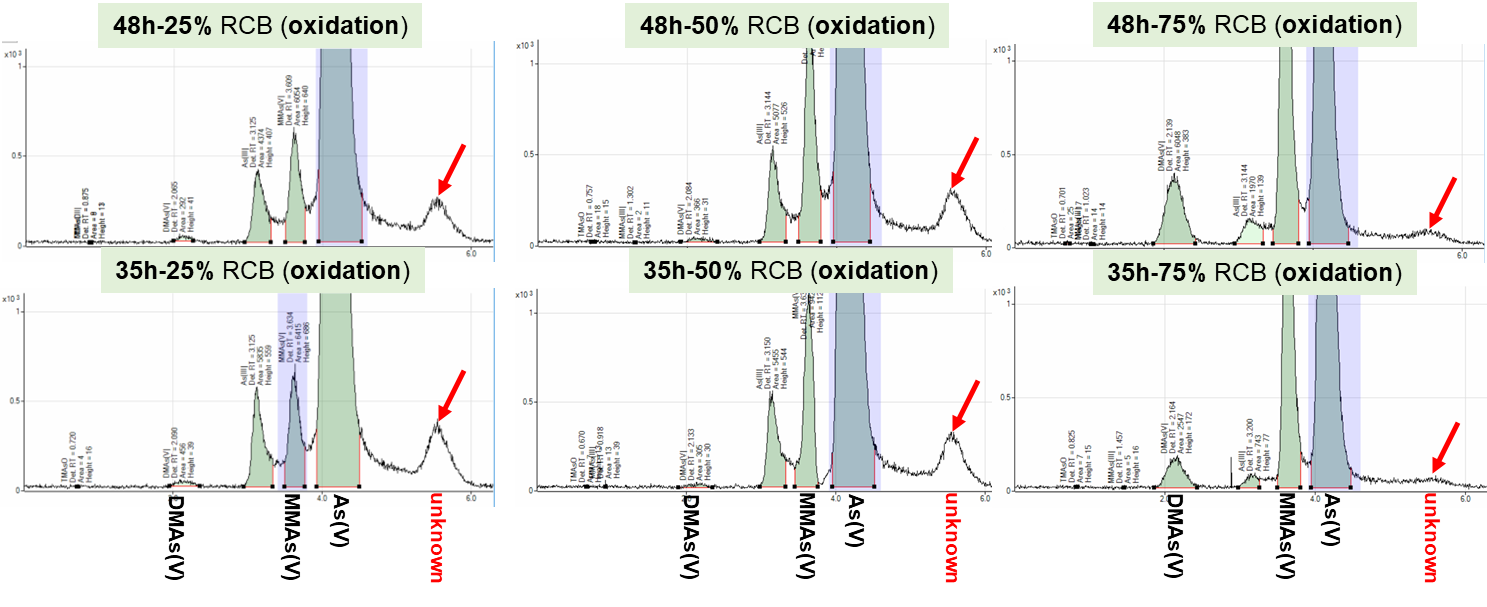

**Figure S19.** The unknown As peak in anaerobic RCB dilutions (75%, 50%, or 25% RCB) inoculated with *Paraclostridium bifermentans* strain EML and 25 μM iAs(III) obtained in oxidized samples at 35 and 48 hours of incubation. The unknown peak was also detected in 100% RCB but was very small compared with other RCB dilutions (not shown).

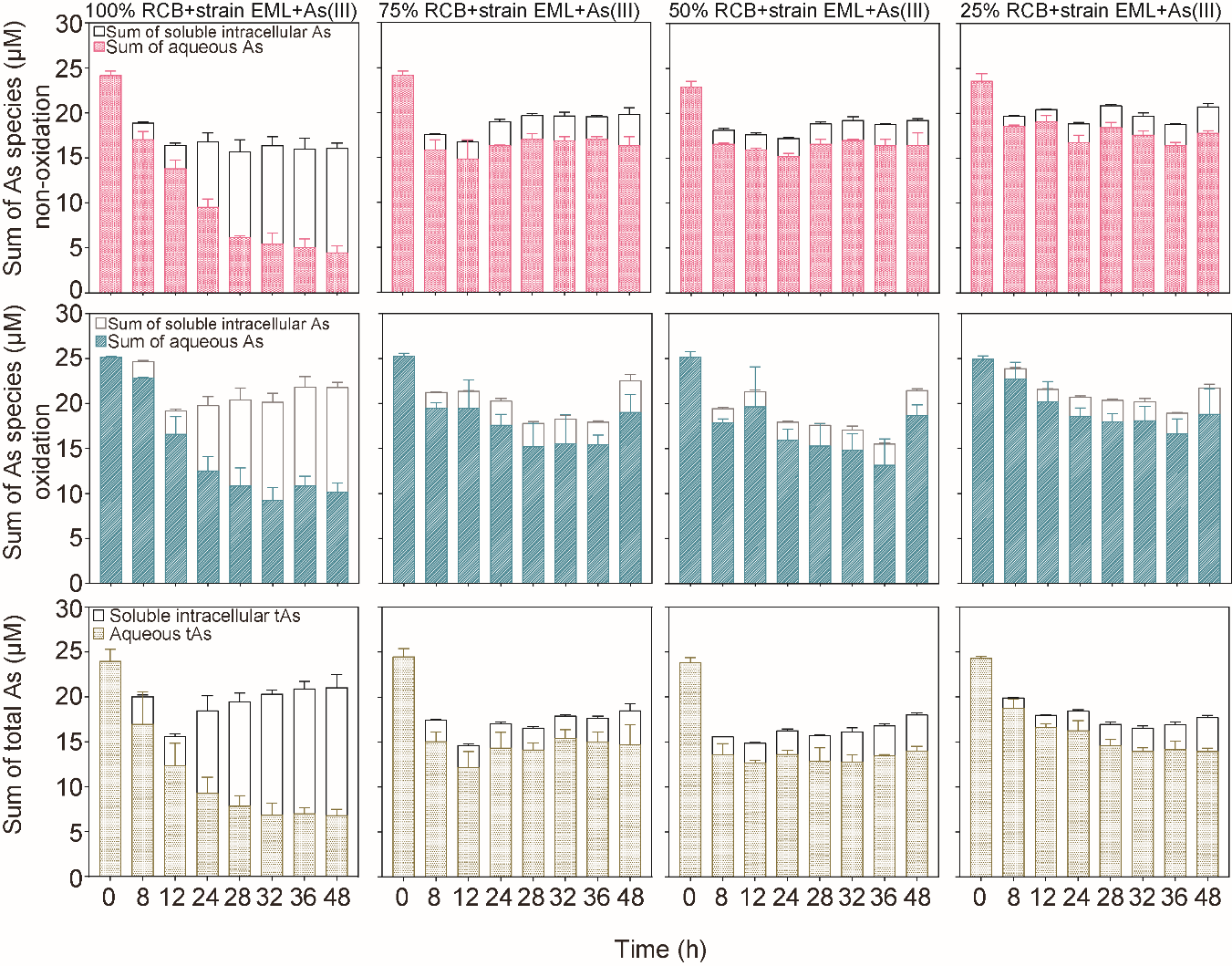

**Figure S20.** Time-dependent comparison of As mass balance in anoxic dilutions (100%, 75%, 50%, and 25%) of RCB inoculated with *Paraclostridium bifermentans* strain EML and 25 μM iAs(III). First row: sum of aqueous As species and soluble intracellular As species (determined by HPLC-ICP-MS). Second row: sum of aqueous As species and soluble intracellular As species (determined by HPLC-ICP-MS) post-oxidation with H_2_O_2_. Third row: total As (determined by ICP-MS). Individual values for each biological replicate are shown in Supporting Information Table S24.

**References**

(1) Viacava, K.; Qiao, J. T.; Janowczyk, A.; Poudel, S.; Jacquemin, N.; Meibom, K. L.; Shrestha, H. K.; Reid, M. C.; Hettich, R. L.; Bernier-Latmani, R. Meta-omics-aided isolation of an elusive anaerobic arsenic-methylating soil bacterium. *The ISME J.* **2022**, *16*, 1740–1749.
(2) Kerl, C. F.; Schindele, R. A.; Brüggenwirth, L.; Colina Blanco, A. E.; Rafferty, C.; Clemens, S.; Planer-Friedrich, B. Methylated thioarsenates and monothioarsenate differ in uptake, transformation, and contribution to total arsenic translocation in rice plants. *Environ. Sci. Technol.* **2019**, *53*, 5787−5796.
(3) Choi, K. H.; Gaynor, J. B.; White, K. G.; Lopez, C.; Bosio, C. M.; Karkhoff-Schweizer, R. R.; Schweizer, H. P. A Tn 7-based broad-range bacterial cloning and expression system. *Nat. Methods* **2005**, *2*, 443-448.
