## supporting methods SM1-SM7; supporting results SR1-SR4; supplemental figures S1-S18; supplemental tables S1-S3. for "Growth substrate limitation enhances anaerobic arsenic methylation by *Paraclostridium bifermentans* strain EML"

**TEXT**

**Supporting Methods:**

**Text SM1: Chemical stability of MMAs(III) (abiotic controls)**

All abiotic controls were conducted at 30°C in the dark without shaking. Three types of controls were prepared: (i) For fresh RCB abiotic controls, systematic dilutions of RCB (100%, 75%, 50%, or 25% RCB) were prepared in duplicate 120 mL serum bottles containing 50 mL medium following the anaerobic procedures described in the Materials and Methods section. MMAs(III) (3 μM) was added to each medium using sterile, N_2_-flushed syringes and needles. (ii) For spent RCB abiotic controls, a pre-culture of strain EML was grown in RCB anaerobically to mid-exponential growth phase, then the spent medium was prepared by filtering out the cells with a sterile Nalgene Rapid-Flow 0.1 μm PES membrane filter (Thermo Fisher) twice in an anoxic gas chamber (Coy Laboratory Products). Filtered spent medium (25 mL) was injected into pre-autoclaved duplicate serum bottles with a 100% N_2_ atmosphere, followed by the addition of 3 μM MMAs(III) to each medium replicate using sterile, N_2_-flushed syringes and needles. (iii) To assess the chemical reaction between MMAs(III) and sulfide, anaerobic sterile ultrapure water was prepared in duplicate serum bottles, amended with 3 μM MMAs(III) and then 6 μM sodium sulfide using sterile, N_2_-flushed syringes and needles.

For all cases, samples for aqueous As speciation were obtained by collecting 1 mL of solution with sterile, N_2_-flushed syringes and needles, filtered through 0.22 μm filters, and stored at -80°C after directly flash-freezing in liquid nitrogen. To probe whether methylated and thiolated As species were formed, As species were oxidized by adding 10% (v/v) H_2_O_2_ and storing overnight in a 1% HNO_3_ solution. As speciation was determined by HPLC-ICP-MS on the Agilent 8900 ICP-QQQ instrument (Agilent Technologies) using the previously described anion exchange protocol (1). Three As standards were prepared, including MMAs(III) and MMAs(V) (both commercially available), and MMMTAs(V) (that was synthesized).

**Text SM3: Validation of iAs(III)-resistance in *E. coli***

To investigate iAs(III) resistance in *E. coli* strain MG1655 and rule out the possible confounding effects on growth caused by the presence of iAs(III) in co-culture systems, 120 mL serum bottles were prepared with 50 mL anaerobic 100% RCB medium. Pre-cultures of WT *E. coli* and ArsP-expressing *E. coli* (henceforth ArsP *E. coli*) were grown in RCB anaerobically to mid-exponential growth phase. Both WT and ArsP *E. coli* (~0.5 mL) were inoculated separately into each RCB medium supplied with 10, 15, and 25 μM iAs(III) in triplicate using sterile N_2_-flushed syringes and needles. At selected time points and for each condition, triplicate bottles were sampled, and growth was measured as optical density at 600 nm (OD_600_) in a spectrophotometer. In addition, aqueous As species was also quantified at the beginning and the end of the incubation by HPLC-ICP-MS as described in the main text Materials and Methods section (Arsenic Speciation).

**Supporting Results:**

**Text SR1: iAs(III)-resistance in *E. coli***

Anaerobic growth of WT *E. coli* and ArsP *E. coli* was conducted in 100% RCB supplied with 10, 15, and 25 μM iAs(III). As illustrated in Fig. S3, both *E. coli* strains exhibited similar growth patterns under different iAs(III) concentrations, i.e., grew rapidly during 20 hours of incubation, then reached a plateau after 24 hours. This result demonstrates that the presence of iAs(III) did not affect the growth of WT *E. coli* and ArsP *E. coli* in RCB.

**FIGURES**

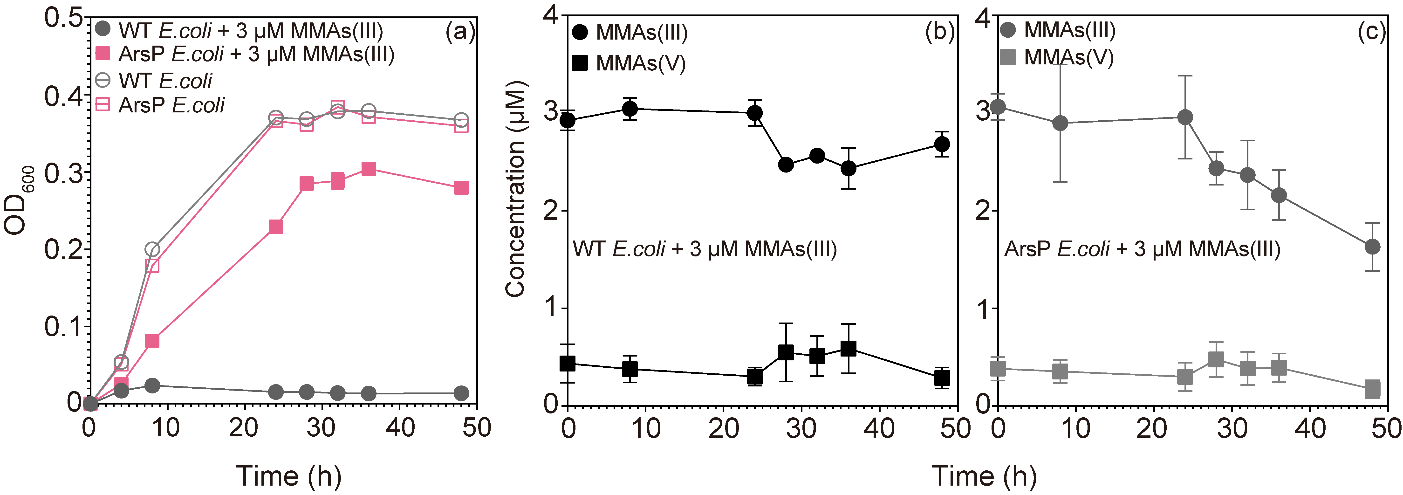

**FIG S1.** (a) Growth curves (OD_600_) of *Escherichia coli* K-12 wild-type strain MG1655 (WT *E. coli*) and engineered WT *E. coli* harboring an expressing MMAs(III)-resistance gene (*arsP*) (ArsP *E. coli*) in anoxic 1/4 tryptic soy broth (TSB) medium in the presence or absence of 3 μM MMAs(III). (b) and (c) Time-dependent stability of aqueous MMAs(III) in 1/4 TSB incubating with WT *E. coli* and ArsP *E. coli*. Data are shown as mean values with error bars. Individual values for each biological triplicate are included in Data Table S1.

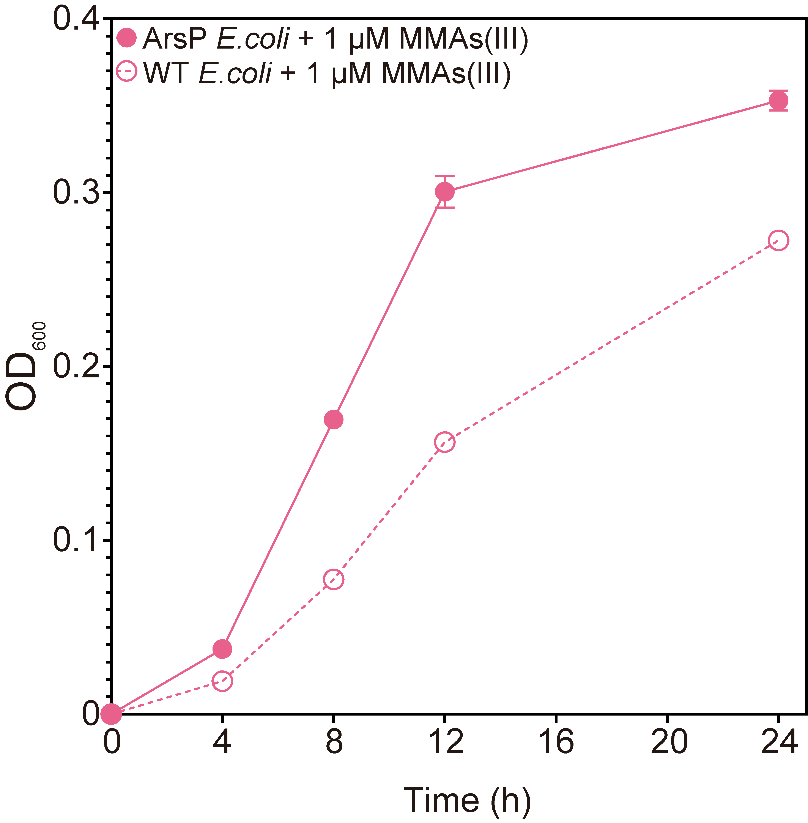

**FIG S2.** Growth curves (OD_600_) of WT *E. coli* and ArsP *E. coli* in anoxic 1/4 TSB in the presence or absence of 1 μM MMAs(III). Data are shown as mean values with error bars. Individual values for each biological triplicate are included in Data Table S2.

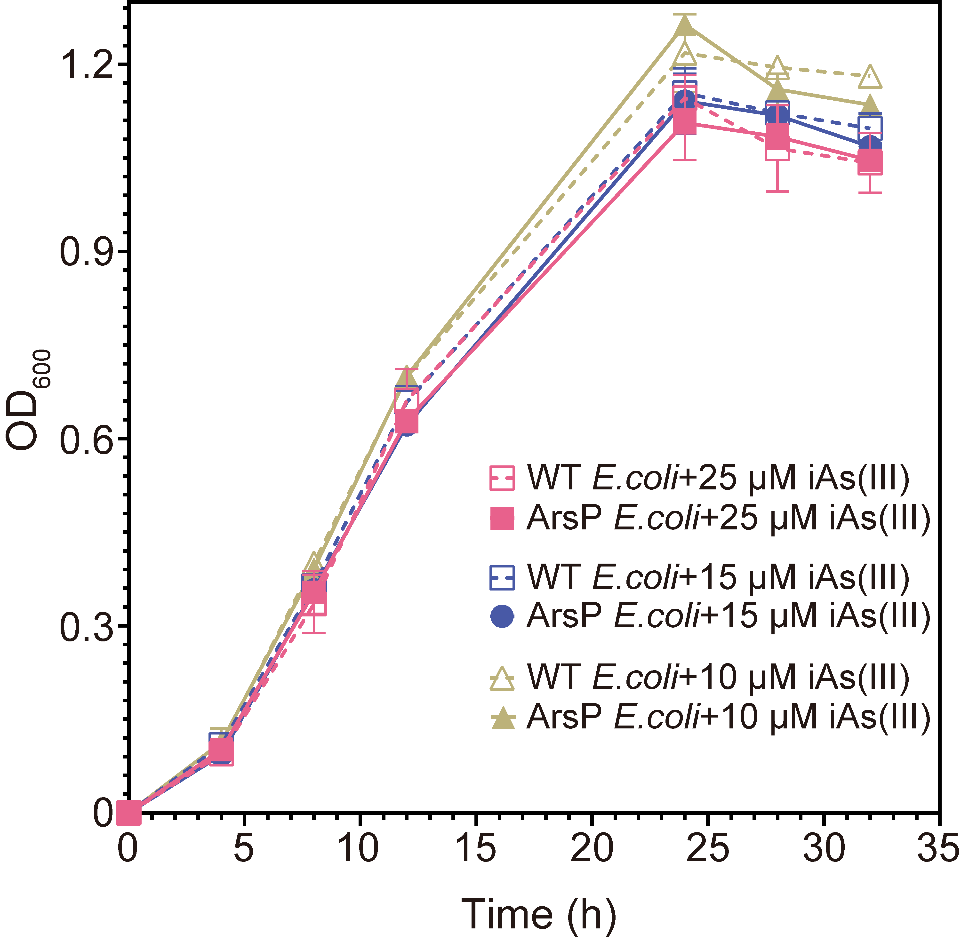

**FIG S3.** Growth curves (OD_600_) of WT *E. coli* and ArsP *E. coli* in anoxic RCB in the presence or absence of 10, 15, or 25 μM iAs(III). Data are shown as mean values with error bars. Individual values for each biological triplicate are included in Data Table S3.

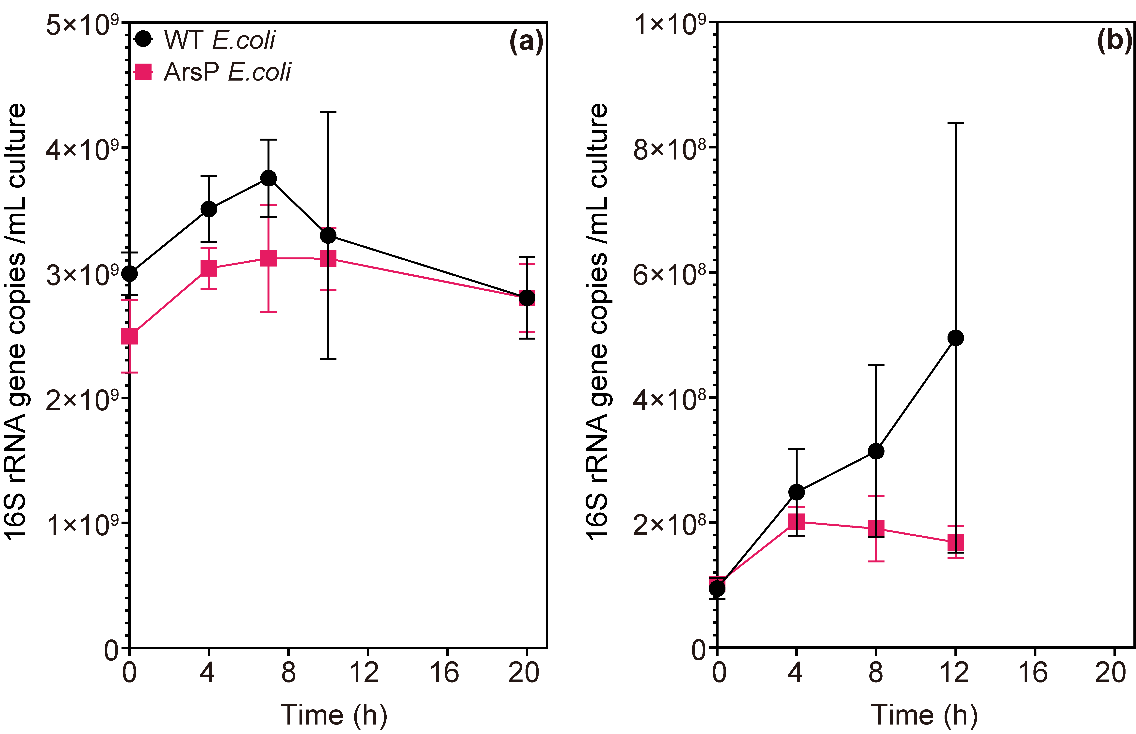

**FIG S4.** Growth curves (16S rRNA gene copy number) of WT *E. coli* and ArsP *E. coli* in anaerobic co-culture with *Paraclostridium bifermentans* strain EML in anoxic RCB supplied with 25 μM iAs(III). The result corresponds to the first and second attempts to optimize inoculation ratio between the co-culture strains in Supplementary Information Text Results SR3. *E. coli* (either strain) was added only after 6 hours of strain EML growth alone. The two conditions are: (a) 0.5 mL strain EML + 0.3 mL *E. coli*, (b) 0.5 mL strain EML + 0.03 mL *E. coli*. Data are shown as mean values with error bars. Individual values for each biological triplicate are included in Data Table S4.

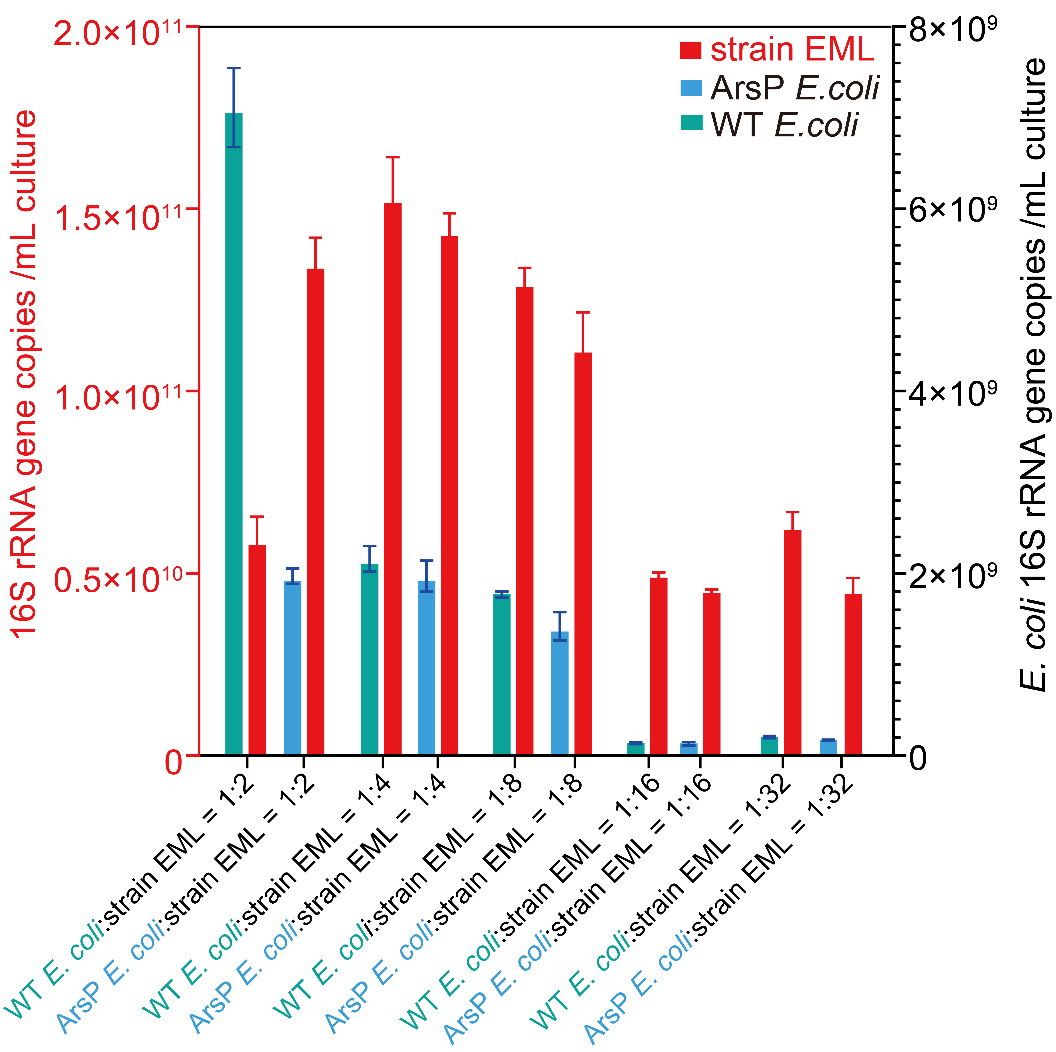

**FIG S5.** Growth (16S rRNA gene copy number) of the co-culture members in anoxic RCB supplied with 25 μM iAs(III) after 12 hours of incubation. The result corresponds to the third attempt to optimize inoculation ratio between the co-culture strains in Supplementary Information Text Results SR3. *Paraclostridium bifermentans* strain EML (red) was co-culture with either WT *E. coli* (green) or ArsP *E. coli* (blue) anaerobically at various inoculation ratios of *E. coli*: strain EML = 1:2, 1:4, 1:8, 1:16, and 1:32 (v/v). Data are shown as mean values with error bars. Individual values for each biological triplicate are included in Data Table S5.

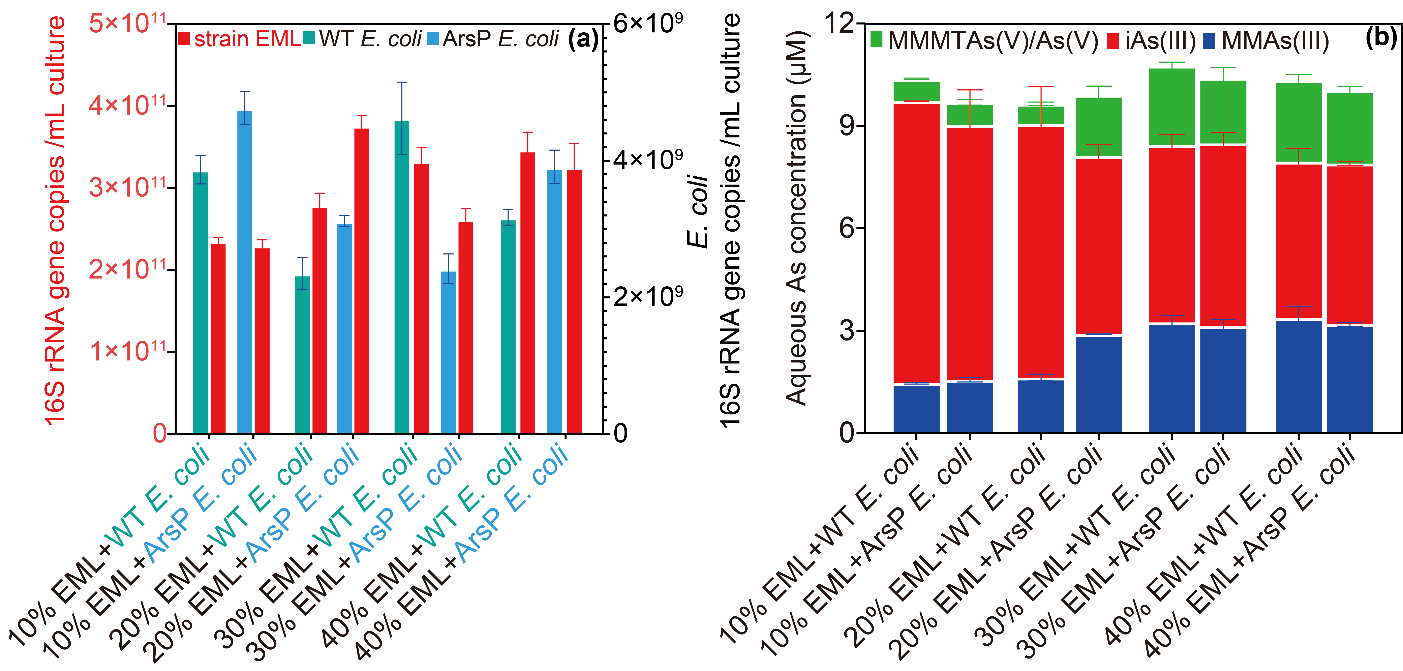

**FIG S6.** (a) Growth (16S rRNA gene copy number) of the co-culture members in anoxic RCB supplied with 25 μM iAs(III) after 12 hours of incubation. The result corresponds to the fourth attempt to optimize inoculation ratio between the co-culture strains in Supplementary Information Text Results SR3. *Paraclostridium bifermentans* strain EML (red) (various volumes of cell culture) was co-cultured with either WT *E. coli* (green) or ArsP *E. coli* (blue) (a 50 μL culture at exponential phase). The amount of strain EML in RCB medium was systematically varied from the cell pellet of a 5 mL culture (10%), 10 mL culture (20%), 15 mL culture (20%), or 20 mL culture (40%). (b) Aqueous As speciation in the above-described co-culture systems. Data are shown as mean values with error bars. Individual values for each biological triplicate are included in Data Table S6.

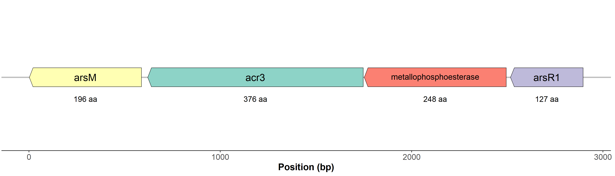

**FIG S7.** Genetic organization of the plasmid-encoded *ars* operon from *Paraclostridium bifermentans* strain EML. Arrows represent open reading frames and orientation of transcription.

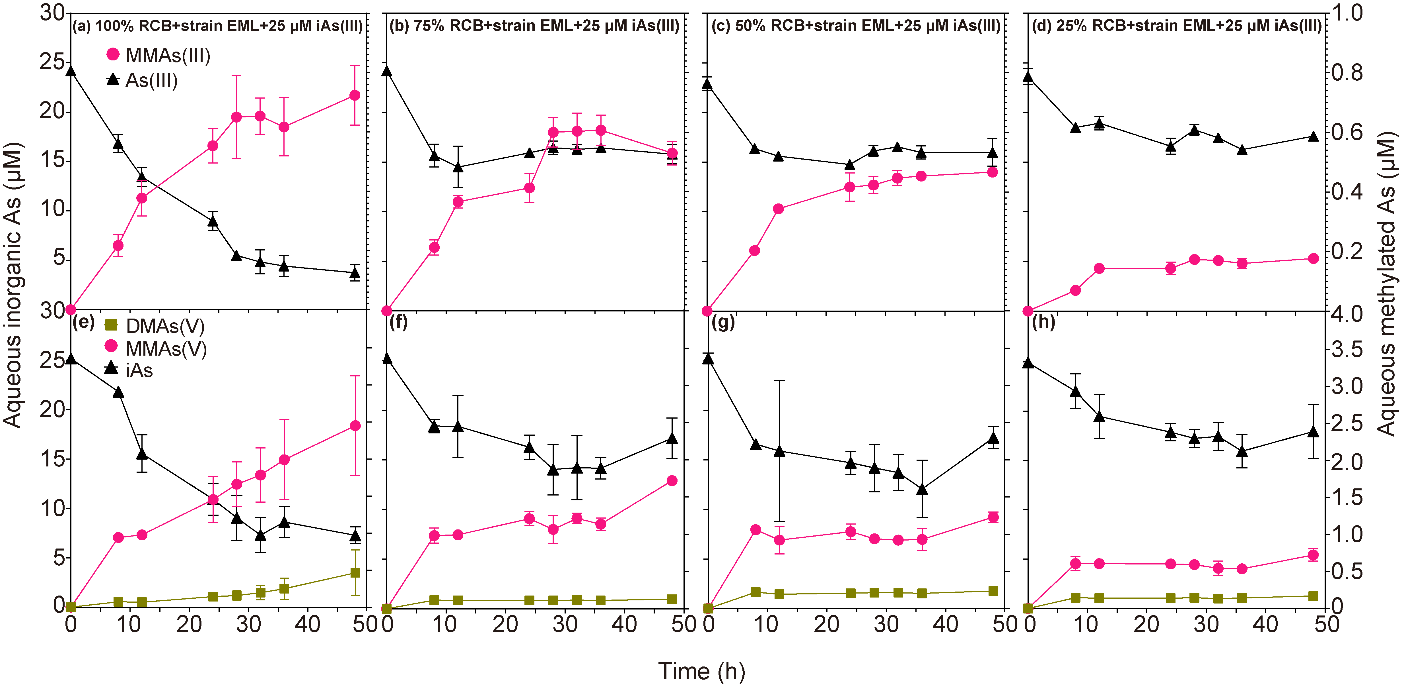

**FIG S8.** Time-dependent concentrations of aqueous As species produced by *Paraclostridium bifermentans* strain EML in anaerobic RCB dilutions (100%, 75%, 50%, or 25% RCB) in the presence of 25 μM iAs(III). (a)-(d), aqueous As species determined by HPLC-ICP-MS. (e)-(f), aqueous As species determined by HPLC-ICP-MS post-oxidation with H_2_O_2_. Individual values for each biological replicate are listed in Data Table S8.

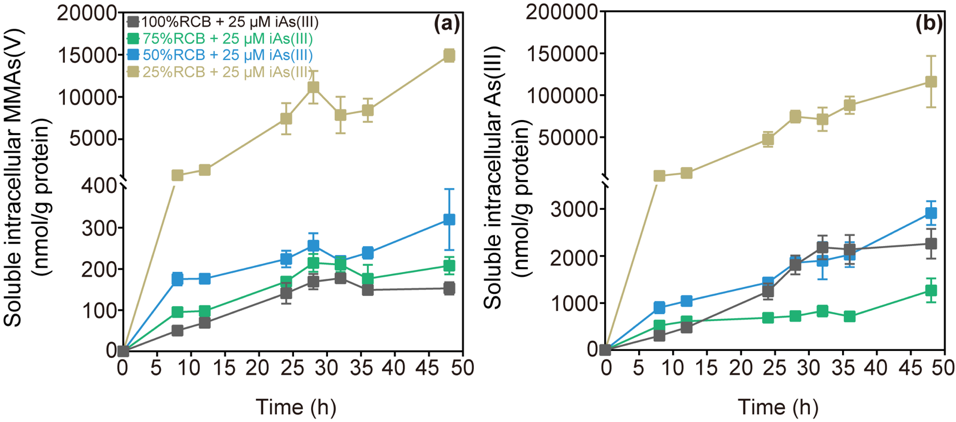

**FIG S9.** Time-dependent concentration of normalized soluble intracellular MMAs(V) (a) and intracellular iAs(III) (b) in anoxic RCB dilutions (100%, 75%, 50%, or 25% RCB) inoculated with *Paraclostridium bifermentans* strain EML and amended with 25 μM iAs(III). Individual values for each biological replicate can be found in Data Tables S9 and S21.

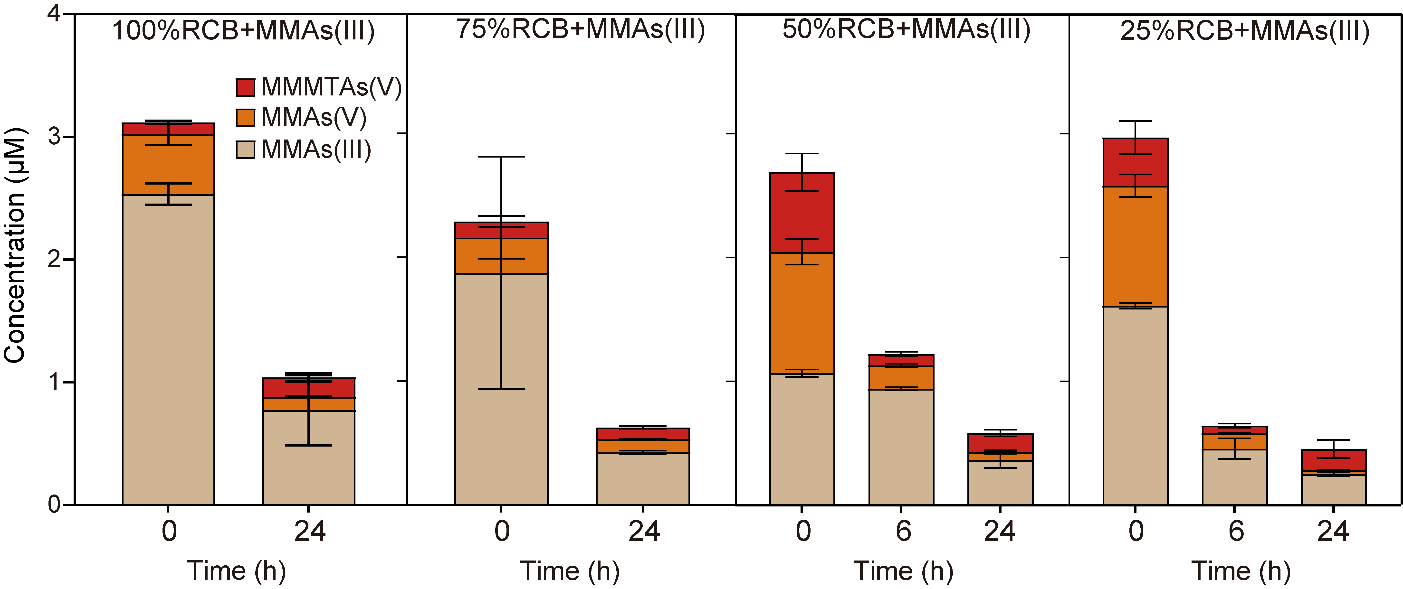

**FIG S10.** Time-dependent stability of aqueous MMAs(III) (3 μM) in anoxic RCB dilutions (100%, 75%, 50%, or 25% RCB). Individual values for each replicate are listed in Data Table S10.

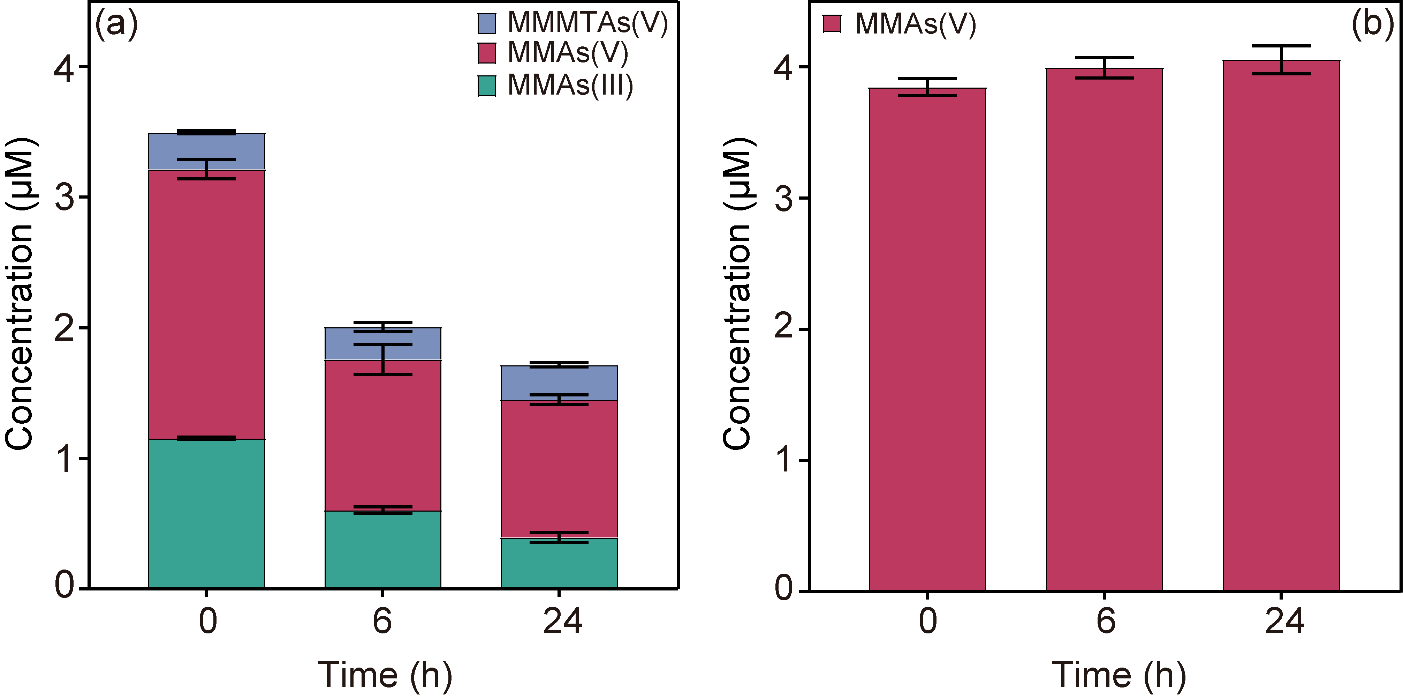

**FIG S11.** Time-dependent stability of aqueous MMAs(III) (3 μM) in anoxic spent RCB (100% RCB). (a) As species without oxidation, and (b) As species post-oxidation. Individual values for each replicate are listed in Data Table S11.

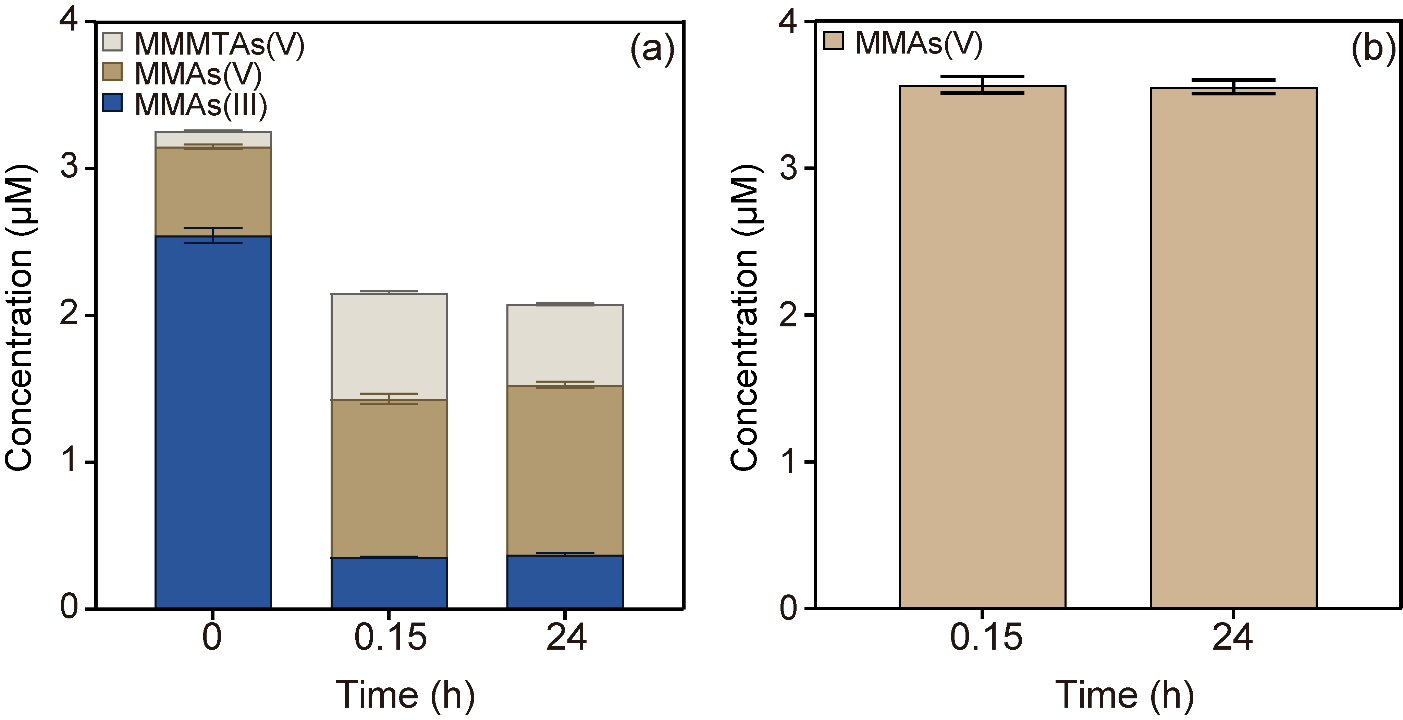

**FIG S12.** Time-dependent chemical reaction between MMAs(III) and sulfide in anoxic water solution. (a) As species without oxidation, and (b) As species with post-oxidation. Individual values for each replicate are listed in Data Table S12.

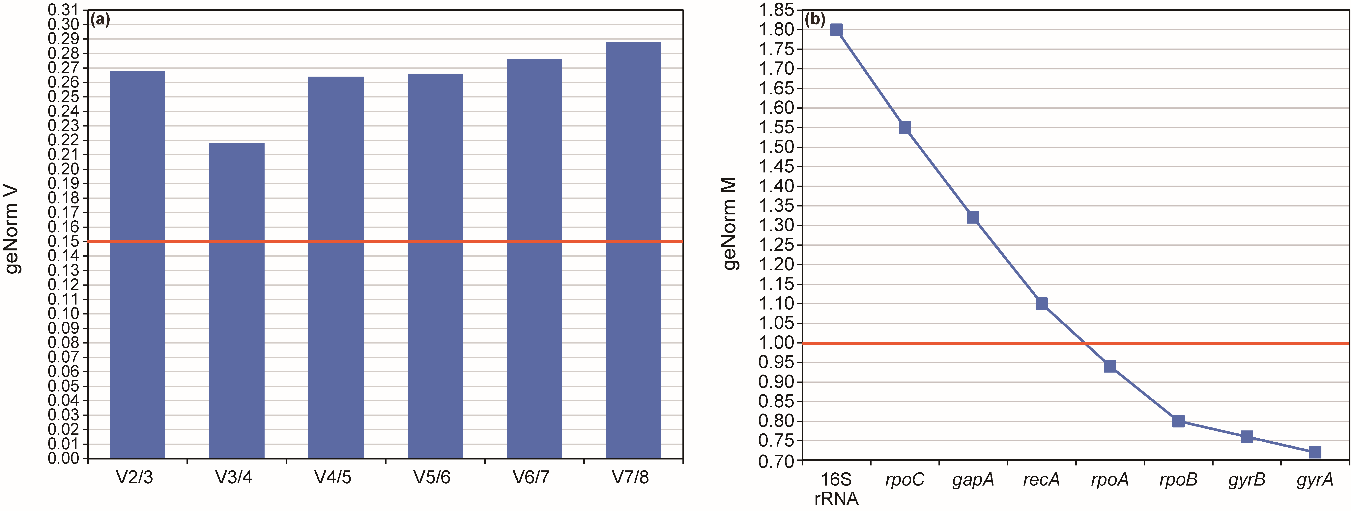

**FIG S13.** (a) The determination of minimum number of reference genes based on the average pairwise variation (V) of normalization factors and (b) the calculated expression stability of the candidate reference genes using the geNorm program within qBase plus software. The red line indicates the minimum thresholds (geNorm V (0.15) and geNorm M (1.0)) required for reference genes. None of the genes fulfill this requirement.

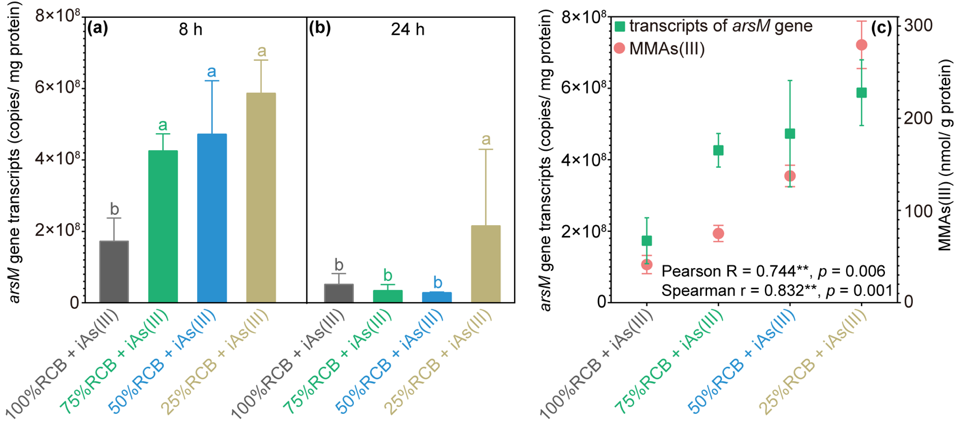

**FIG S14.** Transcript numbers of *arsM* gene of *Paraclostridium bifermentans* strain EML in anoxic dilutions (100%, 75%, 50%, and 25%) of RCB in the presence of 25 μM iAs(III) at 8 (a) and 24 (b) hours of incubation. Correlation analysis of *arsM* transcripts and MMAs(III) concentration at 8 hours of incubation (c). Different letters showed significant difference between RCB dilutions at *P* < 0.05. Individual values for each biological replicate are shown in Data Table S14. This figure is different from Fig. 3 in that the absolute quantification relies on using the same amount total RNA for reverse transcription regardless of biomass amount used for the RNA extraction.

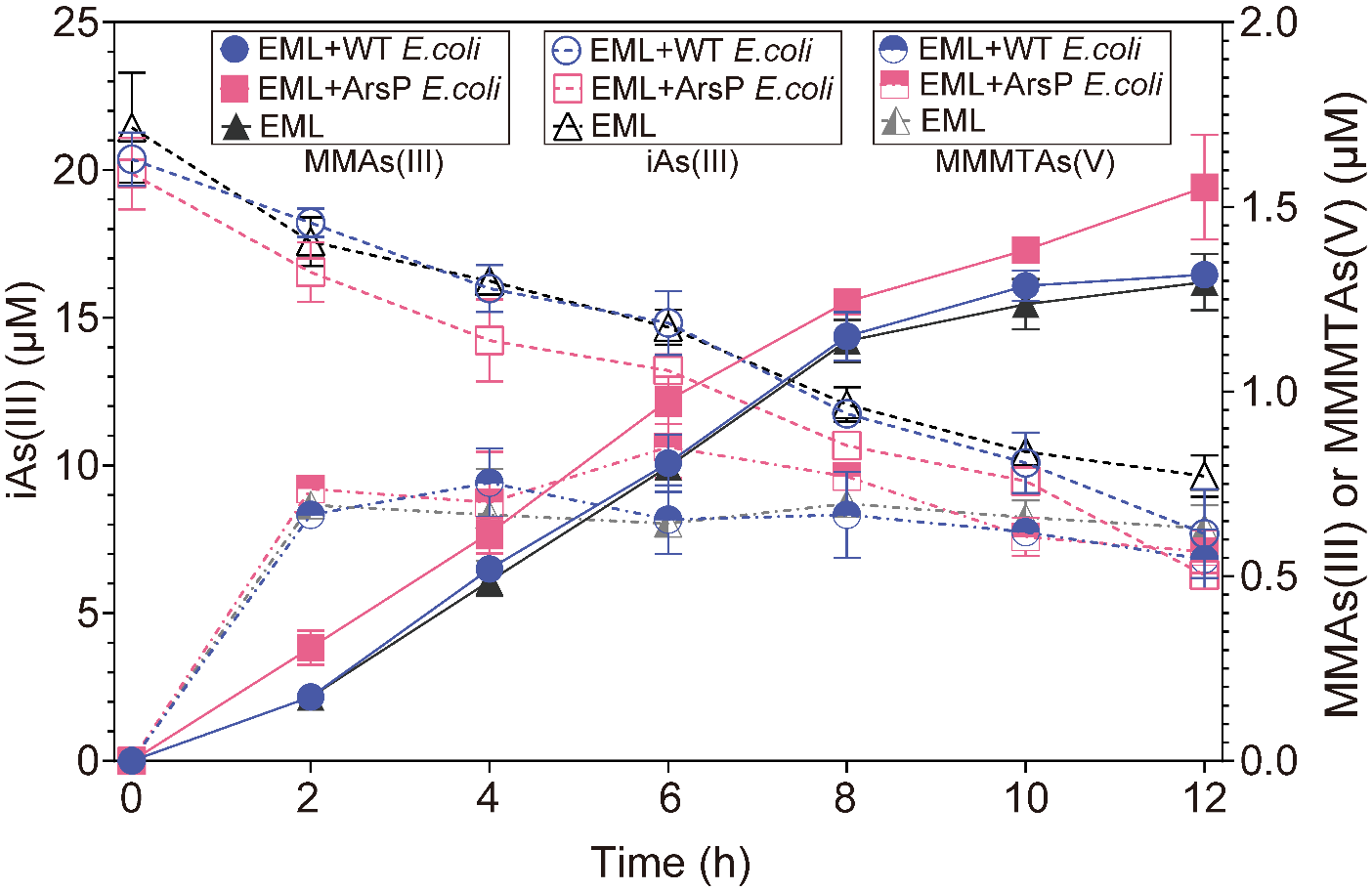

**FIG S15.** Time-dependent concentrations of aqueous As species in anaerobic co-culture *Paraclostridium bifermentans* strain EML with either WT *E. coli* or ArsP *E. coli* in anoxic RCB with 25 μM iAs(III). Individual values for each biological replicate can be found in Data Table S15.

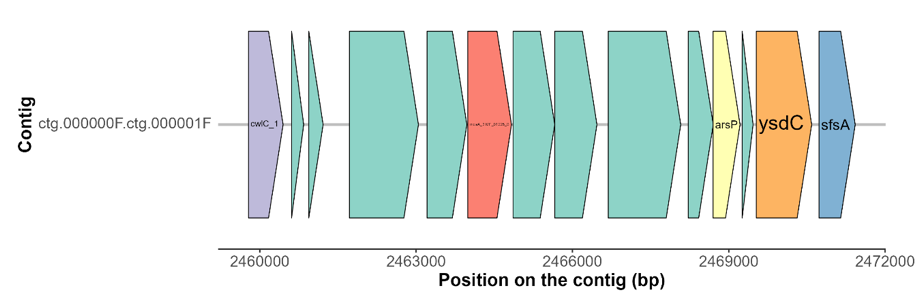

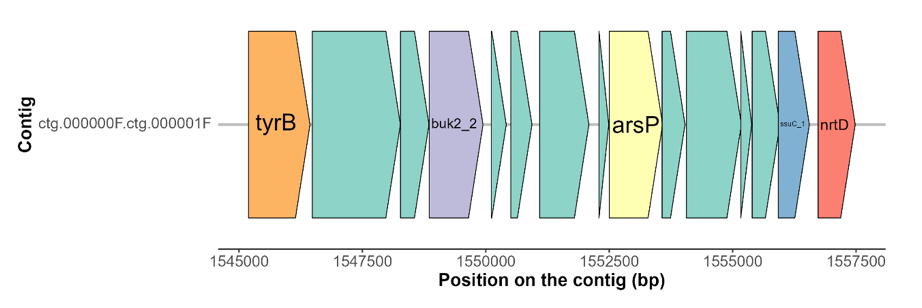

**FIG S16.** The two chromosomally encoded *arsP* genes identified from *Paraclostridium bifermentans* strain EML. Arrow represents open reading frames and orientation of transcription. *ysdC*: Putative aminopeptidase YsdC, *sfsA*: sugar fermentation stimulation protein A, *ssuC*: aliphatic sulfonate ABC transporter membrane subunit, *nrtD*: nitrate/nitrite transport system ATP-binding protein. No other identifiable arsenic-related genes were identified in the vicinity of these two gene. Genes with unknown or uncertain functions are represented as green color.

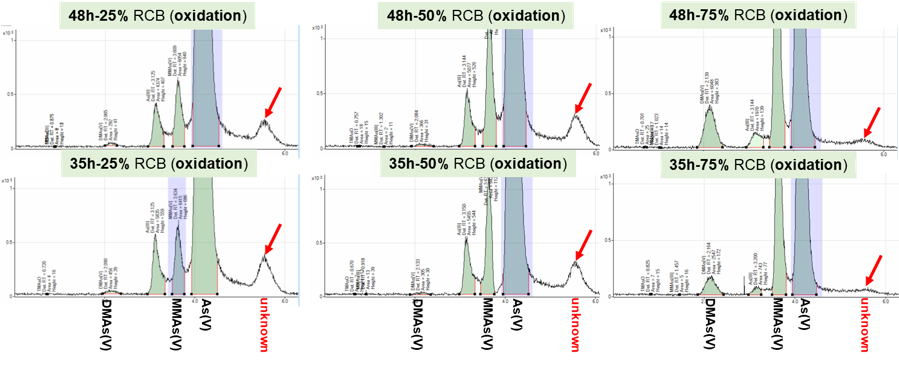

**FIG S17.** The unknown As peak in anaerobic RCB dilutions (75%, 50%, or 25% RCB) inoculated with *Paraclostridium bifermentans* strain EML and 25 μM iAs(III) obtained in oxidized samples at 35 and 48 hours of incubation. The unknown peak was also detected in 100% RCB but was very small compared with other RCB dilutions (not shown).

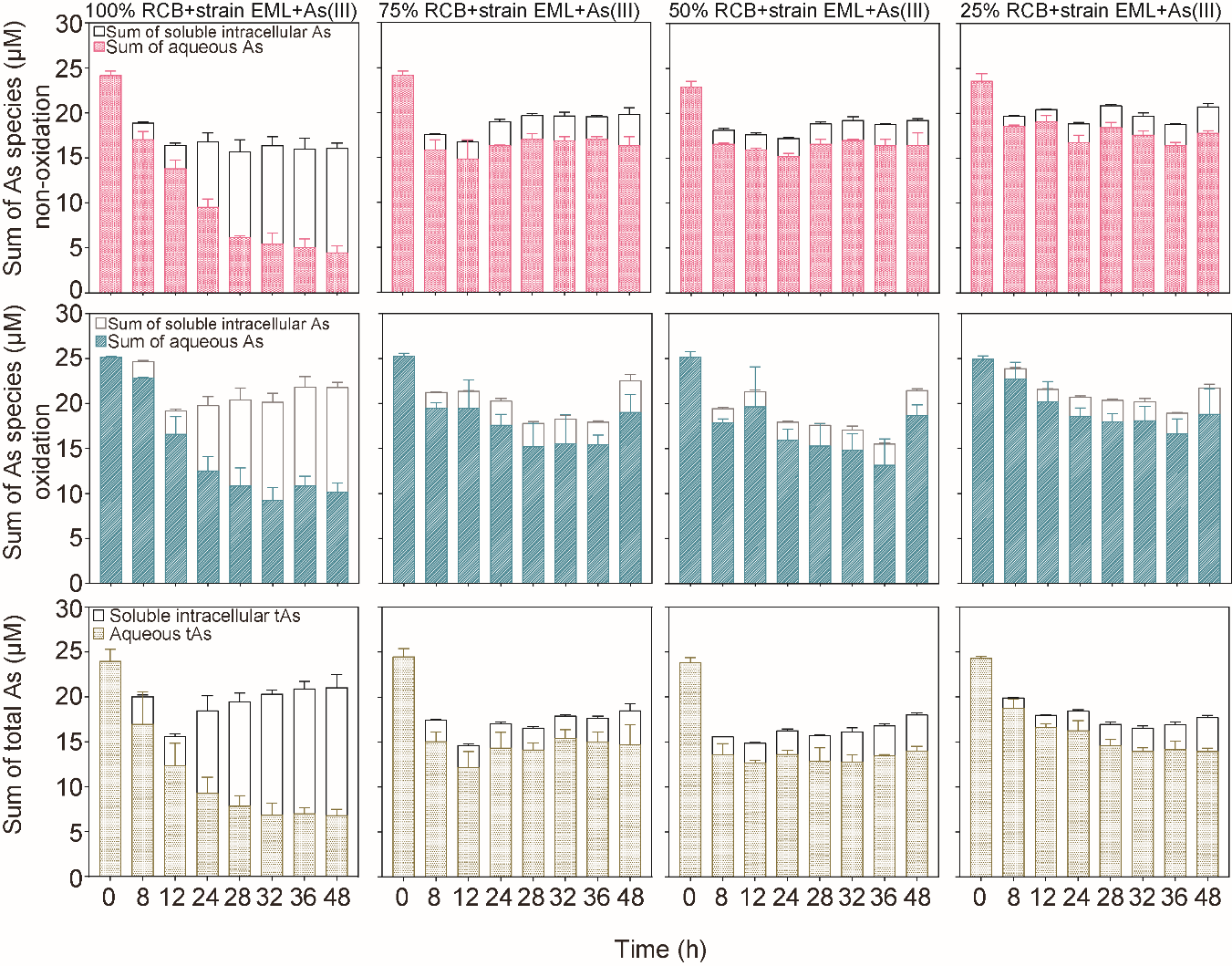

**FIG S18.** Time-dependent comparison of As mass balance in anoxic dilutions (100%, 75%, 50%, and 25%) of RCB inoculated with *Paraclostridium bifermentans* strain EML and 25 μM iAs(III). First row: sum of aqueous As species and soluble intracellular As species (determined by HPLC-ICP-MS). Second row: sum of aqueous As species and soluble intracellular As species (determined by HPLC-ICP-MS) post-oxidation with H_2_O_2_. Third row: total As (determined by ICP-MS). Individual values for each biological replicate are shown in Data Table S18.

**TABLES**

**Table S1.** The experimental treatments used in the variable growth substrate experiments.

| Treatment | Replicate | Milli-Q water (mL) | RCB (mL) | iAs(III) (μM) | Strain EML (mL) | Headspace |
| --- | --- | --- | --- | --- | --- | --- |
| 100% RCB+*P. bifermentans* strain EML+iAs(III) | 1 | 0 | 50 | 25 | 0.5 mL of pre-culture (OD_600_ = 2.1) | 100% N_2_ |
|  | 2 |  |  |  |  |  |
|  | 3 |  |  |  |  |  |
| 100% RCB+*P. bifermentans* strain EML | 4 |  |  | 0 |  |  |
|  | 5 |  |  |  |  |  |
|  | 6 |  |  |  |  |  |
| 75% RCB+*P. bifermentans* strain EML+iAs(III) | 7 | 12.5 | 37.5 | 25 |  |  |
|  | 8 |  |  |  |  |  |
|  | 9 |  |  |  |  |  |
| 75% RCB+*P. bifermentans* strain EML | 10 |  |  | 0 |  |  |
|  | 11 |  |  |  |  |  |
|  | 12 |  |  |  |  |  |
| 50% RCB+*P. bifermentans* strain EML+iAs(III) | 13 | 25 | 25 | 25 |  |  |
|  | 14 |  |  |  |  |  |
|  | 15 |  |  |  |  |  |
| 50% RCB+*P. bifermentans* strain EML | 16 |  |  | 0 |  |  |
|  | 17 |  |  |  |  |  |
|  | 18 |  |  |  |  |  |
| 25% RCB+*P. bifermentans* strain EML+iAs(III) | 19 | 37.5 | 12.5 | 25 |  |  |
|  | 20 |  |  |  |  |  |
|  | 21 |  |  |  |  |  |
| 25% RCB+*P. bifermentans* strain EML | 22 |  |  | 0 |  |  |
|  | 23 |  |  |  |  |  |
|  | 24 |  |  |  |  |  |

**Table S2**. Selected reference genes for gene expression by real-time RT-qPCR.

| Target gene | Function | Primer set | Primer sequence | Thermal profile for real-time PCR | Amplicon size (bp) |
| --- | --- | --- | --- | --- | --- |
| *rpoA* | alpha subunit of RNA polymerase | rpoA-F | ACTGTGCATGGTCCTTGTGCTT | 95°C 5 min, 50 cycles of 95°C 5 s, 60°C 20 s, and 72°C 20 s | ~135 |
|  |  | rpoA-R | TCATGAATTCACAACAGTGTCTGGT |  |  |
| *rpoB* | beta subunit of RNA polymerase | rpoB-F | CGCACGGTTGGCGTCATCATTC |  | ~195 |
|  |  | rpoB-R | AGAGCCCAAGCAAATGAGCCTT |  |  |
| *rpoC* | beta subunit of RNA polymerase | rpoC-F | CGGCAAGACCCTTTCTAGCTCCA |  | ~125 |
|  |  | rpoC-R | GCGTGGTCTGATGGCCAATGCA |  |  |
| *gyrA* | DNA gyrase A subunit | gyrA-F | ACGTCCCCAACAATACGGGCAG |  | ~132 |
|  |  | gyrA-R | AGCTGGACGTGCCCTACCAGAT |  |  |
| *gyrB* | DNA gyrase B subunit | gyrB-F | CCTCCGGCAGAATCCCCTTCGA |  | ~81 |
|  |  | gyrB-R | ACCTGGAAAATTAGCTGACTGTGC |  |  |
| *recA* | DNA recombination/repair protein RecA | recA-F | AGAGCACGACCACCTGTTGTAGT |  | ~144 |
|  |  | recA-R | AGCAAGACTTATGTCGCAAGCT |  |  |
| *gapA* | D-glyceraldehyde-3-phosphate dehydrogenase | gapA-F | AGGACTTGGAACCCTTACTGCA |  | ~94 |
|  |  | gapA-R | ACCAACAACAACAGGTGCAGCT |  |  |
| 16S rRNA | small subunit ribosomal RNA molecules of ribosomes | 16S rRNA-F | AGCTGACGACAACCATGCAC |  | ~82 |
|  |  | 16S rRNA-R | TTGACATCCCACTGACCTCTCC |  |  |

**Table S3**. The concentration of aqueous As species for each replicate in anaerobic dilutions of sterile RCB (100%, 75%, 50%, or 25% RCB) amended with 25 μM iAs(III).

| 100% RCB + strain EML + 25 μM iAs(III)-no oxidation [μM] | | | | | | | | | |
| --- | --- | --- | --- | --- | --- | --- | --- | --- | --- |
| Time/ h | iAs(III) | | | iAs(V) | | | iAs | | |
| 0 | 22.30 | 21.13 | 21.72 | 1.22 | 2.20 | 2.08 | 23.52 | 23.33 | 23.80 |
| 48 | 21.37 | 22.28 | 22.87 | 1.17 | 1.27 | 1.87 | 22.54 | 23.55 | 24.74 |
| 75% RCB + strain EML + 25 μM iAs(III)-no oxidation [μM] | | | | | | | | | |
| Time/ h | iAs(III) | | | iAs(V) | | | iAs | | |
| 0 | 21.14 | 20.50 | 21.17 | 1.37 | 1.86 | 1.29 | 22.51 | 22.36 | 22.46 |
| 48 | 22.98 | 22.06 | 21.55 | 1.92 | 1.18 | 1.16 | 24.90 | 23.24 | 22.71 |
| 50% RCB + strain EML + 25 μM iAs(III)-no oxidation [μM] | | | | | | | | | |
| Time/ h | iAs(III) | | | iAs(V) | | | iAs | | |
| 0 | 20.43 | 21.79 | 21.75 | 1.42 | 1.79 | 1.49 | 21.85 | 23.58 | 23.24 |
| 48 | 21.73 | 22.95 | 22.72 | 1.24 | 1.41 | 1.93 | 22.97 | 24.36 | 24.65 |
| 25% RCB + strain EML + 25 μM iAs(III)-no oxidation [μM] | | | | | | | | | |
| Time/ h | iAs(III) | | | iAs(V) | | | iAs | | |
| 0 | 20.51 | 22.07 | 21.44 | 1.90 | 1.43 | 1.48 | 22.41 | 23.50 | 22.92 |
| 48 | 21.15 | 22.73 | 21.41 | 1.01 | 1.30 | 1.80 | 22.16 | 24.03 | 23.21 |

**References**

1. Viacava K, Qiao JT, Janowczyk A, Poudel S, Jacquemin N, Meibom KL, Shrestha HK, Reid MC, Hettich RL, Bernier-Latmani R. (2022) Meta-omics-aided isolation of an elusive anaerobic arsenic-methylating soil bacterium. *The ISME Journal*, ***16***, 1740–1749.
2. Kerl CF, Schindele RA, Brüggenwirth L, Colina Blanco AE, Rafferty C, Clemens S, Planer-Friedrich B. (2019) Methylated thioarsenates and monothioarsenate differ in uptake, transformation, and contribution to total arsenic translocation in rice plants. *Environmental Science & Technology*, ***53***, 5787−5796.
3. Choi KH, Gaynor JB, White KG, Lopez C, Bosio CM, Karkhoff-Schweizer RR, Schweizer HP. (2005) A Tn 7-based broad-range bacterial cloning and expression system. *Nature Methods*, ***2***, 443-448.
